# *De novo* design of site-specific protein interactions with learned surface fingerprints

**DOI:** 10.1101/2022.06.16.496402

**Authors:** Pablo Gainza, Sarah Wehrle, Alexandra Van Hall-Beauvais, Anthony Marchand, Andreas Scheck, Zander Harteveld, Stephen Buckley, Dongchun Ni, Shuguang Tan, Freyr Sverrisson, Casper Goverde, Priscilla Turelli, Charlène Raclot, Alexandra Teslenko, Martin Pacesa, Stéphane Rosset, Sandrine Georgeon, Jane Marsden, Aaron Petruzzella, Kefang Liu, Zepeng Xu, Yan Chai, Pu Han, George F. Gao, Elisa Oricchio, Beat Fierz, Didier Trono, Henning Stahlberg, Michael Bronstein, Bruno E. Correia

## Abstract

Physical interactions between proteins are essential for most biological processes governing life. However, the molecular determinants of such interactions have been challenging to understand, even as genomic, proteomic, and structural data grows. This knowledge gap has been a major obstacle for the comprehensive understanding of cellular protein-protein interaction (PPI) networks and for the *de novo* design of protein binders that are crucial for synthetic biology and translational applications. We exploit a geometric deep learning framework operating on protein surfaces that generates fingerprints to describe geometric and chemical features critical to drive PPIs. We hypothesized these fingerprints capture the key aspects of molecular recognition that represent a new paradigm in the computational design of novel protein interactions. As a proof-of-principle, we computationally designed several *de novo* protein binders to engage four protein targets: SARS-CoV-2 spike, PD-1, PD-L1, and CTLA-4. Several designs were experimentally optimized while others were purely generated *in silico*, reaching nanomolar affinity with structural and mutational characterization showing highly accurate predictions. Overall, our surface-centric approach captures the physical and chemical determinants of molecular recognition, enabling a novel approach for the *de novo* design of protein interactions and, more broadly, of artificial proteins with function.

## Introduction

Designing novel protein-protein interactions (PPIs) remains a fundamental challenge in computational protein design, with broad basic and translational applications in biology. The challenge consists of generating amino acid sequences that engage a target site and form a quaternary complex with a given protein. This represents a stringent test of our understanding of the physicochemical determinants that drive biomolecular interactions^1^. Robust computational methods to design *de novo* PPIs could be used to rapidly engineer protein-based therapeutics such as antibodies and protein inhibitors or vaccines, among others, and therefore are of major interest for biomedical and translational applications^2–8^. Despite recent advances in rational PPI design^2, 6, 8^ and prediction^9^, designing novel protein binders against specific targets is very challenging, particularly when no structural elements from preexisting binders are known. Current state-of-the-art methods for *de novo* PPI design^2, 6, 10, 11^, such as hotspot-centric approaches^6^ and rotamer information fields^2, 8^, rely on placing disembodied residues on the target interface and then optimizing their presentation on a protein scaffold. Intrinsic limitations of these approaches relate to the very weak energetic signatures provided by scoring functions to single-side chain placements, which is compounded in flat interfaces that lack deep pockets. These methods also face the challenge of finding compatible protein scaffolds to precisely display the generated constellations of residues. To circumvent these limitations, new approaches are needed to design *de novo* binders to various surface types and protein sites.

A long-standing model of molecular recognition postulates that PPIs form between protein molecular surfaces with chemical and geometric complementarity^12, 13^. The complementarity features arise as a consequence of the energetic contributions that are critical to stabilize PPIs, including van der Waals interactions (geometric complementarity), hydrophobic effect, and electrostatics interactions (chemical complementarity)^12^. At the structural level, most protein interfaces contain surface regions that become inaccessible to solvent upon complex formation, which we refer to as *buried* or *core interface*, as well as patches that are involved in the interface but remain solvent-exposed, which we refer to as the *interface rim*. Residues within the buried areas tend to be much less tolerant to mutations^14, 15^ and have a large energetic contribution towards the PPI formation, often referred to as hotspots. Rim regions are generally more polar and tolerant to mutations, giving also important contributions to affinity and, more notably, specificity^14, 16^. Guided by these general principles of molecular recognition, we introduce a novel protein design approach based on the critical importance of the fully buried patches of the interface to drive protein interactions. We implemented these design principles by exploiting surface fingerprints learned from interacting protein surfaces which capture features that are determinant for molecular recognition. Our novel approach allows for ultra-fast and accurate prediction of privileged sites for PPI design, and reduces the complexity for hotspot search and grafting. We leveraged this design workflow to successfully engineer and characterize binders against four therapeutic targets of interest, namely SARS-CoV-2 spike, PD-1, PD-L1, and CTLA-4.

### Design of *de novo* PPIs using learned surface fingerprints

In previous work, we introduced a geometric deep learning framework, MaSIF (Molecular Surface Interaction Fingerprinting), to generate surface fingerprints from the geometric and chemical features of molecular surfaces and learn patterns that determine the propensity of protein interactions^17^. Within this framework we developed the MaSIF-site tool to predict areas with propensity to form PPIs on the surface of proteins. MaSIF-site receives as input a protein decomposed into patches and outputs a per-vertex regression score on the propensity of each surface point to become a buried site within a PPI. We also developed MaSIF-search, another tool to evaluate surface complementarity between binding partners. MaSIF-search was designed as a Siamese neural network architecture^18^ trained to produce similar fingerprints for the target patch vs. the binder patch, and dissimilar fingerprints for the target patch vs. the random patch. As MaSIF tools had robust performance in PPI-related *prediction* tasks, we hypothesized that we could leverage them to *design* novel PPIs by targeting sites only using structural information from the target protein. To address the *de novo* PPI design problem we devised a three-stage computational approach depicted in Fig. 1: I) prediction of target buried interface sites with high binding propensity using MaSIF-site (Fig. 1a); II) surface fingerprint-based search for complementary structural motifs (*binding seeds*) that display the required features to engage the target site, a protocol we refer to as MaSIF-seed (Fig. 1a,b); III) binding seed transplantation to protein scaffolds to confer stability and additional contacts on the designed interface (Fig. 1c) using established transplantation techniques^19^.

**Figure 1:**
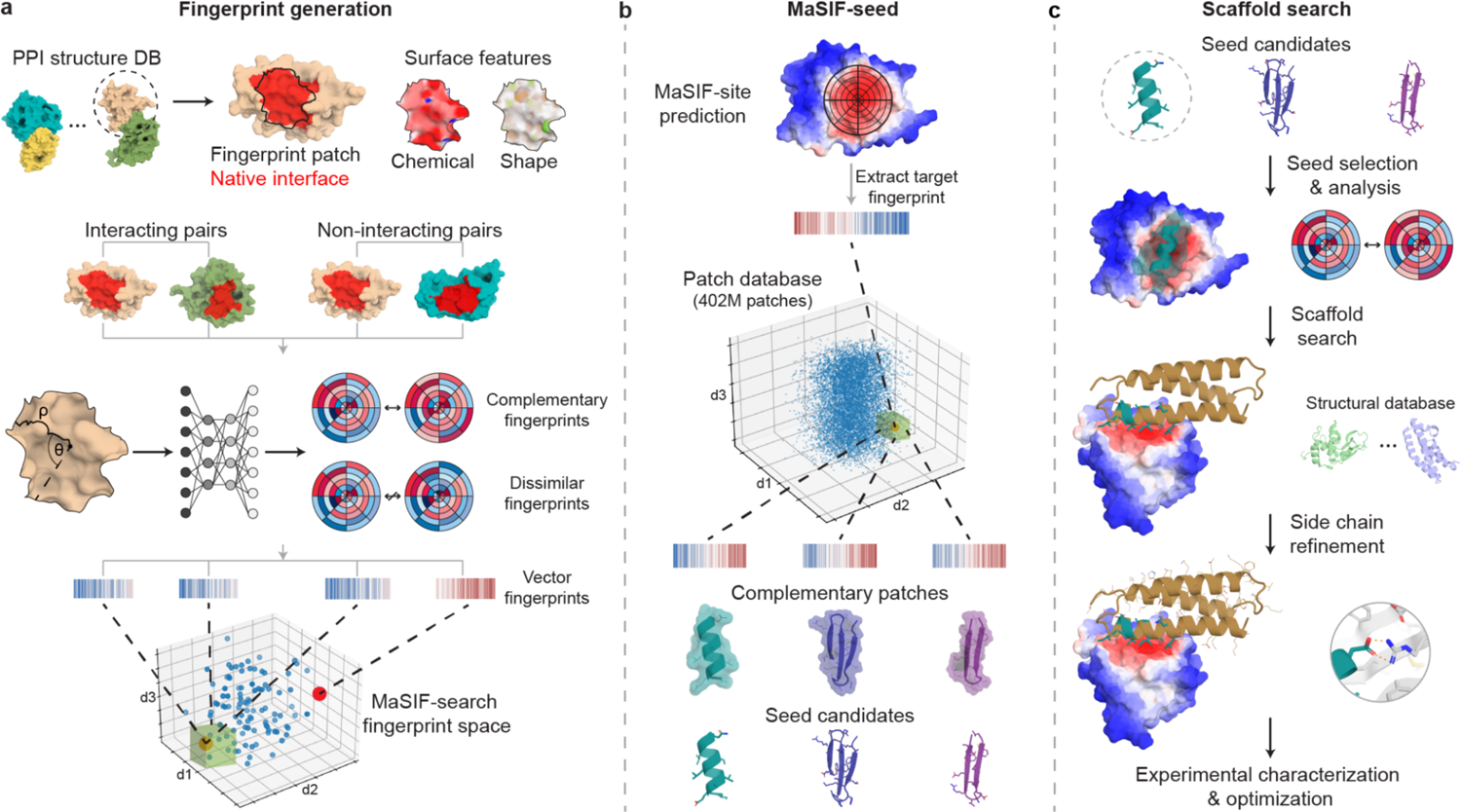
Surface-centric design of *de novo* site-specific protein binders. **a**, Protein binding sites are spatially embedded as vector fingerprints. Protein surfaces are decomposed into overlapping radial patches and a neural network trained on native interacting protein pairs learns to embed the fingerprints such that complementary fingerprints are placed in a similar region of space. We show an illustration for a subsample of the fingerprints projected in a space reduced to three dimensions. The green box highlights a region of complementary fingerprints. **b**, MaSIF-seed - a new method to identify new binding seeds. A target patch is identified by MaSIF-site based on the propensity to form buried interfaces. Using MaSIF-seed, fingerprint complementarity is evaluated between the target patch and all fingerprints in a large database (∼402 M patches), subsequently the pairs of fingerprints are ranked. The top patches are aligned and rescored to allow for a more precise evaluation of the seed candidates. **c**, Scaffold search, seed grafting and interface redesign. The selected seeds are transferred to protein scaffolds and the rest of the interface is redesigned using Rosetta. The top designs are selected and tested experimentally.

The new MaSIF-seed protocol tackles the problem of identifying binding seeds that can mediate productive binding interactions (Fig. 1, Supplementary Fig. S1). This task stands as a remarkable challenge in protein design due to the vast space of structural possibilities to explore, as well as the required precision given that subtle atomic-level changes, such as misplaced methyl groups^19, 20^, uncoordinated water molecules in the interface, or incompatible charges, are sufficient to disrupt PPIs^21^.

In MaSIF-seed, protein molecular surfaces are decomposed into overlapping radial patches with a 12 Å radius, capturing on average nearly 400 Å^2^ of surface area, consistent with the buried surface areas observed in native interfaces (Supplementary Fig. S2). For each point within the patch, we compute chemical and geometric features, as well as a local geodesic polar coordinate system to locate points within the patch relative to each other. A neural network is then trained to output vector fingerprint descriptors that are complementary between patches of interacting protein pairs and dissimilar between non-interacting pairs^17^ (Fig. 1a, Supplementary Fig. S1). Matched surface patches are aligned to the target site and scored with a second neural network, outputting an interface post-alignment (IPA) score to further improve the discrimination performance of the surface descriptors (see methods).

To benchmark our method, we assembled a test set composed of 114 dimeric complexes, which contained 31 complexes where the binding motif was a single alpha-helical segment and 83 where the binding motif was composed of less than 50% helical segments (Supplementary Fig. S3). As decoy sets, we used 1000 motifs (ranging from 600K-700K patches) which in the case of the helical set also had helical secondary structure and in the non-helical set were composed of two- and three-strand beta sheets.

We benchmarked MaSIF-seed relative to other docking methods to identify the true binder from the co-crystal structure in the correct orientation (<3 Å iRMSD) among 1000 decoys (Supplementary Fig. S4). MaSIF-seed identified the correct binding motif in the correct orientation as the top scoring result in 18 out of 31 cases, and 41 out of 83 cases for the helical and non-helical sets, respectively. While the best performing method, ZDock+ZRank2^22–24^ identified only 6 out of 31 as top results in the helical set, 21 out of 83 in the non-helical set. In addition to superior performances MaSIF-seed was considerably faster, showing speed increases between 20-200 fold, which mostly depend on the number of patches derived from each motif. In our benchmark we also performed comparisons with faster methods which showed much lower performances than ZDock+ZRank2 (Table 1 and Supplementary Table 1).

**Table 1.** Benchmark of MaSIF-seed against other docking methods in recovering the native binder in the correct conformation from co-crystal structures for 31 helix-receptor complexes or 83 non-helix seed-receptor complexes, discriminating between 1000 decoys. ^a^Benchmarked method. ^b-d^Number of receptors for which the method recovered the native binding motif (<3 Å iRMSD) within the ^b^top 1, ^c^top 10, and ^d^top 100 results. ^e^Number of receptors for which the method did not recover the native binding motif in the top 100 results. ^f^Average running time in minutes, excluding pre-computation time.

|  | Method<sup>a</sup> | # in top 1<sup>b</sup> | # in top 10<sup>c</sup> | # in top 100<sup>d</sup> | >100<sup>e</sup> | Avg time (m)<sup>f</sup> |
| --- | --- | --- | --- | --- | --- | --- |
| **Helical seeds** | MaSIF-seed | 18 | 18 | 20 | 11 | 15 |
|  | ZDock | 3 | 4 | 8 | 23 | 2715 |
|  | ZDock+ ZRank2 | 6 | 12 | 21 | 10 | 2946 |
| **Non-helical seeds** | MaSIF-seed | 41 | 47 | 49 | 34 | 118 |
|  | ZDock | 7 | 9 | 22 | 61 | 2206 |
|  | ZDock+ ZRank2 | 21 | 33 | 45 | 38 | 2400 |

An analysis of the cases where MaSIF-seed performed best showed that its success relied first on PPIs where the interaction site could be correctly identified by the method, and second to those where the majority of contacts lie on a radial patch at the interface core, and with a high shape complementarity in that region (Supplementary Fig. S5a). This is consistent with how MaSIF-seed was designed to capture protein interfaces using a radial geodesic patch.

Encouraged by MaSIF-seed’s speed and accuracy in discriminating the true binders from decoys based on rich surface features, we sought to design *de novo* protein binders to engage challenging and disease-relevant protein targets. We thus assembled a motif database including approximately 640 K structural fragments (402 M surface patches/fingerprints) with distinct secondary structures (approximately 390 K and 250 K of non-helical and helical motifs, respectively), extracted from the PDB (see methods). We computationally designed and experimentally validated binders against four structurally diverse targets: the receptor binding domain (RBD) of the SARS-CoV-2 spike protein where we identified a neutralization-sensitive site; the two partners of the PD-1/PD-L1 complex, an important protein interaction in immuno-oncology that displays a flat interface considered “hard-to-drug” by small molecules (Supplementary Fig. S6); CTLA-4, another important target for immuno-oncology. We show that our method can be applied to a variety of structural motifs as binding seeds (helical and non-helical), generating functional designs directly from the computational simulations.

### Targeting a predicted neutralization-sensitive site on SARS-CoV-2 spike protein

We applied our surface-centric approach to design *de novo* binders to target the SARS-CoV-2 RBD. First, we used MaSIF-site to predict surface sites on the RBD with high propensity to be engaged by protein binders. We selected a site distinct from the ACE2 binding region, but overlapping such that a putative binder could inhibit the ACE2-RBD interaction (Fig. 2a). At the time, there were no known binders to this site. We searched a subset of our database containing 140 million surface fingerprints derived from helical fragments to find binding seeds that could target the selected site. The 7713 binding seeds MaSIF-seed provided showed two prominent features: I) a contact surface devoid of residues with strong binding hotspot features (e.g. large hydrophobic residues); II) an equivalent distribution of binding seeds in two distinct orientations of the helical fragment, with the seeds binding at 180° from each other (Fig. 2b), hinting that both binding modes are plausible. Remarkably, both orientations of the binding seeds present very similar signatures at the surface fingerprint level (Supplementary Fig. S7) and at the sequence level (Fig. 2b).

**Figure 2:**
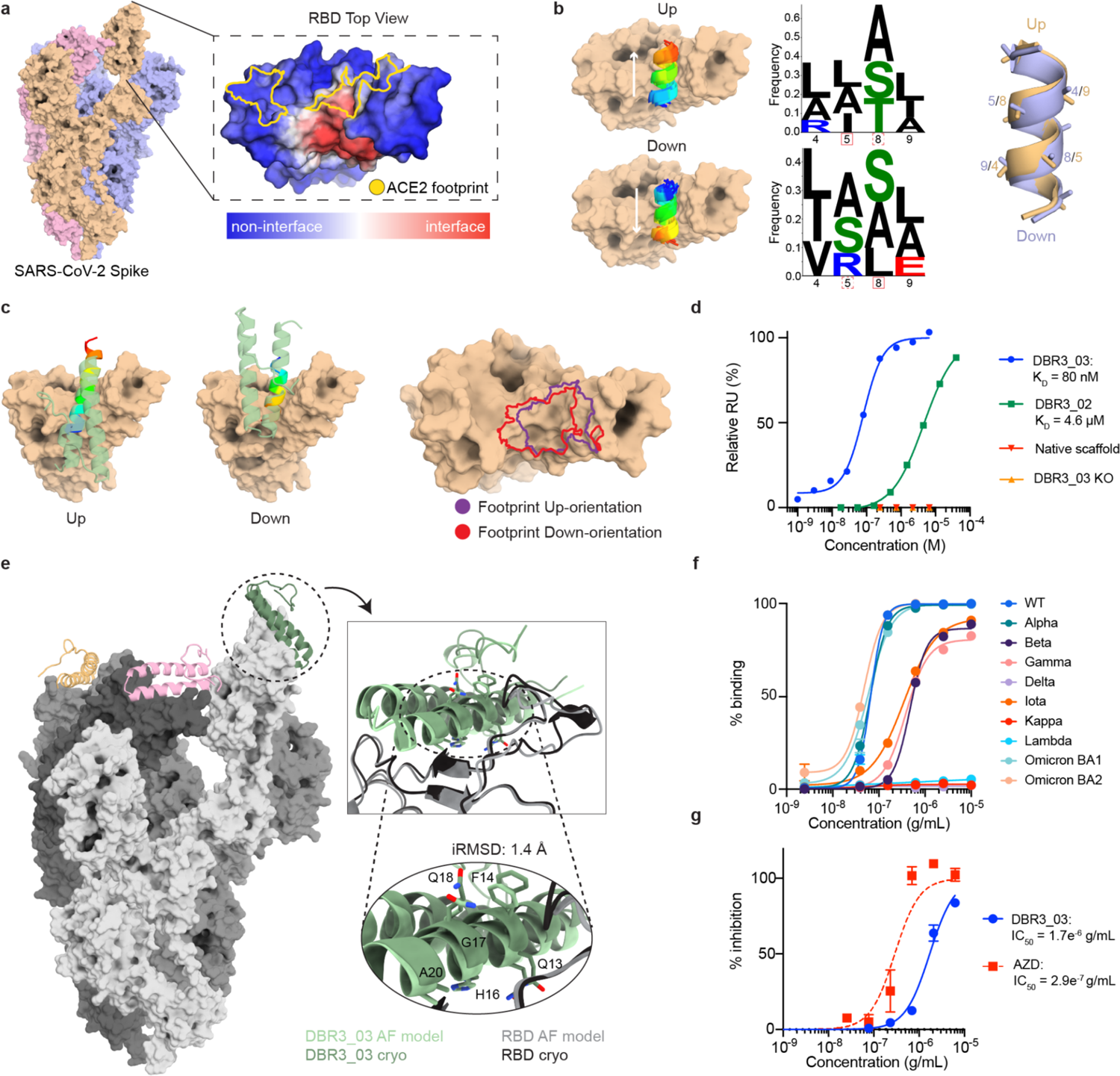
Design and optimization of a SARS-CoV-2 binder targeting the RBD. **a**, MaSIF-site prediction of interface propensity of the RBD. The ACE2 binding footprint (yellow outline) is distinct from the predicted binding site (red). **b**, MaSIF-seed predicts helical seeds that cluster into anti-parallel orientations, referred to as up or down configurations. Sequence logo plots highlight the similarity between the sequences of the two seed clusters, regardless of orientation. **c**, The scaffold (PDB ID: 5vny) used to make DBR3_01 allows for binding in up- or down-orientation, sharing similar footprints. **d**, SPR data of improved DBR3 binders with controls. DBR3_03 has an affinity of 80 nM with RBD. **e**, A Cryo-EM structure (dark green) aligns to the AlphaFold prediction with an iRMSD of 1.4 Å. The trimeric spike protein (gray) has one DBR3_03 bound per RBD (orange, pink, green). **f**, Fc-DBR3_03 binds to spike protein from most variants of concern, except those with the L452R mutation. EC_50_ of DBR3_03 are listed in Supplemental Table 3. **g**, Fc-DBR3_03 neutralizes live omicron-virus in cell-based inhibition assays with an EC_50_ of 1.7E^-6^ g/mL, compared to the AstraZeneca (AZD, 8895 and 1061) mix that has an EC_50_ of 2.9e^-7^ g/mL.

We synthesized one of the top ranked binding seeds as a linear peptide, but no binding interaction was detected by Surface Plasmon Resonance (SPR) (Supplementary Fig. S8). Therefore, using the Rosetta MotifGraft protocol we identified several protein scaffolds compatible with both binding modes of the seed (Fig. 2c), transplanted the seed hotspot side chains from a top-ranking seed onto the scaffolds, and used Rosetta to optimize the binder interface (Fig. 1c). Sixty-three designs based on twenty scaffolds, ranging from 7 to 23 mutations relative to the native proteins, were screened with yeast display (Supplementary Fig. S9). From this initial round of designs, DBR3_01 showed weak binding in yeast display experiments. Moreover, binding of DBR3_01 was competitive with soluble ACE2 (Supplementary Fig. S9), suggesting that the binder was targeting the correct RBD site. Furthermore, DBR3_01 showed slightly increased binding compared to the native scaffold protein and a double point mutant on the designed interface residues, further supporting that the seed residues were participating in the binding interaction (Supplementary Fig. S9, Supplementary Table 2). Next, we sought to improve the binding affinity of the design by performing two mutagenesis libraries: first, a directed library in the designed interface was prepared (Supplementary Fig. S10), which yielded DBR3_02 with 4 mutations and a K_D_ of 4.6 µM determined by SPR (Fig. 2d, Supplementary Fig. S10); second, we screened a site saturation mutagenesis (SSM) library which resulted in the enrichment of 3 point mutants, one of which overlapped with a mutation from the first library (Supplementary Fig. S11). Adding these 3 mutations to DBR3_02 resulted in DBR3_03 that showed a K_D_ of 80 nM and was folded and stable (Fig. 2d, Supplementary Fig. S12). Here, we started from a computationally designed binder with very low affinity as observed with yeast display, yet undetectable by SPR, and after introducing 6 mutations we observed an improvement greater than 60 fold in binding affinity. The mutations all occurred in the binding helix of the design. Of these mutations, A17G and S20A, residing in the core of the interface, appear to have relieved steric clashes and reduced buried unsatisfied polar atoms, respectively.

To structurally characterize the binding mode of DBR3_03 we solved a cryo-EM structure of the design in complex with the trimeric spike protein at 2.9 Å local resolution (Fig. 2e and Supplementary Fig. S13-15). The structure confirmed the predicted binding sites on both partners. Importantly, the binder adopted the orientation of the helical binding seed that was marginally less favored by MaSIF’s fingerprint descriptors (down-orientation) (Fig. 2b). Interestingly, the initial design DBR3_01 showed similar metrics when the interfaces were analyzed in both directions (Supplementary Fig. S7), pointing to known limitations of surface fingerprints in unbound docking type of problems^17^. This led us to attempt another state-of-the-art protein docking method, AlphaFold (AF) multimer^25^ to predict the complex of DBR3_03 with the spike RBD and obtained a 1.4 Å iRMSD between the AF prediction and the experimental structure (Fig. 2e). This result presents a powerful demonstration of the synergies between machine learning techniques purely based on structural features and those that leveraged large sequence-structure datasets for structure prediction tasks. At the structural level DBR3_03 engages the RBD with a 1452 Å^2^ of buried interface area (surface area buried on both sides of the complex), which is much smaller than the average buried surface area of antibodies (approximately 2071 ± 456 Å^2^ ^26^), yet still results in a high affinity interaction. The designed interface lacks canonical hotspot residues and engages the RBD through small residues and is composed of 21% backbone and 79% side chain contacts. Given the pandemic situation with SARS-CoV-2 and the general need for rational design of protein-based therapeutics to fight viral infections, we next engineered an Fc-fused DBR3_03 (Fc-DBR3_03) construct and tested its neutralization capacity on a panel of SARS-CoV-2 variants in virus-free and pseudovirus surrogate assays (Fig. 2f,g, Supplementary Fig. S16, Supplementary Table 3)^27^. We compared the breadth and potency of our design to those of clinically approved monoclonal antibodies. In virus-free assays we observed that Fc-DBR3_03 had comparable potency to that of Imdevimab (REGN10987), an antibody used clinically, for the WT spike and bound to the omicron strain while RGN87 did not (Supplementary Fig. 16). Neutralization activity in pseudovirus assays was tested and Fc-DBR3_03 neutralized omicron, albeit less potently than the AstraZeneca (AZN) clinically approved antibody mix (Fig. 2g). A cryo-EM structure showed that the binding mode was nearly identical (1.4 Å backbone RMSD) between DBR3_03-WT-RBD complex and DBR3_03-omicron-RBD complex (Supplementary Fig. S17-19). Importantly, Fc-DBR3_03 showed a very broad reactivity to many SARS-CoV-2 variants (Fig. 2f) which is attributable to the sequence conservation of the targeted site and the small binding footprint of the design. The design was sensitive to the L452R/Q mutation present in the delta, lambda and kappa variants (Supplementary Fig. S16b, Fig. 2f), but introducing a single point mutation (L24G) to relieve the clash between L452R and the binder led to the design binding to delta (Supplementary Fig. S16). Our results highlight the value of the surface fingerprinting approach to reveal target sites in viral proteins and for the subsequent design of functional antivirals with broad activity.

### Targeting flat surface sites in immune checkpoint receptors

Surface sites presenting flat structural features are difficult to target with small molecule drugs, leading to their categorization as undruggable. To test our fingerprint-based approach, we sought to design binders to target the PD-1/PD-L1 interaction, which is central to the regulation of T-cell activity in the immune system^28^. We used MaSIF-site to find high propensity protein binding sites in PD-L1, and unsurprisingly, the identified site overlapped significantly with the native binding site engaged by PD-1 (Fig. 3a). This site is extremely flat at the structural level, ranking in the 99^th^ percentile in terms of interface flatness (ranked #7 among 1068 transient interfaces, details in methods) (Supplementary Fig. S20), one of the dominant structural features that makes this site hard-to-drug by small molecules. Next, we used MaSIF-seed to find binding motifs to engage the site, among the top results helical motifs clustered in both orientations packing in the beta-sheets of PD-L1 (Supplementary Fig. S21). In the most populated cluster (Supplementary Fig. S21), we observed sequence convergence for a 12 residue fragment (Fig. 3b). We then used Rosetta MotifGraft to search for putative scaffolds to display this fragment and used RosettaDesign to optimize contacts at the interface. We tested 16 designs based on 5 different scaffolds for binding to PD-L1 on the surface of yeast. Two designs based on two different scaffolds showed low binding signals (Supplementary Fig. S22), which we refer to as DBL1_01 and DBL2_01 (Fig. 3c). The specificity of the interaction was confirmed by testing hotspot knockout controls of each design (Supplementary Fig. S22). To improve the binding affinity of DBL1_01 we constructed a combinatorial library with mutations in the predicted binding region, while maintaining the hotspot residues predicted by MaSIF-seed (Supplementary Fig. S23). From this library we selected a variant, DBL1_02 with 5 mutations found mostly in the interface rim of the design and improving the formation of polar contacts. The most substantial change occurred at position 53, a mutation of alanine to glutamine that introduces a hydrogen bond with PD-L1 (Supplementary Fig. S23). To improve the design’s expression and stability we constructed a second library targeting residues in the protein core to optimize core packing (Supplementary Fig. S23). Combining mutations from both libraries, we obtained DBL1_03 with 11 mutations from the starting design, which was folded and monomeric in solution, and showed a binding affinity of 2 µM (Fig. 3d, Supplementary Fig. S12), comparable to that of PD-1 (K_D_ = 8.2 µM)^29^. To further assess the optimality of each residue at the interface of the designed binder we screened a SSM library sampling 19 positions, based on DBL1_03. The most relevant positions are shown in Figure 3f (all positions in Supplementary Fig. S24). The SSM results revealed that the four hotspot residues placed by MaSIF-seed were crucial, as any other residue was deleterious for binding (Fig. 3f). However, in the interface rim many mutations could provide affinity improvements strongly suggesting that this region of the interface was suboptimal (Fig. 3f). Based on these data, we generated the DBL1_04 variant which resulted in a 10-fold increase of the binding affinity showing a K_D_ of 256 nM to PD-L1 (Fig. 3d). Both DBL1_03 and DBL1_04 showed cell-surface binding, comparable to PD-1, on cells expressing PD-L1. The specificity of the designed interaction was confirmed by the binding inability of single-residue mutants at the interface (Supplementary Fig. S24).

**Figure 3:**
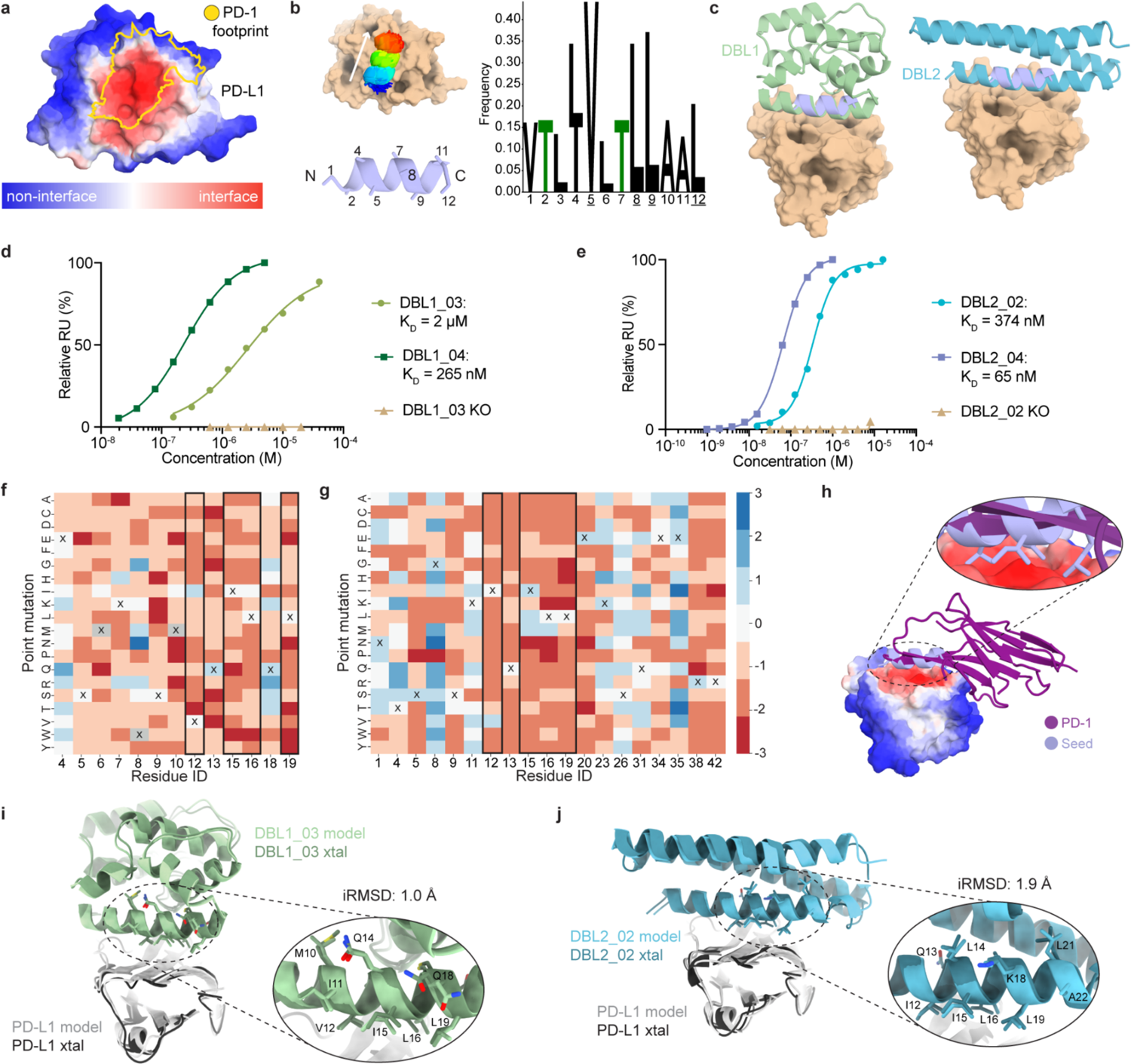
*De novo* design and optimization of PD-L1 binders targeting a flat surface. **a**, MaSIF-site prediction of interface propensity of PD-L1. The predicted interface (red) overlaps with the binding site of the native interaction partner PD-1 (yellow). **b**, Helical seeds were predicted by MaSIF-seed and clustered. The dominant cluster showed strong amino acid preferences (Z-score > 2). Hotspot residues are underlined. **c**, Binders based on two different scaffold proteins utilizing the selected seed were identified. **d**, Binding affinities of DBL1 designs after combinatorial (light green) and SSM library optimization (dark green), measured by SPR. Mutation of a hotspot residue (V12R) ablates binding of DBL1_03 (wheat). **e**, Binding affinities of DBL2 designs after combinatorial (light blue) and SSM library optimization (dark blue), measured by SPR. Mutation of a hotspot residue (V12R) knocks out binding of DBL2_02 (wheat). **f**, SSM for regions of interest in the binding interface of DBL1_03. X indicates the original residue of DBL1_03 and hotspot residue positions are framed in black boxes. Blue indicates enrichment in the binding population, while red shows enrichment in the non-binding population. **g**, SSM data in the binding interface of DBL2_03. The X indicates the original residue of DBL2_02. **h**, Binding mode of the selected seed in comparison with the native interaction partner PD-1. **i**, Crystal structure of DBL1_03 in complex with PD-L1. The computational model (light green) is aligned with the crystal structure (dark green). The inset shows the alignment of the residues in the binding seed. **j**, Crystal structure of DBL2_02 in complex with PD-L1. Shown by aligning the computational model (light blue) with the crystal structure (dark green). The inset shows the alignment of the residues in the binding seed represented in sticks.

The second lead design, which utilizes the same seed but is based on a different scaffold, DBL2_01, could not be solubly expressed and therefore we designed a combinatorial library to improve expression and binding affinity (Supplementary Fig. S23). From this library we isolated the variant DBL2_02 which had six mutations and expressed in *E. coli*. From the six mutations, three were predicted to be in the interface (Y23K, Q35E, Q42R) and improved binding affinity by forming additional salt bridges with PD-L1 (Supplementary Fig. S23). The K_D_ to PD-L1 determined by SPR was 374 nM, more than 10-fold higher than the native ligand PD-1. Since both designs shared the same binding seed we transplanted the SSM mutations of the DBL1_04 design and generated the DBL2_03, which showed a 3-fold improvement in binding affinity (K_D_ = 120 nM) (Supplementary Fig. S25), indicating that the binding seed was engaging PD-L1 in a similar fashion to that of DBL1_03. To further assess the influence of each residue in the designed binding interface we performed an SSM analysis on 19 interface residues of DBL2_03 (Fig. 3f, Supplementary Fig. S25). The SSM profile reiterated that the hotspot residues placed by MaSIF-seed were very restricted in variability, showing that these residues were accurately predicted. In contrast, several positions on the interface rim were suboptimal and mutations to polar amino acids resulted in affinity enhancements. Based on the SSM data, we generated the DBL2_04 design with additional polar mutations (Fig. 3g, Supplementary Fig. S25) which showed an improved K_D_ of 65 nM (Fig. 3e). To experimentally validate the binding mode, we co-crystallized the designs with PD-L1 (Supplementary Fig. S26). Overall for both designs, the structures (Fig. 3i,j) showed excellent agreement with our computational models with 0.8 Å and 2.0 Å for the overall backbone and 1.0 Å and 1.9 Å for the full atom interface RMSDs of DBL1_03 and DBL2_02, respectively, showing an exquisite accuracy of the predictions in the interface region. The buried interface area of the designs with PD-L1 was between 1424 Å^2^ and 1438 Å^2^, compared to 1648 Å^2^ for the buried interface area of PD-1 (PDB ID: 4ZQK). The chemical composition of the designed interface is similar in both designs, ∼59% of the surface area is hydrophobic and the remaining area is hydrophilic for DBL1_03 and correspondingly for DBL2_02. These values are comparable to those of the PD-1/PD-L1 interaction (52% hydrophobic surface), showing that we have designed interfaces with similar chemical compositions of the native interaction using a distinct backbone conformation (Fig. 3h). The discovery of novel binding motifs by MaSIF-seed is striking when comparing the backbone motif used by the native PD-L1 binding partner, PD-1, and the designed binders. While the native PD-1 uses a beta-hairpin to engage the site, the designed binders do so through an alpha-helix motif, illustrating the capability of our approach to explore outside of the structural repertoire of native binding motifs. The general trend arising from the designed PD-L1 binders is that despite the accurate predictions of core residues in the interface, through mutagenesis studies, the designed polar interactions are suboptimal. To address these and other limitations of our computational approach, we performed additional computational design steps to improve the pipeline and tested it on the design of binders to target PD-1.

### One-shot design of *de novo* protein binders with native-like affinities

Despite the successes in designing site-specific binders to engage two different targets, the computational designs still required *in vitro* evolution to enable expression and detectable binding affinities that could be biochemically characterized. To address these issues, we used a structurally diverse library of binding seeds (helical and beta-sheet motifs) and assembled a more comprehensive design pipeline (Fig. 4a) performing: I) sequence optimization of selected seeds; and II) biased design for polar contacts in the scaffold interface^30^. To test this approach, we designed *de novo* binders to target three proteins (PD-L1, PD-1 and CTLA-4). For each of the design targets we selected the top 2000 designed sequences according to several structural metrics (see methods) and tested them using yeast display coupled with deep sequencing readout. According to our deep sequencing readout we obtained binders for all three targets using diverse structural motifs to mediate the binding interaction (Supplementary Table 4). Several binders were biochemically characterized to varying degrees. For PD-1 we found three designs based on *de novo* miniprotein^31, 32^ scaffolds with interfaces mediated by helical motifs (DBP13_01, DBP40_01 and DBP52_01) (Fig. 4b, Supplementary Fig. S27) that showed a moderate to strong binding signal on the surface of yeast. The most promising candidate binding to PD-1, DBP13_01, was investigated in more detail (Fig. 4b-e). To confirm whether the binding interaction was mediated through the designed interface, we tested several control constructs, which included the native miniprotein scaffold and DBP13_01 variants with predicted knockout mutations (Fig. 4b), all of which abolished binding (Fig. 4c). The interaction site on PD-1 was further probed via a competition assay with Nivolumab^33^, which blocked the DBP13_01/PD-1 interaction as expected due to the overlapping binding footprints (Supplementary Fig. S28). DBP13_01 did not bind to a close sequence homologue (porcine PD-1) supporting the specificity of the designed interactions (Supplementary Fig. S28). The DBP13_01/PD-1 interaction showed a K_D_ of 4.2 ± 2 µM (n = 3, Fig. 4d) as determined by SPR, similar to the affinity of the native PD-L1/PD-1 interaction (K_D_ = 8.2 µM)^29^. This was a promising result given that the design was not subjected to experimental optimization by *in vitro* evolution. Next, we performed an SSM experiment and observed that mutations at the predicted core interface positions (L23, L27, I30, M31) were generally deleterious for binding, supporting the structural and sequence accuracy of the design (Fig. 4e, Supplementary Fig. S29). Moreover, we readily improved the affinity to sub-micromolar by introducing two mutations identified in the SSM data (M31F+H33S, DBP13_02) (Fig. 4d). The predicted complex structure by AlphaFold Multimer (AF) was in agreement with that of MaSIF, with an interface footprint that is largely overlapping with the designed residues, and 3.3 Å of backbone RMSD and 2.9 Å of interface full atom RMSD (Supplementary Fig. S30). Although these results are supported by the SSM data, they are a predictive exercise and cannot be interpreted as absolute evidence that the designed binding mode is occurring, which ultimately will require an experimental structure.

**Figure 4:**
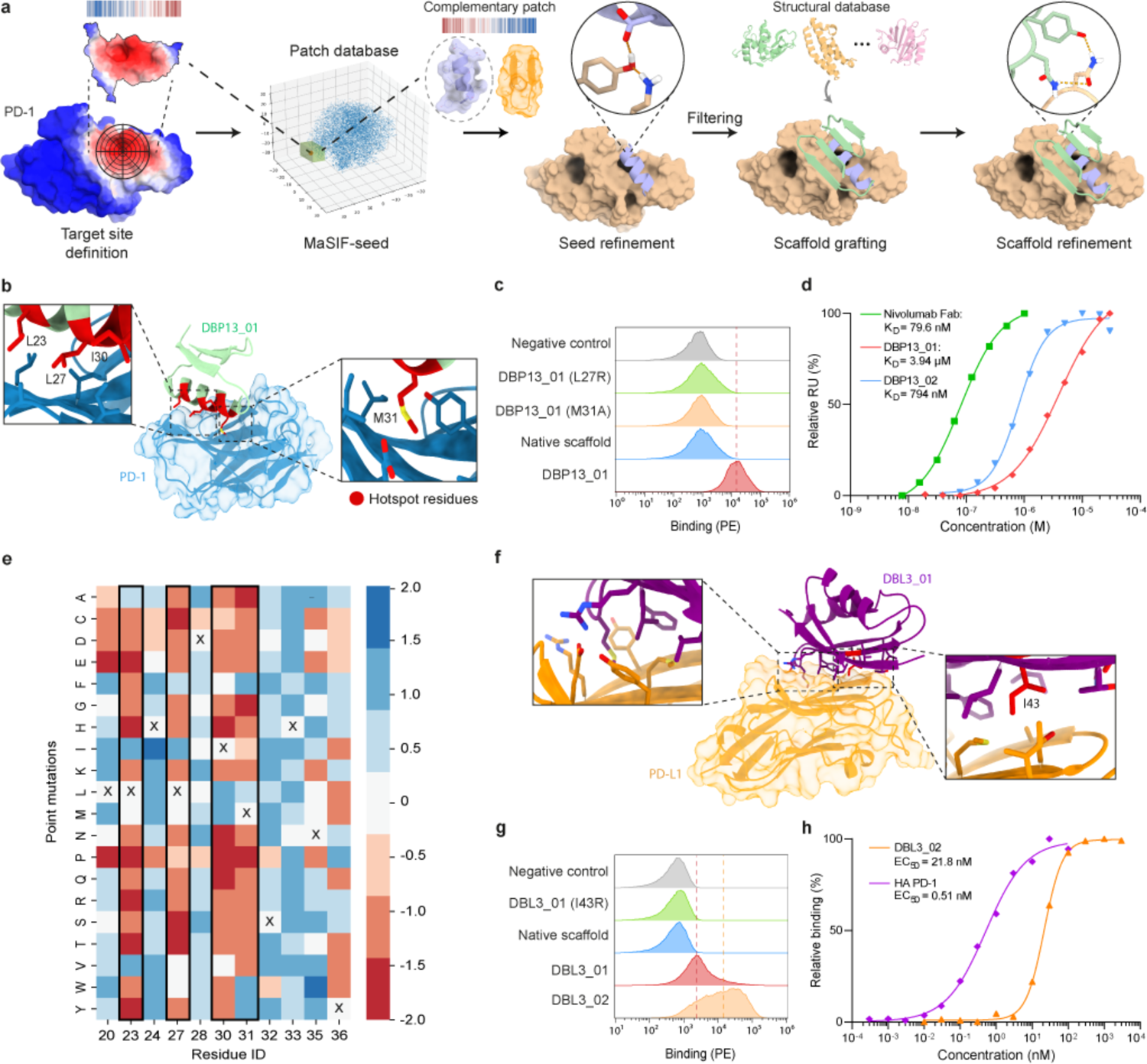
Optimized workflow and *de novo* binders for PD-1. **a**, Improved design computational workflow where two steps of design are used, at the seed and at the scaffold level, with an emphasis on building new hydrogen bond networks **b**, PD-1 (blue) targeted by DBP13_01 (green) with highlighted hotspots residue from the binding seed (red). The two inset panels show crucial residue for binding. **c**, Histogram of the binding signal measured by flow cytometry for DBP13_01, the native miniprotein scaffold, two variants of DBP13_01 with crucial residues mutated, and a negative control with unlabeled yeast. Dashed line indicates the geometric mean of DBP13_01 binding signal. **d**, Binding affinities determined by SPR of Nivolumab Fab (green squares), DBP13_01 (red diamonds) and DBP13_02 (blue triangles). DBP13_01 dissociation constant was obtained with 3 independent measurements. **e**, SSM heatmap showing interface residues and the enrichment of each point mutation. “X” indicates the original amino acids in DBP13_01. Blue indicates enrichment in the binding population, while red shows enrichment in the non-binding population. Hotspot residues are highlighted with a black box. **f**, PD-L1 (orange) targeted by DBL3_01 (purple). The two inset panels show a close-up on interface residues, including one crucial residue tested for knock-out mutants (I43, red). **g**, Histogram of the binding signal measured by flow cytometry for DBL3_01, DBL3_02, the native protein scaffold, one knock-out mutant and a negative control with unlabeled yeast. **h**, PD-L1 ligand titration on yeast displaying DBL3_02 (orange triangle) or high-affinity PD-1 (HA-PD-1, purple diamonds).

Similarly, we experimentally confirmed the specificity of a beta-sheet based-binder to PD-L1 (DBL3_01) (Fig. 4f) with a predicted knock-out mutant and a competition assay with high-affinity PD-1 (Fig. 4g and Supplementary Fig. S28). These data were supported by an AF prediction matching our design model with a 0.97 Å backbone RMSD (Supplementary Fig. S30). Binding to PD-L1 was further improved on yeast by mutating two exposed cysteines to serines in the scaffold, which may stabilize the protein and avoid unwanted disulfide bonds (DBL3_02, Fig. 4g and Supplementary Fig. S28). This design adopts a different backbone conformation than the native PD-1:PD-L1 interaction which further demonstrates MaSIF-seed’s ability to generalize beyond interactions found in nature (Supplementary Fig. S28). We also estimated the affinity on a yeast display-based assay determining an apparent K_D_ of 21.8 nM, 42.7-fold higher than the known high-affinity PD-1, which has been reported to have a true K_D_ of 110 pM^34^ (Fig. 4h).

We also performed experimental characterization for two other binders targeting PD-L1 (DBL4_01) and CTLA-4 (DBC2_01) and observed that the binding interactions are specific to targeted sites by competition and mutagenesis experiments performed using yeast display (Supplementary Fig. S28 and S31). It is important to note that for several of these binders the AF predictions were not in agreement with our models but that nevertheless the experimental results provide solid evidence that the correct interfaces are involved in the designed interactions (Supplementary Table 4).

Overall, the results show that by starting the interface design process driven by surface fingerprints and introducing additional features of native interfaces (e.g. hotspot optimization, polar contacts) we can design site-specific binders, using a variety of structural motifs with native-like affinities purely by computational design.

## Discussion

Physical interactions between proteins in living cells are one of the hallmarks of function^35^. Our incomplete understanding of the complex interplay of molecular forces that drive PPIs has greatly hindered the comprehension of fundamental biological processes as well as the capability to engineer such interactions from first principles. It has been particularly challenging for protein modeling methodologies that use discrete atomic representations to perform *de novo* design of PPIs^2, 6, 10, 11^. In large part, this is due to the small number of molecular interactions involved in most protein interfaces and to the very small energetic contributions that determine binding affinities, making physics-based energy functions less reliable^36^. To address this gap, we developed an enhanced data-driven framework to represent proteins as surfaces and learn the geometric and chemical patterns that ultimately determine the propensity of two molecules to interact. We proposed a new geometric deep learning tool, MaSIF-seed, to overcome the PPI design challenge by both identifying patches with a high propensity to form buried surfaces and binding seeds with complementary surfaces to those patches. By computing fingerprints from protein molecular surfaces, we rapidly and reliably identify complementary surface fragments that can engage a specific target within 402 million candidate surfaces. This, in practice, solves an important challenge in protein design by efficiently handling search spaces of daunting scales.

The identified binding seeds were then used as the interface driving core to design novel binding proteins against challenging targets: a novel predicted interface in the SARS-CoV-2 spike protein, which ultimately yielded a SARS-CoV-2 inhibitor, PD-1/PD-L1 protein complex and CTLA-4, exemplifying sites that are difficult to target with small molecules due to its flat surface. Several designed binders showed close mimicry to computationally predicted models and achieved high binding affinities, often, after experimental optimization. In the case of purely computationally designed binders, the PD-1 binder showed low micromolar affinity without experimental optimization, which is the range of many native PPIs^37^, and several other binders targeting PD-L1 and CTLA-4 were shown to be specific to the targeted sites. By using surface fingerprints, we identified novel structural motifs that can mediate *de novo* PPIs presenting a route to expand the landscape of motifs that can be used to functionalize proteins and be critical for the *de novo* design of function.

For all targets, the original binding seed arguably provided the principal driver of molecular recognition representing the design’s binding interface core (Supplementary Fig. S32), maintaining a high surface similarity in this region between the original seed and the final design (Supplementary Fig. S33). However, contacts at the buried interface region are necessary though in most cases, likely not sufficient for high affinity binding, and in the three designed binders for PD-L1 and RBD, optimization of the polar interface rim through libraries was necessary to improve binding to a biochemically detectable range (K_D_ at the micromolar level). Our *de novo* designs agree with previous findings^6, 38^ that small changes in the polar interface rim (for example in the hydrogen bond network surrounding the interface) can result in substantial differences in binding affinities. Encouragingly, by using a larger and more structurally diverse library of binding seeds together with an optimized design pipeline we obtained several *in silico* only designed binders to a variety of targets, which represents a major step forward for the robust design of *de novo* PPIs.

In our study several limitations of the approach became evident, namely the absence of conformational flexibility and adaptation of the protein backbone to mutations and the difficulty of designing polar interactions that balance the hydrophobic patches of the interface contributing for affinity and specificity, which has also been observed by other authors^21, 38, 39^. In future methodological developments, neural network architectures could be optimized to capture such features of native interfaces. The emergence of generative algorithms that can construct backbones conditioned to the target binding sites or the seed motifs, as recently described by other groups^40, 41^, present another exciting route where our conceptual framework based on surfaces is likely to become more useful to overcome important challenges on the design of molecular recognition.

Here we presented a surface-centric design approach that leveraged molecular representations of protein structures based on learned geometrical and chemical features. We showed that these structural representations can be efficiently used for the design of *de novo* protein binders, one of the most challenging problems in computational protein design. We anticipate that this conceptual framework for generation of rich descriptors of molecular surfaces can open possibilities in other important biotechnological fields like drug design, biosensing or biomaterials in addition to providing a means to study interaction networks in biological processes at the systems levels.

## Methods

### Computing buried surface areas in protein complexes

A dataset of protein-protein interactions was downloaded from the PDBBind database^42^ containing all interactions with a reported affinity stronger than 10 µM; since these PPIs have a reported affinity, all were assumed to be transient. The PDBBind database does not report the chains involved in the interaction with the reported affinity; thus, for simplicity, only those complexes containing exactly two chains in the PDB crystal structure were considered for the analysis.

The MSMS program^43^ was used to compute all molecular surfaces in this work (density = 3.0, water radius = 1.5 Å). Since MSMS produces molecular surfaces with highly irregular meshes, PyMESH^44^ was used to further regularize the meshes at a resolution of 1.0 Å. For a given protein subunit that appears in a complex, we define the subunit’s buried surface as the patch that becomes inaccessible to water molecules upon complex formation. Since in our implementation a surface is defined by a discretized mesh, we compute the buried surface region as follows. The buried surface of both the subunit and the complex are first independently computed. Then, the minimum distance between every subunit surface vertex and any complex surface vertex are computed. Subunit vertices that are farther than 2.0 Å from a vertex in the surface of the complex are labeled as part of the buried surface, as these vertices no longer exist in the surface of the complex. The size of buried areas was determined by computing the area of each vertex labeled as a buried surface vertex.

We note that computing buried surface areas using this method can result in measurements that are different from those widely used in the field, which use the solvent accessible surface area and count the buried interface of all subunits into a single value (the buried SASA area). Here we use the molecular surface (also known as solvent excluded surface) and count a single subunit. Therefore, while in Supplementary Fig. 1 we show areas computed using this method to compare to patch sizes, throughout the rest of this work we refer to the more widely used buried SASA areas.

### Patch generation in the MaSIF framework

#### Decomposing protein surfaces into radial patches

In order to process protein surface information, all molecular surfaces were decomposed into overlapping radial patches. This means that each vertex on the surface becomes the center of a radial patch of a given radius. To compute the geodesic radius of patches, throughout this work we used the Dijkstra algorithm^45^, a fast and simple approximation to the true geodesic distance in the patch. We used a radius size of 12 Å for patches, limited to at most 200 points, which we found corresponds roughly to 400 Å^2^ (Supplementary Fig. S2), a value close to the median size of the buried interface of transient interactions (Supplementary Fig. S2). Exceptionally, for the MaSIF-site application (described below) we limited the patch to 9 Å or 100 points to reduce the required GPU RAM for this application^17^.

#### Computing angular and radial coordinates

An essential geometric deep learning component in our pipeline is to compute angular and radial coordinates in the patch that enable MaSIF to map features in a 2D plane. The radial coordinate is computed using the Dijkstra algorithm, where the geodesic distance (meaning the distance taken to ‘walk’ along the surface) from the center of the patch to every vertex is computed. To compute the angular coordinate, all pairwise geodesic distances between vertices in the patch are computed, and then the multidimensional scaling algorithm^46^ in scikit-learn^47^ is used to map all vertices to the 2D plane. Then, a random direction in the 2D plane is computed as the 0° frame of reference, and the angle of every vertex in the plane with respect to this frame of reference is computed. Computing the angular and radial coordinates is the slowest step in the MaSIF precomputation. However, we have provided experimental code to compute these coordinates much faster in our github repository under a branch called “fast-masif-seed”.

#### Geometric and chemical features used to describe molecular surfaces

Each point in a patch of the computed molecular surface was assigned an array of two geometric features (shape index^48^, distance-dependent curvature^49^), and three chemical features (hydrophobicity^50^, Poisson-Boltzmann electrostatics^51^, and a hydrogen bond potential^52^). These features are identical to those described in Gainza *et al*^17^.

#### Computing the largest circumscribed patch in buried surface areas

From each labeled interface point, we used the Dijkstra algorithm to compute the shortest distance to a non-interface point. The interface point with the largest distance to a non-interface point was labeled as the center of the interface, and the distance to the nearest non-interface point as the radius of the largest circumscribed patch.

### Calculation of surface planarity

The surface planarity of all target interfaces, with respect to a database of PPIs (Supplementary Fig. S20) was calculated as follows. 690 PPIs crystallized as dimers from the PDBBind database were used as the dataset, resulting in 1380 interfaces as each chain was analyzed separately. Interfaces with an approximate area lower than 150 Å^2^ or more than 1000 Å^2^ were discarded, resulting in 1068 interfaces. The vertices in the buried interface area of each chain were computed, as explained in section Computing buried surface areas in protein complexes above, and the 3D coordinates of those vertices in the interface were extracted from each chain. Then, the multidimensional scaling method^46^ from scikit-learn^47^ was used to position interface vertices in a 2D plane, with the optimization goal of maintaining the distances between all pairs of vertices as close as possible in the 2D embedding as they were in 3D space. The root mean square difference between the distances in the original 3D space vs. the 2D space was used as the measure of planarity. Interfaces that are very planar in 3D have small values under this metric as an embedding in 2D preserves the distance between vertices, while non-planar interfaces have larger values as an embedding in 2D must significantly alter their 3D distances.

### Geometric deep learning layer in MaSIF

Geometric deep learning enables the application of traditional techniques from deep learning to data that does not lie in Euclidean spaces, such as a protein molecular surface. At the core of MaSIF lies a mapping from a molecular surface patch to a 2D Euclidean Tensor. The mapping is performed through a learned soft polar grid around each patch center vertex, using the angular and radial coordinates. Once the mapping is performed, a traditional convolutional neural network layer is performed, with an angular max pooling layer, which deals with the rotation ambiguity of geodesic patches. Further details on these techniques are detailed in Gainza *et al*^17^ and Monti *et al*^53^.

### MaSIF-site - prediction of protein interaction sites

The MaSIF-site tool^17^ was trained to predict areas with propensity to form protein-protein interactions on the surface of proteins. Here, MaSIF-site was used to predict surface areas with propensity to form a PPI in 114 targets of our benchmark (Supplementary Fig. S4) and all the design targets (SARS-CoV-2 RBD, PD-L1, PD-1 and CTLA4). MaSIF-site receives as input a protein decomposed into patches and outputs a per-vertex regression score on the propensity of each point to become a buried surface area within a PPI. MaSIF-site computes a regression score on each point of the surface, yet it becomes necessary to identify the precise patch that we will use to define each interface. Thus, to select interface patches in target proteins, the output of MaSIF-site was decomposed into 12 Å overlapping patches, and the per-vertex prediction for all points in the patch are averaged to obtain a score for each patch.

### Training of MaSIF-site

MaSIF-site was trained on a database of protein-protein interactions sourced from PRISM^54^, PDBBind^42^, the ZDock benchmark^55^, and SabDab^56^. Proteins from these databases that failed to run through the MaSIF pipeline due to, for example, too many incomplete residues in the deposited structure, were discarded. Each instance of these databases, which we refer to as ‘subunits’ could consist of one or multiple chains (e.g., an antibody), and was crystallized in complex with a partner subunit. In total 12002 subunits from deposited structures passed the threshold. These subunits were then clustered by sequence identity at 30% identity and up to one representative from each cluster was selected, resulting in 3362 subunits. Then, a matrix of all pairwise TM-scores for this set was computed, and a hierarchical clustering algorithm was used on this matrix to split the dataset into 3004 subunits for the training set and 358 for the testing set.

The molecular surface for each subunit was computed using MSMS^43^ and the buried interface area was labeled as described above. The architecture of MaSIF-site (Supplementary Fig. S1b and further described in Gainza *et al*^17^) consisted of three layers of geodesic convolution. The network received as input the full surface of a protein (with batch size of 1) decomposed into overlapping patches of size 9 Å. During training, each vertex of the input was labeled with the ground truth, with a value of 1 if the vertex belonged to the buried area and a label of zero otherwise. The output of the network is a per-vertex assignment between 0 and 1 for the prediction of that vertex on whether it belongs to the buried surface area or not. A sigmoid activation function was used as the output layer, and a binary cross function as the loss function. Adam^57^ was used as the optimization function. MaSIF-site was implemented in Tensorflow^58^, and trained for 40 hours on a single GPU machine, which allowed for 43 epochs. The MaSIF-site neural network implementation in Tensorflow contains a total of 9267 parameters.

### Masif-search - identifying complementary surfaces using fingerprints

MaSIF-search^17^ was used to compute fingerprints for every overlapping patch in proteins of interest. MaSIF-search was trained on a dataset of 6001 protein-protein interactions (described in Gainza *et al*^17^) to receive as input the features of the target, a binder, and a random patch from a different protein. MaSIF-search was designed as a Siamese neural network architecture^18^ trained to produce similar fingerprints for the target patch vs. the binder patch, and dissimilar fingerprints for the target patch vs. the random patch. In order to decrease training time and improve performance, the features of the target were multiplied by −1 (with the exception of hydropathy), turning the problem from one of complementarity to one of similarity.

### Training of MaSIF-search

MaSIF-search was trained on a database of 6001 protein-protein interactions in co-crystal structures sourced from PRISM^54^, PDBBind^42^, the ZDock benchmark^55^, and SabDab^56^. A split between the training and testing set was performed by extracting the atoms at the interface for all 6001 PPIs and computing a TM-score between all pairs using TM-align. A hierarchical clustering algorithm was used to cluster the pairwise matrix, which was used to split the data into a training set of 4944 PPIs and a testing set of 957 PPIs. As in MaSIF-site, each side of the interaction could consist of one or multiple chains (e.g. an antibody), and we refer to each side as a subunit. In each PPI, pairs of surface vertices within 1.0 Å of each other were selected as interacting pairs.

MaSIF-search produces fingerprints for patches with a radius of up to 12.0 Å in geodesic distance from a central vertex, and is trained to make these patches similar for interacting patches and dissimilar for non-interacting patches (Fig. 1a, Supplementary Fig. S1). We find that MaSIF-search performs best when trained on interacting pairs that lie in the center of highly complementary interfaces and these pairs were filtered to remove points outside of the interfaces or in interfaces with poor complementarity (further described in Gainza *et al*^17^). The MaSIF-search network receives as input the features of a patch from one of these pairs (the ‘binder’), the inverted input features of its interacting patch (the ‘target’), and a patch randomly chosen from a different interface in the training set (the ‘random’ patch) (Supplementary Fig. S1). The neural network was trained on a Siamese neural network architecture to produce fingerprints that are similar for the ‘binder’ and ‘target’ patches while at the same time being dissimilar between ‘target’ and ‘random’. Similarity and dissimilarity was measured as the Euclidean distance between the fingerprints. A total of 85,652 true interacting pair patches and 85,652 noninteracting pair patches were used for training/validation, and 12,678 true interacting and 12,678 noninteracting pairs were used for the testing set.

Each of the 5 input features was computed in a separate channel consisting of a MaSIF geometric deep learning convolutional layer. Then the output from all channels was concatenated, and a Fully Connected Layer was used to output a fingerprint of size 80. In each batch, 32 pairs of interacting patches and 32 pairs of non-interacting patches were used. Adam was used as the optimizer, and a learning rate of 10^-3^ was used. The d-prime cost function^59^ was used as the loss function. MaSIF-search was trained for 40 hours in a GPU, after which it was automatically killed, resulting in 260,000 iterations of the data. The MaSIF-search neural network implementation contained a total of 66080 trainable parameters and was implemented in Tensorflow.

### Patch alignment and IPA scoring neural network

In the MaSIF-search pipeline, surfaces are computed for each protein of interest, and both a MaSIF-search fingerprint and a MaSIF-site prediction are computed for each surface vertex. All fingerprints within a user defined threshold for similarity to a target patch (defined at 1.7 by default) are then selected for a second-stage alignment and rescoring. In this step, the patch is extracted from the source protein, along with all the fingerprints for all vertices in the patch (since they were all precomputed). The random sample consensus (RANSAC) algorithm implemented in Open3D^60^ then uses the fingerprints of all the vertices in the target and matched patch to find an alignment between the patches. The RANSAC algorithm chooses three random points in the binder patch and computes the Euclidean distance of the surface MaSIF-search fingerprints between these points and all those points in the target patch; the most similar fingerprints provide the RANSAC algorithm with 3 correspondences to compute a transformation between the patches.

Once a candidate patch is aligned, the interface post-alignment (IPA) neural network (NN) is used to score the alignment with a score between 0 and 1 on the prediction of whether the alignment corresponds to a real interaction or not. Upon patch alignment, each vertex in the candidate patch is matched to the closest vertex in the target patch, and three features are computed per pair of vertices: (i) 1/(distance), the euclidean distance in 3D between the vertices; (ii) the product of the normal between the vertices, and (iii) 1/(fingerprint distance), the euclidean distance between the MaSIF-search fingerprints between the two vertices. A fourth feature, which we call ‘penetration’ is computed by computing the distance between each of the vertices in the candidate patch and all the atoms in the target. Thus, the IPA NN receives as input a vector of size Nx4, where N is the number of vertices in the candidate patch (up to 200 vertices). The IPA NN consists of 5 layers of 1D convolution, followed by a Global Averaging Pool layer and 7 fully connected layers. The 5 layers of 1D convolution contain 16, 32, 64, 128, and 256 filters, respectively, with a kernel size of 1 and a stride of 1, and each layer was followed by a batch normalization layer and a Rectified Linear Unit layer. The fully connected layers contained 128, 64, 32, 16, 8, 4, and 2 dimensions. Each fully connected layer was also followed by a Rectified Linear Unit layer, with the exception of the last layer which was followed by a softmax layer. The network was optimized with Adam^57^, with a learning rate of 10^-4^ and a categorical cross entropy loss function.

The IPA NN was trained as follows. The same dataset used for MaSIF-search, containing 4944 PPIs and a testing set of 957 PPIs was used. For each protein pair, one protein was chosen as the target, and the patch at the center of the interface was selected as the target patch. Then the partner protein along with 10 randomly chosen other proteins were aligned to it. Any alignment of the true partner within 3 Å RMSD of the co-crystal structure was considered as a positive. Any alignment from the true partner at greater than that RMSD or of any other protein was considered as a negative. Features were computed for all alignments and used for the IPA NN training. The IPA NN was trained with batches of 32 for 50 epochs.

### Assembly of surface fingerprints database for selection of binding seeds

#### Decomposition of the PDB into alpha helical peptides

A snapshot of the non-redundant set of the PDB was downloaded and decomposed into alpha helices, removing all non-helical elements. The DSSP program^61^ was used to label each residue according to their secondary structure. Fragments with 10 or more consecutive residues with a helical (‘H’) label assigned by DSSP were extracted. Each extracted helical fragment was treated as a monomeric protein, and surface features were computed for each one. MaSIF-search fingerprints and MaSIF-site labels were then computed for all extracted helices. MaSIF-seed uses both fingerprint similarity and interface propensity to identify suitable seeds. Ultimately, our binding seed database was composed of approximately 250 K helical motifs from which 140 M fingerprints were extracted.

#### Retrieval of beta-strand motifs

To collect beta-strand motifs, a snapshot of the non-redundant set of the PDB was preprocessed with the MASTER software^62^ to allow for fast structural matches. Two template motifs, one consisting of two beta strands and one consisting of three beta strands, were deprived of loops and served as input to MASTER to find sets of structurally similar motifs that would ultimately become the motif dataset for MaSIF. The search allowed for a variable backbone length of 1-10 amino acids connecting the beta strands of the template. RMSD cutoffs were set at 2.1 Å and 3 Å for two-stranded and three-stranded beta sheets, respectively. Similar to the preparation of helical motifs, each beta fragment was treated as a monomeric protein and surface features were generated, followed by the generation of MaSIF-search fingerprints and MaSIF-site labels. Ultimately, our beta-strand binding seed database was composed of approximately 390 K motifs from which 260 M fingerprints were extracted.

### MaSIF-seed - a pipeline to identify binding seeds to target novel interfaces

Based on the different modules within the MaSIF framework^17^, we developed a novel pipeline to identify potential binding seeds to targets. For each target, first MaSIF-seed was used to label each point in the surface for the propensity to form a buried surface region. Then, a fingerprint was computed for the target site. Finally, after scanning the entire protein, the best patch was selected. In one case, the SARS-CoV-2 RBD, the fourth best site was selected as it was the site with the highest potential to disrupt binding to the natural receptor. Then, a MaSIF-search fingerprint was computed for the target patch, inverting the target features before inputting them to the MaSIF-search network. The Euclidean distances between the target fingerprint and the millions of fingerprints in the binding seed database were then computed, and all patches with a fingerprint distance below defined thresholds were accepted. In this paper the thresholds utilized were <2.0 for PD-L1, PD-1 and CTLA-4, and <1.7 for the RBD.

Once fingerprints are matched, a second-stage alignment and scoring method uses the RANSAC algorithm as described above. After RANSAC produces an alignment, the IPA neural network classifies true binders vs. non-binders^17^ and outputs an IPA score (described above). Those candidate binders with an IPA score of more than 0.90-0.97 in the neural network score were accepted.

### Computational benchmark for binding motif recovery

#### Helix:receptor motifs

A set of transient interactions from PDBBind was scanned to identify proteins that bind to helical motifs. A binding motif was determined to be a helix if 80% of residues are helical and the total number of residues does not exceed 60. The selected complexes were filtered to remove pairs of PPIs with high homology and a set of 31 unique PPIs was used, subsequently MaSIF-search fingerprints and MaSIF-site fingerprints were computed. MaSIF-seed was benchmarked against a hybrid pipeline of existing, fast, well-established docking tools on the dataset of helix:receptor proteins: PatchDock^63^, ZDock^22, 64^, and ZRank2^23^. For each helix:receptor pair, the helix from the co-crystal structure was placed along 1000 randomly selected helices from the motif database. Then the methods were benchmarked to evaluate their capacity to rank the correct helix from the co-crystal structure, with an alignment RMSD <3.0 Å from the conformation of the co-crystal structure, versus the remaining 1000 helices. We note that each helix can potentially bind in many possible orientations, and in the case of methods that were not preceded by a MaSIF-site identification of the target site, the helix can bind on many sites on the receptor. The measured time for all methods included only the scoring time, except for MaSIF-seed where the alignment time was also included in the calculation. **MaSIF-seed -** All of MaSIF-seed’s neural networks (MaSIF-search, MaSIF-site and the IPA score) were retrained for this benchmark to remove helix:receptor pairs from the training set. In each case, MaSIF-site was used to identify the patch in the target protein with the highest interface propensity, and the fingerprint for the selected patch was compared to the fingerprints of all patches in the database. The rigid orientation of each helix in the benchmark was randomly rotated and translated prior to any alignment. Patches were discarded if their MaSIF-search fingerprint’s euclidean distance to that of the target site was greater than 1.7. After alignment, patches were further filtered if the IPA score was less than 0.96. **PatchDock+MaSIF-site -** On each receptor protein, MaSIF-site was used to identify and label the target site, while PatchDock^63^ was used to dock all 1001 helices, setting the target site based on a specific residue using the ReceptorActiveSite flag in PatchDock. The PatchDock score was used to produce the ranking of all conformations for all 1001 helices. **ZDock -** Was run on standard parameters and its standard scoring was used similar to PatchDock. In the ZDock+MaSIF-site case, all residues outside of the MaSIF-site selected patch were blocked using the compute_blocked_res_list.sh provided in ZDock. **ZDock+ZRank2** - In this variant, the top 2000 results from ZDock with each of the 1001 peptides for each of the 31 receptors were re-scored using ZRank2. The ZRank2 score was then used to score all of the docking poses.

### Non-helix:receptor motifs

The same set of transient interactions from PDBBind was filtered for proteins interacting through non-helical motifs. The secondary structure types of the proteins were annotated with DSSP^61^, followed by computing the contribution of helical segments (DSSP annotation of H, G, or I) to the interface. Only interfaces with less than 50% helical segments were selected. Additional filtering was performed by requiring a mean shape complementary at the interface of >0.55 and a maximum inscribed patch area of >150 Å^2^. From these native complexes, seeds were extracted by selecting residues within 4 Å distance to the receptor and extending the backbone of these residues on their N- and C-terminus until the DSSP annotation changed to capture complete secondary structure elements. In total 83 complexes were collected for the benchmark.

The decoy set was constructed from 1000 randomly selected beta-strand seeds from the MaSIF-seed pipeline, containing 500 two-stranded and 500 three-stranded beta motifs. The benchmark was performed similarly to the helix:receptor benchmark described above with adapting the fingerprint’s euclidean distance cutoff to a value of 2.5 and allowing MaSIF-seed to evaluate the top two sites in each receptor. These modifications were performed for this benchmark as it increased the accuracy while still performing at least 20 times faster than comparable competing tools. Only ZDock and ZDock/ZRank2 were benchmarked in the non-helical benchmark as ZDock/ZRank2 was shown to be the best in the helical benchmark.

### Clustering of seed solutions

In each design case all of the top matched seeds were clustered by first computing the root mean square deviation between all pairwise helices, computed on the C-alpha atoms of each pair of helices, in the segment overlapping over the buried surface area. The pairwise distances were then clustered using metric multidimensional scaling^65^ implemented in scikit-learn^47^.

### Seed and interface refinement

For the “one-shot” protocol, seed candidates proposed by MaSIF were refined using Rosetta and a FastDesign protocol with a penalty for buried unsatisfied polar atoms in the scoring function^30^. Beta sheet-based seeds containing >33% contact residues found in loop regions were discarded. 33, 200, and 109 refined seeds were selected based on the computed binding energy, shape complementarity, number of hydrogen bonds and counts of buried unsatisfied polar atoms for PD-1, CTLA-4, and PD-L1, respectively.

### Grafting of seeds onto monomeric scaffolds and computational design with Rosetta

A representative seed was selected from each solution space, and then matched using Rosetta MotifGraft to a database of 1300 monomeric scaffolds in the case of the RBD and PD-L1 designs. For the optimized protocol selected seeds were grafted to a database of 4,347 small globular proteins (<100 amino acids), originating from the PDB^66^, two computationally designed miniprotein databases^31, 32^ and one AF2 proteome prediction database^9, 67^. Seeds were cropped to the minimum number of side chains making contact before grafting. Moreover, loop regions from beta sheet-based seeds were completely removed. After side-chain grafting by Rosetta, a computational design protocol was used to design the remaining interface. Final designs were selected for experimental characterization based on the computed Rosetta binding energy, the shape complementarity, number of hydrogen bonds and counts of buried unsatisfied polar atoms.

### Yeast surface display of single designs

DNA sequences of designs were purchased from Twist Bioscience containing homology overhangs for cloning. DNA was transformed with linearized pCTcon2 (Addgene #41843) or a modified pNTA vector with V5 tag into EBY-100 yeast using the Frozen-EZ Yeast Transformation II Kit (Zymo Research). Transformed yeast were passaged once in minimal glucose medium (SDCAA) before induction of surface display in minimal galactose medium (SGCAA) overnight at 30°C. Transformed cells were washed in cold PBS with 0.05-0.1% BSA and incubated with the binding target for 2 hours at 4°C. Cells were washed once and incubated for an additional 30 minutes with appropriate antibodies (Supplementary Table 5). Cells were washed and analyzed using a Gallios flow cytometer (Beckman Coulter). For quantitative binding measurements, binding was quantified by measuring the fluorescence of a PE-conjugated anti-human Fc antibody (Invitrogen) detecting the Fc-fused protein target. Yeast cells were gated for the displaying population only (V5 positive) (Supplementary Fig. 9a).

### Yeast libraries

Combinatorial sequence libraries were constructed by assembling multiple overlapping primers (Supplementary Table 6) containing degenerate codons at selected positions for combinatorial sampling of the binding interface, core residues or hydrophobic surface residues. Primers were mixed (10 µM each) and assembled in a PCR reaction (55 °C annealing for 30 sec, 72 °C extension time for 1 min, 25 cycles). To amplify full-length assembled products, a second PCR reaction was performed, with forward and reverse primers specific for the full-length product. For SSM libraries and oligo pools, DNA was ordered from Twist Biosciences and amplified with primers to give homology to the pCTcon2/pNTA backbone. In all cases, the PCR product was desalted and used for transformation.

### Yeast surface display of libraries

Combinatorial libraries, SSM libraries, and oligo pools were transformed as linear DNA fragments in a 5:1 ratio with linearized pCTcon2 or pNTA_V5 vector as described previously into EBY-100 yeast^68^. Transformation efficiency generally yielded around 10^7^ transformants per cuvette. Transformed yeast were passaged at least once in minimal glucose medium (SDCAA) before induction of surface display in minimal galactose medium (SGCAA) overnight at 30°C. Induced cells were labeled in the same manner as the single designs. Labeled cells were washed and sorted on a Sony SH800 cell sorter. For combinatorial libraries and oligo pool libraries, sorted cells were grown in SDCAA and prepared similarly for two additional rounds of sorting. After the third sort cells were plated on SDCAA agar and single colonies were sequenced. SSM libraries were sorted once, collecting both binding and nonbinding populations, and grown in liquid culture for plasmid preparation.

### MiSeq Sequencing

After sorting, yeast cells were grown in SDCAA medium, pelleted and plasmid DNA was extracted using Zymoprep Yeast Plasmid Miniprep II (Zymo Research) following the manufacturer’s instructions. The coding sequence of the designed variants was amplified using vector-specific primer pairs, Illumina sequencing adapters and Nextera barcodes were attached using an additional overhang PCR, and PCR products were desalted with Qiaquick PCR purification kit (Qiagen) or AMPure XP selection beads (Beckman Coulter). Next generation sequencing was performed using Illumina MiSeq with appropriate read length, yielding between 0.45-0.58 million reads/sample. For bioinformatic analysis, sequences were translated in the correct reading frame, and enrichment values were computed for each sequence.

### Protein expression and purification

DNA sequences were ordered from Twist Bioscience and Gibson cloning or T7 ligation used to clone into bacterial (pET21b) or mammalian (pHLSec) expression vectors. Protein binder and target constructs are listed in Supplementary Table 2 and 7 respectively. Mammalian expressions were performed using the Expi293^TM^ expression system from Thermo Fisher Scientific. Supernatant was collected 6 days post transfection, filtered, and purified. *E. coli* expressions were performed using BL21 (DE3) cells and IPTG induction (1 mM at OD 0.6-0.8) and growth overnight at 16-18° C. Pellets were lysed in lysis buffer (50 mM Tris, pH 7.5, 500 mM NaCl, 5% Glycerol, 1 mg/ml lysozyme, 1 mM PMSF, and 1 µg/ml DNase) with sonication, the lysate clarified, and purified. All proteins were purified using an ÄKTA pure system (GE healthcare) with either Ni-NTA affinity or protein A affinity columns followed by size exclusion chromatography. If TEV cleavage was necessary, fused proteins were dialyzed overnight at 4°C (dialysis buffer 20 mM Tris pH 7.5, 150 mM NaCl, 10% glycerol) with excess TEV enzymes.

### Surface plasmon resonance

SPR measurements were performed on a Biacore 8K (GE Healthcare) with HBS-EP+ as running buffer (10 mM HEPES pH 7.4, 150 mM NaCl, 3 mM EDTA, 0.005% v/v Surfactant P20, GE Healthcare). Ligands were immobilized on a CM5 chip (GE Healthcare # 29104988) via amine coupling. 500-1000 response units (RU) were immobilized and designed proteins were injected as an analyte in serial dilutions. The flow rate was 30 µl/min for a contact time of 120 s followed by 800 s dissociation time. After each injection, the surface was regenerated using 3 M magnesium chloride (for PD-L1) or 10 mM glycine, pH 3.0 (for RBD). Data were fit with 1:1 Langmuir binding model within the Biacore 8K analysis software (GE Healthcare #29310604).

### Biolayer Interferometry

BLI measurements were performed on a Gator BLI system. The running buffer was 150 mM NaCl, 10 mM HEPES pH 7.5. Fc-tagged designs were diluted to 5 ug/mL and immobilized on the tips (1-2 nm immobilized). The loaded tips were then dipped into serial dilutions of either spike protein or RBD. Curves were fit using a 1:1 model on the Gator software after subtracting the background.

### Size exclusion chromatography multi-angle light scattering (SEC-MALS)

Size exclusion chromatography with an online multi-angle light scattering device (miniDAWN TREOS, Wyatt) was used to determine the oligomeric state and molecular weight for the protein in solution. Purified proteins were concentrated to 1 mg/ml in PBS (pH 7.4), and 100 µl of the sample was injected into a Superdex 75 300/10 GL column (GE Healthcare) with a flow rate of 0.5 ml/min, and UV_280_ and light scattering signals were recorded. Molecular weight was determined using the ASTRA software (version 6.1, Wyatt).

### Circular Dichroism

Far-UV circular dichroism spectra were measured using a Chirascan™ spectrometer (AppliedPhotophysics) in a 1-mm path-length cuvette. The protein samples were prepared in a 10 mM sodium phosphate buffer at a protein concentration between 20 and 50 µM. Wavelengths between 200 nm and 250 nm were recorded with a scanning speed of 20 nm min^−1^ and a response time of 0.125 secs. All spectra were averaged two times and corrected for buffer absorption. Temperature ramping melts were performed from 20 to 90 °C with an increment of 2 °C/min. Thermal denaturation curves were plotted by the change of ellipticity at the global curve minimum to calculate the melting temperature (T_m_).

### Cell binding analysis

For flow cytometry analysis of DBL1 designs binding to PD-L1 on Karpas-299 cells, 2×10^5^ cells were incubated with 50 µL Fc Block (BD Biosciences, cat #553142) that was pre-diluted 1:50 in FACS buffer (PBS (Gibco/Thermofisher scientific, cat #10010-015) and 2% BSA (Sigma Aldrich, cat #A7906)) for 15 minutes on ice. Samples were subsequently supplemented with 50 µL of PD-L1 binders prepared as follows: high-affinity PD-1_Fc serially diluted 1:2 for 20 dilutions in FACS buffer, starting at 62.5 µg/ml; DBL1_03_Fc and DBL1_04_Fc serially diluted 1:2 for 16 dilutions in FACS buffer, starting at 125 µg/ml; DBL1_03_KO_Fc and PD-1_Fc serially diluted 1:2 for 14 dilutions in FACS buffer, starting at 125 µg/ml. The cell solutions were incubated for 30 minutes. Samples were then washed three times, resuspended in 100 µL of FACS buffer containing secondary R-PE Goat Anti-Human IgG antibody diluted 1:100 (Jackson ImmunoResearch, cat #109-117-008), and incubated for 30 minutes. Samples were then washed three times to remove unbound antibody, resuspended in 100 µL of FACS buffer, and analyzed using LSR Fortessa flow cytometer (BD Biosciences).

### Purification of proteins for crystallography

PD-L1 extracellular domain fragment (from F19 to R238) was over-expressed as inclusion bodies in the BL21 (DE3) strain of *E. coli.* Renaturation and purification of PD-L1 was performed as previously described^69^. Briefly, inclusion bodies of PD-L1 was diluted against a refolding buffer (100 mM Tris, pH 8.0; 400 mM L-Arginine; 5 mM EDTA-Na; 5 mM Glutathione (GSH); 0.5 mM Glutathione disulfide (GSSG)) at 4°C for 24 h. Then the PD-L1 was concentrated and exchanged into a buffer of 20 mM Tris-HCl (pH 8.0) and 15 mM NaCl and further analyzed by HiLoad 16/60 Superdex 75 pg (Cytiva) chromatography. PD-L1 binder designs, DBL1_03 and DBL2_02, were over expressed in *E. coli* as inclusion bodies. Renaturation and purification of the PD-L1 binder designs was performed as the PD-L1 protein. PD-L1 and binder designs were then mixed together at a molar ratio of 1:2 and incubated for 1h on ice. The binder/PD-L1 complex was further purified by HiLoad 16/60 Superdex 75 pg (Cytiva) chromatography.

### Data collection and structure determination

For crystal screening, 1 μl of binder/PD-L1 complex protein solution (10 mg/mL) was mixed with 1 μl of crystal growing reservoir solution. The resulting mixture was sealed and equilibrated against 100 μl of reservoir solution at 4° or 18°C. Crystals of the DBL1_03/PD-L1 complex were grown in 0.2 M potassium formate and 20% w/v PEG 3350. Crystals of the DBL2_02/PD-L1 complex were grown in 0.2 M potassium/sodium tartrate, 0.1 M Bis Tris propane, pH 6.5 and 20 % w/v PEG 3350. Crystals were flash-cooled in liquid nitrogen after incubating in anti-freezing buffer (reservoir solution containing 20% (v/v) glycerol). Diffraction data of crystals were collected at Shanghai Synchrotron Radiation Facility (SSRF) BL19U. The collected intensities were subsequently processed and scaled using the Denzo program and the HKL2000 software package (HKL Research). The structures were determined using molecular replacement with the program Phaser MR in CCP4, with the reported PD-L1 structure (PDB: 3RRQ) as the search model^70^. COOT and PHENIX were used for subsequent model building and refinement^71, 72^. The stereochemical qualities of the final model were assessed with MolProbity^73^. Data collection details and refinement statistics are in Supplementary Table 8.

### Luminex binding assays

Luminex beads were prepared as previously published^27^. Briefly, MagPlex beads were covalently coupled to SARS-CoV-2 spike proteins of different variants. The serial dilutions of the antibodies or design were performed and binding curves were fit using GraphPad Prism nonlinear four parameter curve fitting analysis of the log(agonist) versus response.

### Live virus neutralization assays

The virus neutralization assays were performed as previously published^27^. Briefly, VeroE6 cells were seeded in 96 well plates the day before the infection. The DBR3_03-Fc compound in serial dilutions was mixed with omicron-spike virus and incubated at 37°C for one hour before addition to the cells. The cells with virus were kept a further 48 hours at 37°C, then washed and fixed for crystal violet staining and analysis. Neutralization EC_50_ calculations were performed using GraphPad Prism nonlinear four parameter curve fitting analysis.

### Cryo-EM sample preparation and data acquisition

For cryo-electron microscopy investigations, 3.0 µl aliquots at a concentration of 0.87 or 1.0 mg/ml of the spike_D614G_-binder sample or the spike_Omicron_-binder sample were applied onto glow-discharged carbon-coated copper grids (Quantifoil R2/1, 400 mesh), blotted for 4.0-8.0 s, and flash-frozen in a liquid ethane/propane mixture cooled to liquid nitrogen temperature, using Vitrobot Mark IV (Thermo Fisher Scientific) with 100% humidity and the sample chamber operated at 4 °C. Grids were screened in a Thermo Fisher Scientific (TFS) 200kV Glacios cryo-EM instrument. Suitable grids were transferred to TFS Titan Krios instruments for data collection. Cryo-EM data-collection statistics of this study are summarized in Supplementary Table 9. The spike_D614G_-binder data composed of 20,794 movies was collected on a Titan Krios G4 microscope, equipped with a cold-FEG electron source and operated at 300 kV acceleration voltage. Movies were recorded with the automation program EPU (TFS) on a Falcon4 direct electron detector in counting mode at a physical pixel size of 0.40 Å per pixel and a defocus ranging from −0.8 to −2.0 μm. Exposures were collected as electron event recordings (EER) with a total dose of 80 e^−^/Å^2^ over approximately 3 seconds, corresponding to a dose rate of 4.53 e^-^/px/s. For spike_Omicron_-binder data, 22,266 movies were recorded on a Titan Krios G4 microscope, equipped with TFS SelectrisX imaging filter and Falcon4 camera. Exposures were collected at 60 e^−^/Å^2^ total dose with a physical pixel size of 0.726 Å per pixel over approximately 6 seconds, corresponding to a dose rate of 5.4 e^-^/px/s, at a defocus range of −0.8 to −2.5 μm. Data was analyzed by cryoSPARC v3.3.1^74^.

### Cryo-EM image processing, model building and refinement

Details of the image processing are shown in Supplementary Figures S13-15, S17-19 and Supplementary Table 9. Recorded movies in EER format were imported into cryoSPARC v3.3.1^74^ and gain-normalized, motion-corrected and dose-weighted using the cryoSPARC implementation of patch-based motion correction. CTF estimation was performed using the patch-based option in cryoSPARC. A small set of particles were manually selected and followed by 2D classification to create a 2D template for the subsequent automatic particle picking. For the sample of spike_D614G_ in complex with the de-novo designed binder, 832,816 particles were automatically selected by template-based picker and subjected to three rounds of 2D classifications, resulting in a particle set of 184,763 particles. The particles were grouped into three classes, using the ab-initio and hetero-refine implementations in cryoSPARC. The best 3D class composed of 97,804 particles was further subjected to another round of ab-initio reconstruction and hetero-refinement. The well-resolved class consisting of 67 432 particles resulted in a 2.6 Å overall resolution global map in C1 symmetry. The binder-RBD region was refined with a soft mask, resulting in a local map at 3.1 Å resolution. For the data processing of the spike_Omicron_-binder complex sample, 1,820,333 particles were picked with the cryoSPARC template-based picker. After two rounds of 2D classifications, 981,561 particles were selected and subjected to ab-initio reconstruction and hetero-refinement, resulting in a set of 595,599 particles. Subsequently, the selected particle set was classified by multiple rounds of 3D classifications in cryoSPARC. The best-resolved 3D class containing 50,758 particles resulted in a 2.8 Å overall resolution map and the binder-RBD region was further improved by performing focused refinement with a soft mask, resulting in a map at 3.3 Å resolution. Resolution for all 3D maps was estimated based on the Fourier shell correlation (FSC) with a cutoff value of 0.143.

For model building of the spike_D614G_-binder, the previous model (PDB: 7BNO, spike_D614G_) was used for the region of spike_D614G_ as a starting model. The model was rigid-body fit into the cryo-EM density in UCSF Chimera^75^ and adjusted manually in Coot 0.9.4^76^. *De novo* building for the binder parts was performed manually in Coot 0.9.4. For building the spike_Omicron_-binder structure, the model (PDB: 7QO7, spike_Omicron_) was fitted into the density and rebuilt and adjusted manually, using UCSF Chimera and Coot 0.9.4. After the structural rebuilding, all the atomic models were refined using the Phenix (1.19.2-4158) implementation of real.space.refine with general structural restraints^77, 78^. Comprehensive validation (cryo-EM), model quality assessment and statistics are in Supplementary Table 9. EM densities and atomic models were visualized in UCSF Chimera, UCSF ChimeraX^79^ and Pymol (http://www.pymol.org/).

## Data availability

Raw cryo-EM data is being deposited in the EMPIAR databank. Cryo-EM maps were deposited in the Electron Microscopy Data Bank under the access codes of EMD-14947 (spike_D614G_-binder full and spike_D614G_-binder local maps), EMD-14922 (spike_Omicron_-binder full), and EMD-14930 (spike_Omicron_-binder local). Atomic models were deposited in Protein Data Bank under the access codes of PDB-7ZSS (spike_D614G_-binder), PDB-7ZRV (spike_Omicron_-binder full) and PDB-7ZSD (spike_Omicron_-binder local). Crystal structures have been deposited in the Protein Data Bank under accession codes 7XYQ (DBL1_03/PD-L1 complex) and 7XAD (DBL2_02/PD-L1 complex). MaSIF-seed and the Rosetta design scripts are available at https://github.com/LPDI-EPFL/masif_seed.

## Author contributions

P.G., S.W., A.V., A.M., A.S., M.B. and B.E.C. conceived the work and designed the experiments. P.G., A.M., A.S. and Z.H. performed the computational design and S.W., A.V., S.B. and A.M. performed experimental characterization and optimization. P.G., F.S., A.M., Z.H., and A.S. developed the MaSIF-seed method. D.N. and H.S. solved the cryo-EM structure. S.T., M.P., K.L., Z.X., Y.C., P.H. and G.F.G solved the crystal structures. A.P. and E.O. performed the PD-L1 cell binding assay. A.T. and B.F. synthesized peptides. P.T., C.R., and D.T. performed SARS-CoV-2 binding and neutralization studies. F.S., C.G., S.R., S.G. and J.M. performed experiments and acquired data. P.G., S.W., A.V., A.M., A.S. and B.E.C. wrote the manuscript with input from all authors.

## Acknowledgements

We thank the Dubochet Center for Imaging (DCI) in Lausanne for cryo-EM data collection. The DCI is an initiative of the EPFL, University of Lausanne and University of Geneva. DN and HS were supported by the Swiss National Science Foundation and the NCCR Transcure. MB was supported by an ERC Consolidator grant No. 724228. BC was supported by the Swiss National Science Foundation, the NCCR in Chemical Biology, the NCCR in Molecular Systems Engineering and the ERC Starting grant no. 716058. P.G. was sponsored by an EPFL-Fellows grant funded by an H2020 Marie Sklodowska-Curie. We thank K. Lau and F. Pojer from the PTPSP facility at EPFL for providing SARS-CoV-2 spike proteins and assistance with cryo-EM. We thank SCITAS at EPFL for support in the computational simulations. We thank the GECF for assistance in deep sequencing and FCCS for assistance in FACS. We thank Emmanuel Levy, Sarel Fleishman, and Mihai Azoitei for their feedback on the manuscript.

