## Supplementary material for "*De novo* design of site-specific protein interactions with learned surface fingerprints": supp material

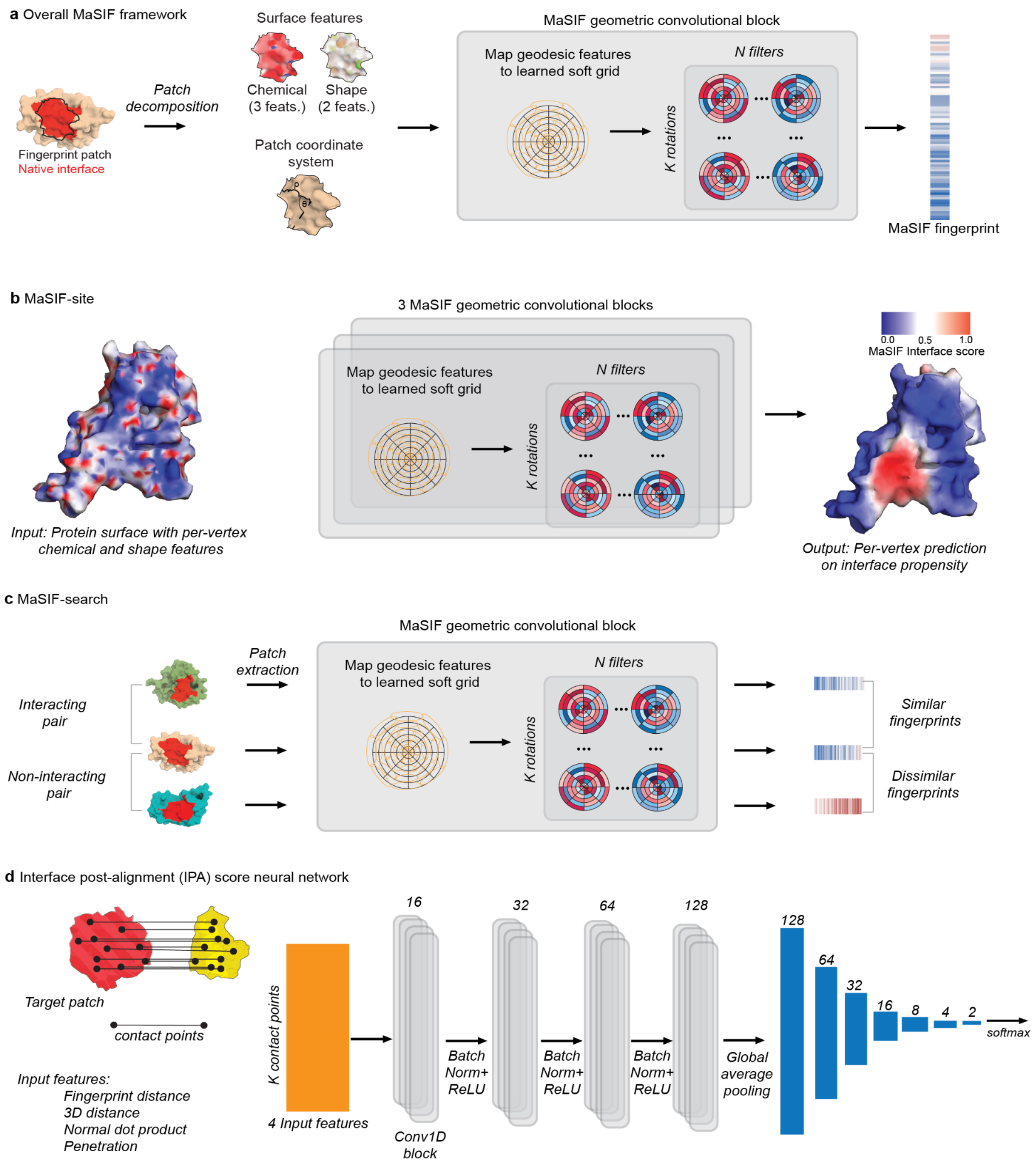

**Supplementary Figure S1: Overview of the neural network architectures used in the MaSIF protocols. a**, General MaSIF framework. Molecular surfaces are decomposed into patches which are annotated with chemical and shape features. The MaSIF network translates these input features into fingerprints that describe the original surface patch. **b**, MaSIF-site neural network. MaSIF-site predicts partner-independent protein interface propensities based on per-vertex chemical and shape features of the protein surface. **c**, MaSIF-search neural network. MaSIF-search embeds protein patches into a space where complementary patches are close to each other. The network was trained on discriminating interacting patches from non-interacting protein surface patches. The network uses MaSIF fingerprints to identify which are compatible and therefore to predict likely interacting proteins. **d,** Interface post-alignment (IPA) scoring neural network. The IPA scoring neural network enables the scoring of protein interfaces based on several input features: fingerprint distance between contacting points, 3D distance of corresponding points, normal dot product, and the distance between surface points in the seed and the closest atom in the target, which we call 'penetration'.

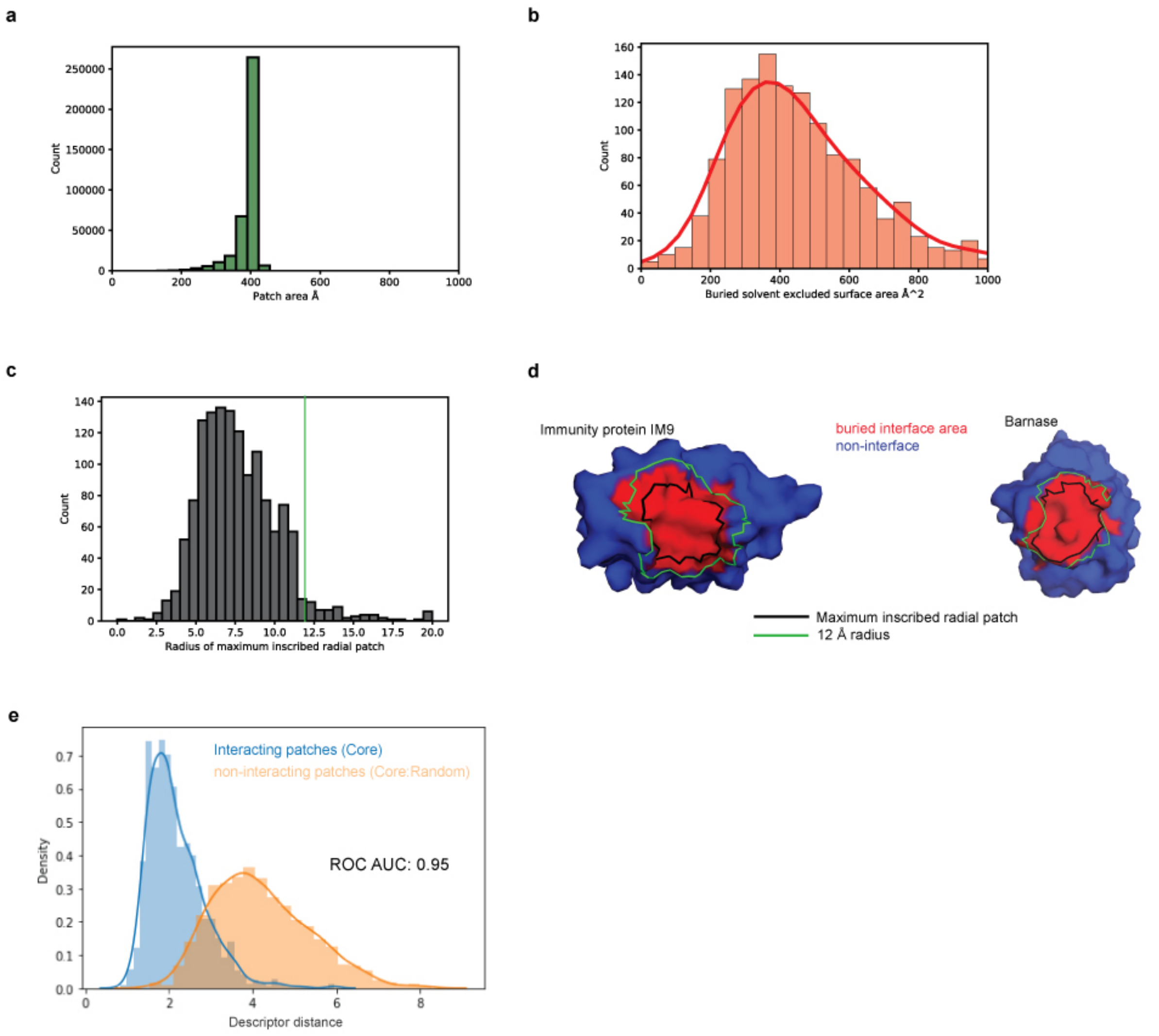

**Supplementary Figure S2: Modeling buried surfaces as radial patches. a,** Histogram of the patch areas of thousands of randomly selected protein patches with a fixed radius of 12 Å. **b,** Histogram of the area of the buried surface area on 1380 dimeric PPIs. We note that areas are computed for only one of the proteins (i.e. each subunit in a PPI is computed separately), and that we used the solvent excluded surface area, while other authors report buried areas on the solvent accessible area that include the buried surface area of both proteins (see methods). **c,** Size of the maximum inscribed radial patch for the 1380 proteins (see methods). Patch area for the radius used here (12 Å), using a set of 30,000 randomly selected patches. **d,** Example of the buried interface area for two well known, high affinity binders, Immunity Protein IM9 (PDB ID: 1EMV) and the protein Barnase (PDB ID: 1BRS). The buried interface of each protein when bound to its partner is shown in red. The maximum inscribed radial patch's circumference is shown in black, and the circumference of a patch with radius 12 Å is shown in green. **e,** Histogram of similarities between MaSIF-search fingerprint similarity between: (blue) pairs of patches that are co-crystallized from transient PPIs, with the fingerprint computed for the patch centered on the largest inscribed radial patch, and (orange) pairs of patches where one was taken from the center of the interface of a random PPI and the other was taken from a random patch surface.

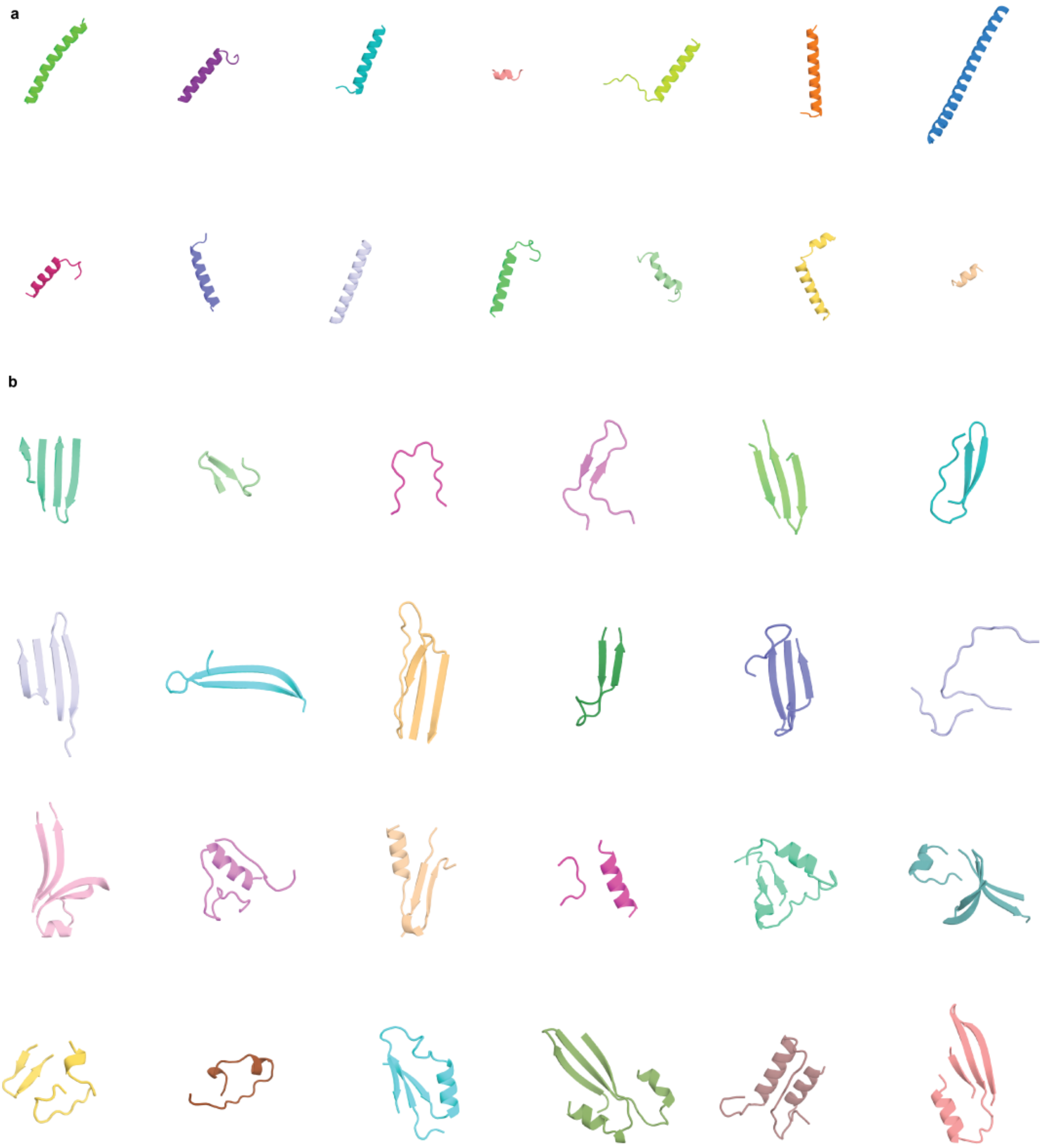

**Supplementary Figure S3: Overview of helical and non-helical seeds used in the recovery benchmark.** Examples of **a,** helical seed, **b,** non-helical seeds that were extracted for the recovery benchmark.

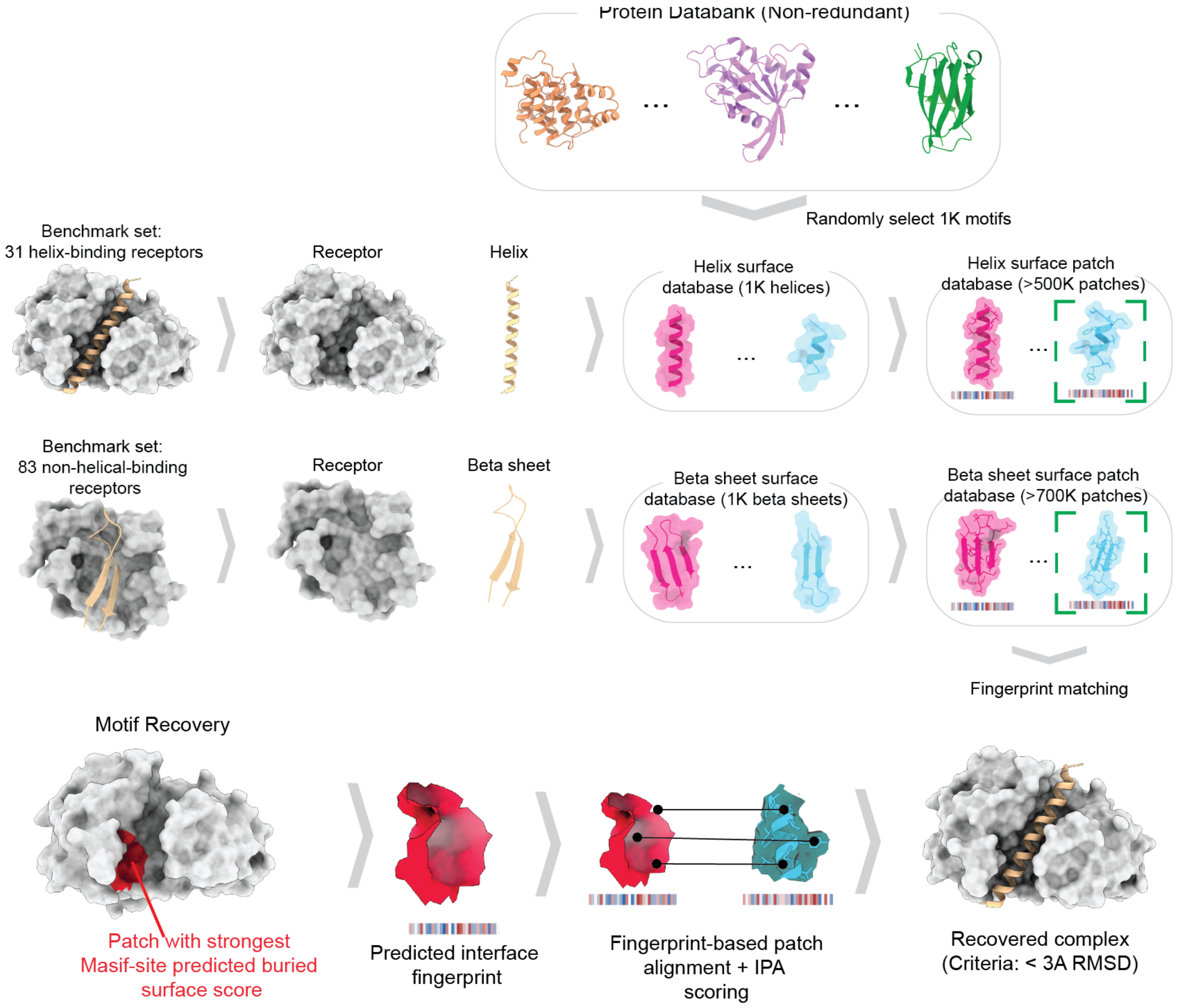

**Supplementary Figure S4: MaSIF-seed benchmarking for the discrimination of helical or non-helical binding motifs**. A non-redundant set of 31 helical and 83 non-helical fragments that bind to known protein receptors was selected as a benchmark set to evaluate MaSIF-seed's capacity to recover true binding motifs from decoys, and to correctly rank them among the top results. To generate the decoy set, a non-redundant set of all protein chains in the Protein Data Bank was decomposed into continuous helical segments (left) and two/three-stranded beta sheets (right), resulting in over 250K helical and over 380K beta motifs, respectively. One thousand of these motifs each were randomly selected to act as decoys in the respective benchmarks. The surfaces for the two sets of 1000 motifs were computed and decomposed into radial patches and for each patch a fingerprint was computed. Recovered complexes were considered correct if an iRMSD < 3 Å was obtained. A comparable procedure was applied to the benchmark tools.

a b

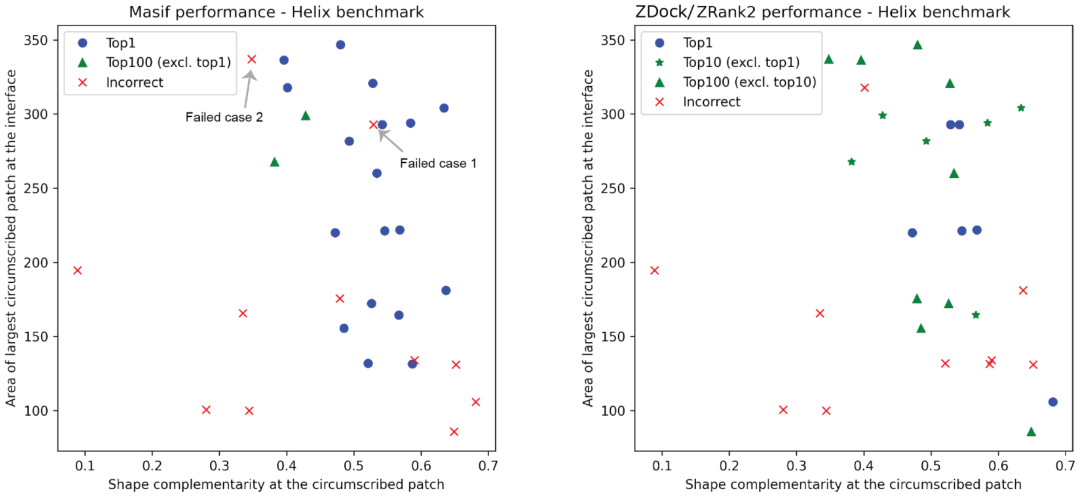

c

| | Benchmark cases solved by MaSIF-seed but not by ZDock/ZRank2 | Benchmark cases solved by ZDock/ZRank2 but not by MaSIF-seed | Benchmark cases solved by both MaSIF-seed and ZDock/ZRank2 |
|---|---|---|---|
| In the Top 1 | 14 | 2 | 4 |
| In the Top 10 | 10 | 4 | 8 |
| In the Top 100 | 4 | 5 | 16 |

d

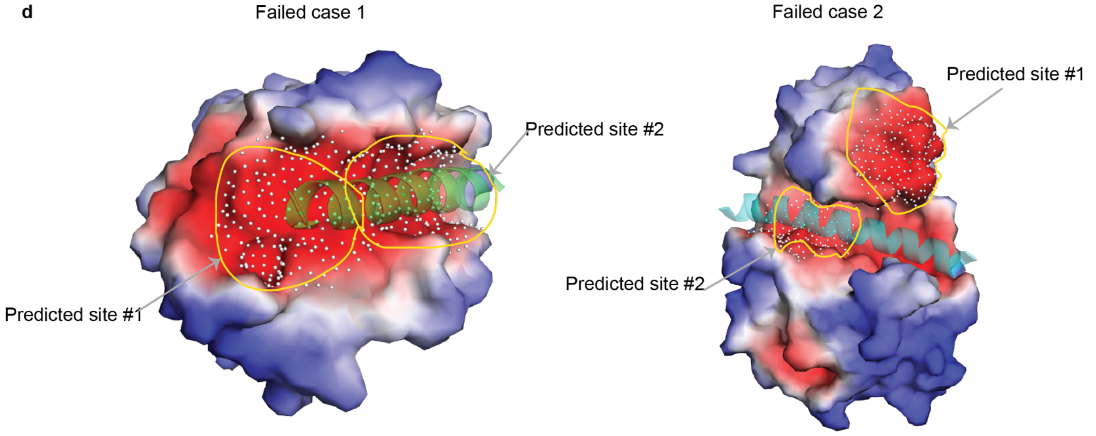

**Supplementary Figure S5: Analysis of successful/failed helical benchmark cases and comparison between MaSIF-seed and ZDock/ZRank2 performance. a-b,** Plotting of Top 1, Top 10, Top 100 and failed cases for MaSIF-seed and ZDock/ZRank2, showing the maximum circumscribed patch area in the buried interface (y-axis) and the median shape complementarity for vertices of that patch (x-axis) for **a,** MaSIF-seed, and **b**, ZDock/ZRank2. **c**, Comparison of cases solved by only MaSIF-seed, only ZDock/ZRank2, or both MaSIF-seed and ZDock/ZRank2 in the Top 1, top 10 or Top 100 rank. **d**, Analysis of two cases that showed both a large circumscribed patch and high complementarity at that patch where MaSIF-seed failed. In both cases, MaSIF-seed failed because it identified a different site as the top site, but increasing the number of sites explored to the top two resulted in successful predictions. The white dots on the surface denote the predicted site patches.

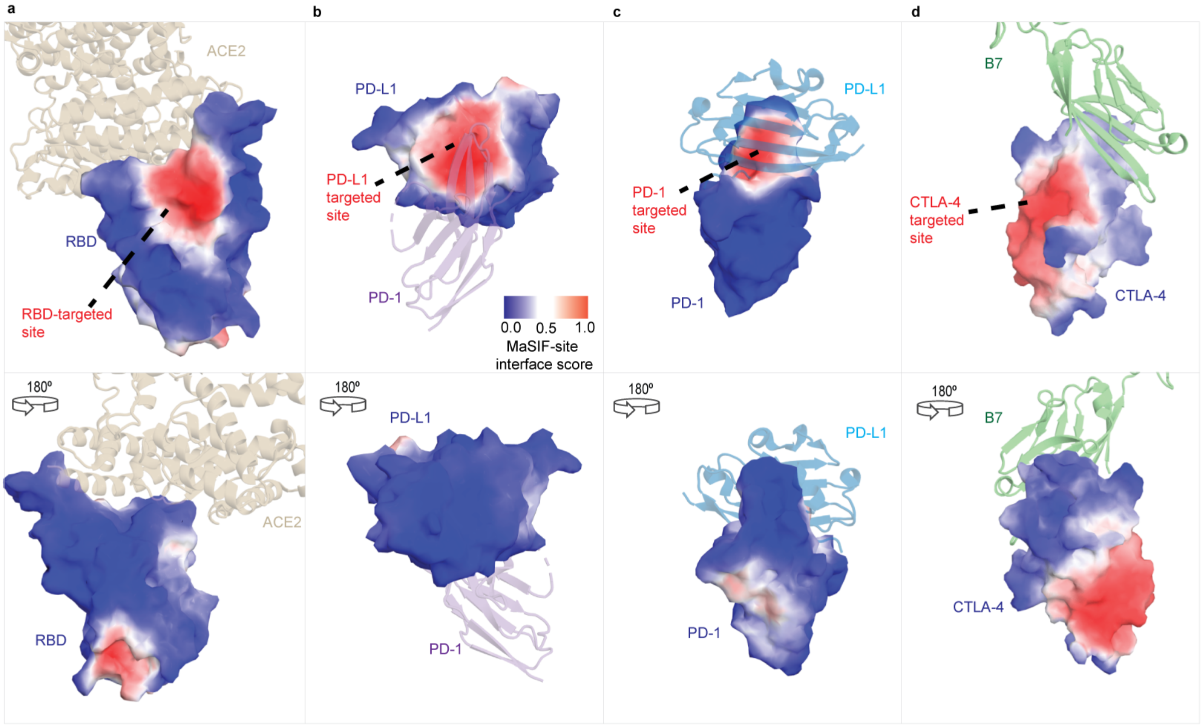

**Supplementary Figure S6: MaSIF-site target site prediction on SARS-CoV-2 RBD, PD-L1, PD-1, and CTLA-4.** Surface mode shows a MaSIF-site per-surface-vertex regression score on the propensity of each point on the surface to form an interface ranging from 0 (blue) to 1 (red) **a-c**, Predictions on each target, with the natural ligand of the target shown in cartoon representation as a reference. The structures highlight the predicted site and the bottom row shows a 180 degree rotation. **a**, MaSIF-site prediction on SARS-CoV-2 RBD (PDB ID: 6M17), with the RBD shown in surface and the ACE2 in beige. **b**, Prediction on PD-L1 (PDB ID: 5JDS), with PD-1 shown in purple. **c**, Prediction on PD-1 (PDB ID: 4ZQK) with the natural binder PD-L1 shown in cyan. **d**, Prediction on CTLA-4 (PDB ID: 5GGV) with the natural binding partner B7 (PDB ID: 1I8L) shown in light green.

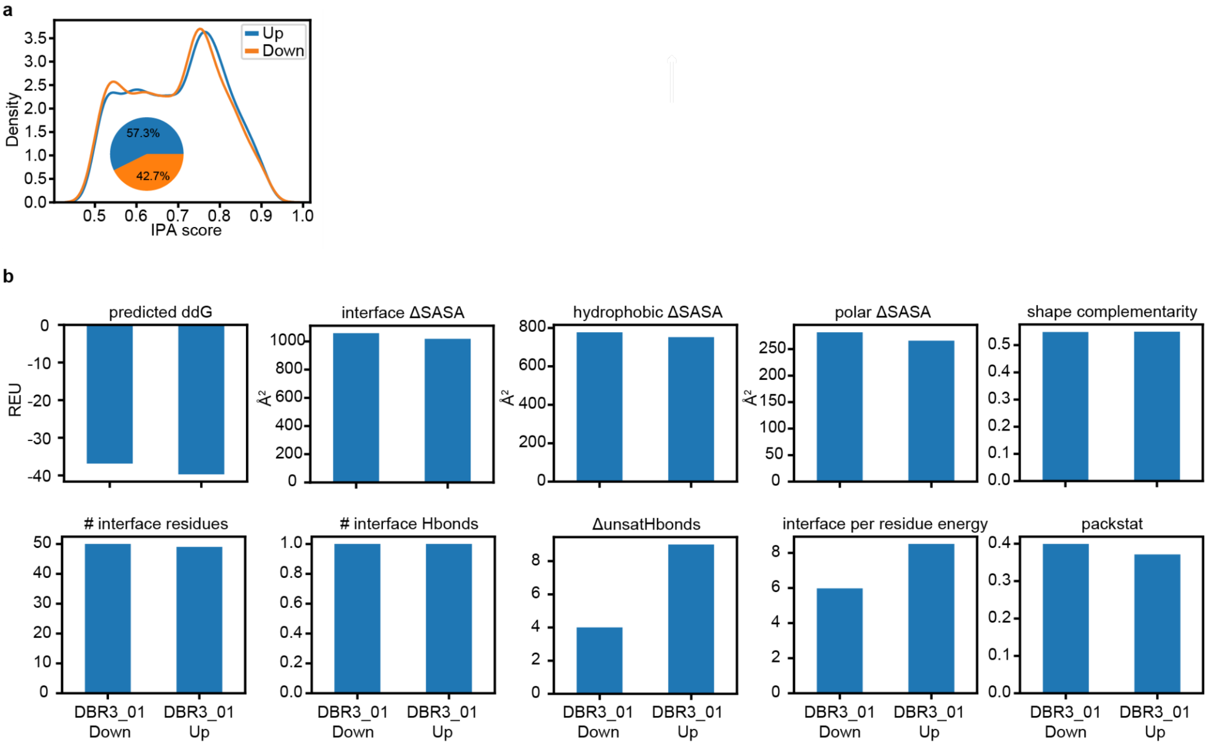

**Supplementary Figure S7: RBD-binder metrics for up- and down-orientations . a**, Distribution of the IPA scores for the seeds of the up- and down-orientations and respective cluster sizes. **b**, Interface metrics of the DBR3_01 model in complex with ACE2 for the up- and down-orientations were computed using Rosetta's interface analyzer. The following Rosetta metrics are shown: predicted ddG = change in Rosetta energy of separated versus complexed binding partners, interface ΔSASA = solvent accessible surface area buried at the interface, hydrophobic ΔSASA = solvent accessible surface area buried at the interface that is hydrophobic, polar ΔSASA = solvent accessible surface area buried at the interface that is polar, shape complementarity = Lawrence and Coleman shape complementarity of the interface surfaces, # interface residues = number of residues at the interface, # interface Hbonds = number of hydrogen bonds across the interface, ΔunsatHbonds = number of buried, unsatisfied hydrogen bonds at the interface, interface per residue energy = average Rosetta energy of each interface residue, packstat = Rosetta's packing statistic score for the interface ranging from 0 (low packing) to 1 (high packing).

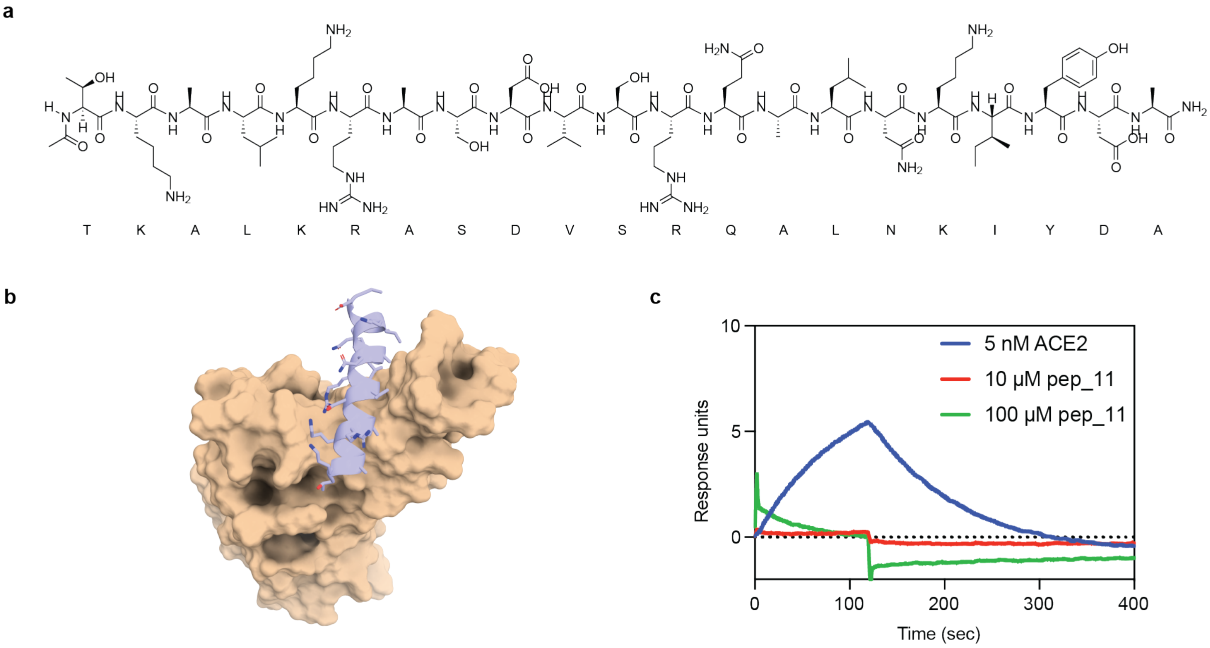

**Supplementary Figure S8: Binding seed identified by MaSIF tested as a synthetic peptide. a**, Structure of the synthesized binding seed. **b**, MaSIF prediction of seed (lavender) binding to RBD (wheat). **c**, SPR data of high concentration of the peptide flowing over RBD. No binding signal is observed for the peptide.

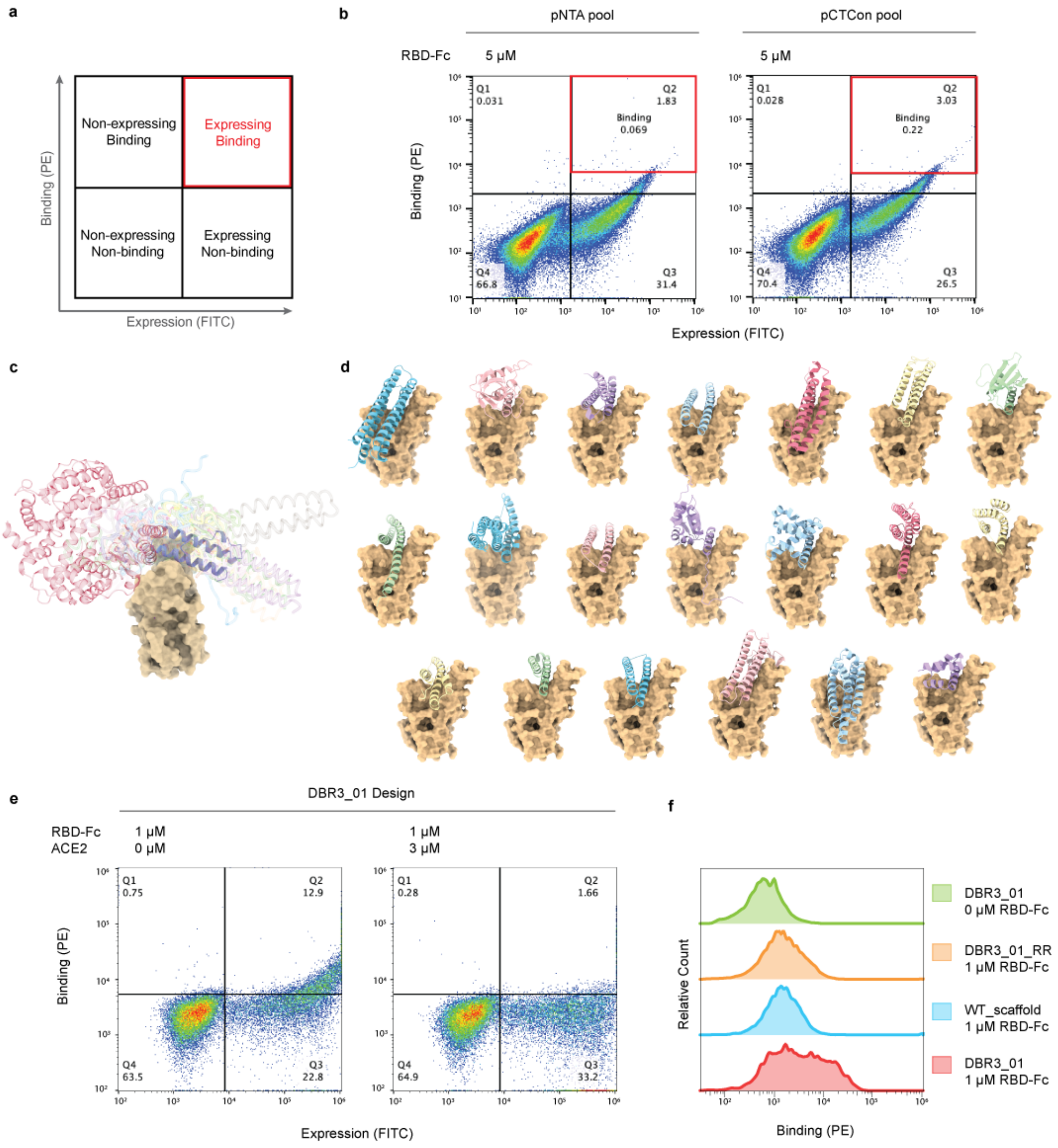

**Supplementary Figure S9: RBD-binder designs displayed on yeast. a,** The yeast display protocol utilizes PE to label binding and FITC to label expression. Yeast appearing in the double positive quadrant are considered potential binders and sorted for enrichment. **b**, Pools of approximately 30 designs were displayed on the surface of yeast and the highest binding populations (red box) sorted for further analysis. **c**, Schematic of RBD (wheat) bound to the various members of the library (transparent silhouettes and purple for DBR3_01) and ACE2 (red) overlapping with the designed binders. **d**, Individual designs DBR1-DBR20. **e**, DBR3_01 design displayed on yeast binds to RBD-Fc (left panel) but the binding is blocked when the RBD-Fc is preincubated with an excess of ACE2, indicating a competitive binding mode. **f**, A point mutant in the binding interface (DBR3_01_RR) and the original scaffold protein (WT_scaffold) show lower binding signal than DBR3_01 with 1 µM RBD-Fc, indicating that the design is engaging the RBD with the predicted interface.

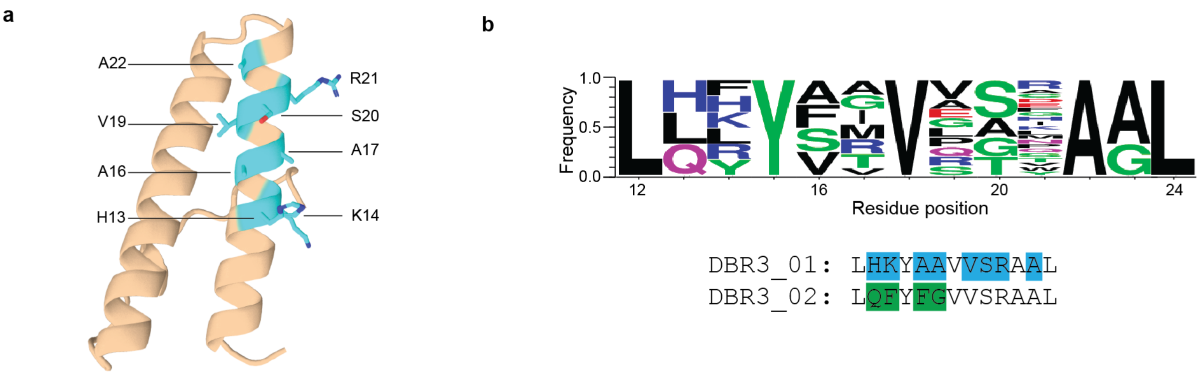

**Supplementary Figure S10: Directed Library for DBR3_01.** **a**, Position of residues included in a combinatorial library to improve binding affinity. **b**, Sequence logo plot of specific mutations allowed within the library. The sequences list the residues mutated in DBR3_01 (highlighted in blue) and the mutations gained through the library in DBR3_02 (green).

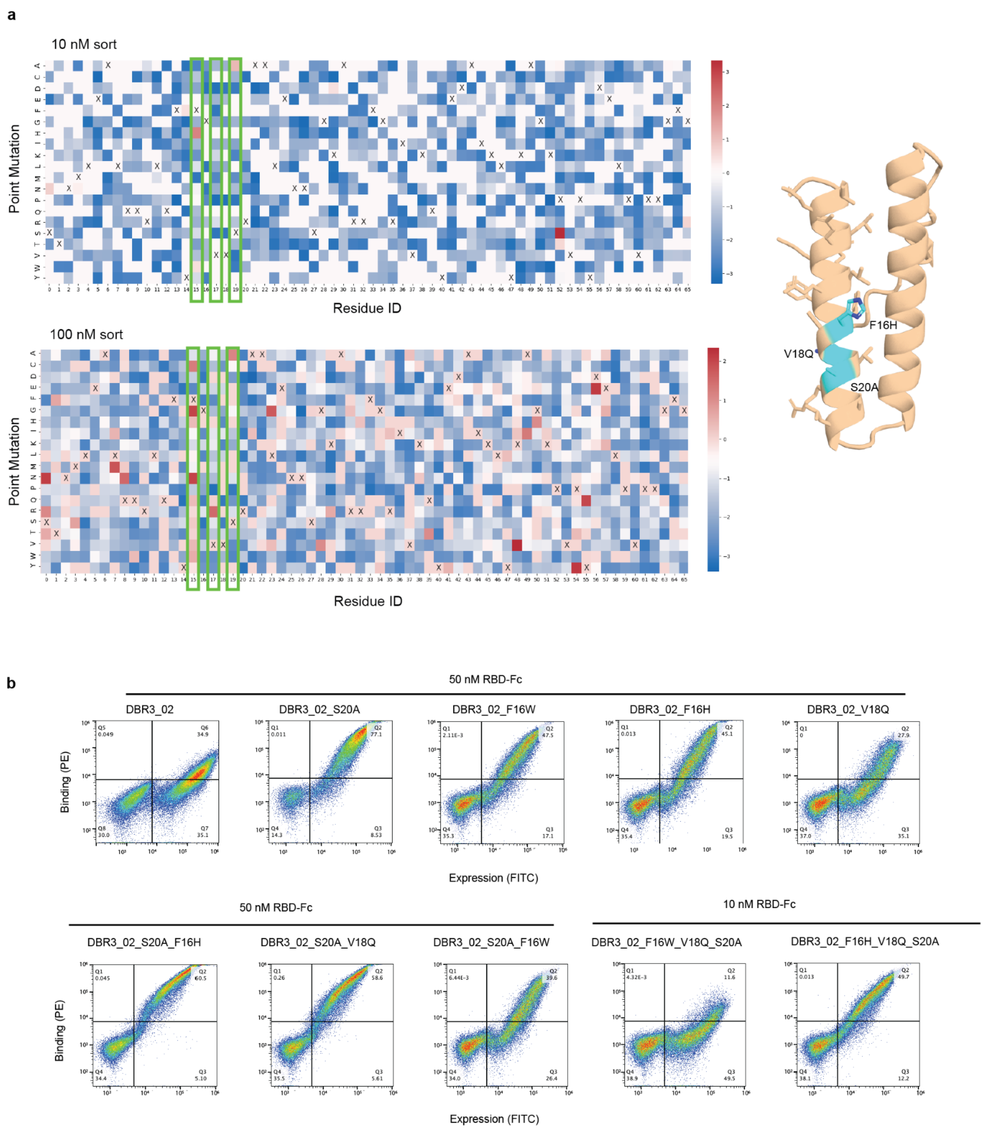

**Supplementary Figure S11: SSM of DBR3_02. a**, Heat maps of DBR3_02 SSM at two concentrations of RBD-Fc. X indicates the original amino acid of DBR3_02. Red indicates an enrichment of the mutation in the binding population, blue indicates an enrichment in the non-binding population. Three positions, green box, were enriched in both concentrations. The positions of these mutations are highlighted on the DBR3_03 structure. **b**, Yeast display of DBR3_02 with mutations from the SSM introduced shows increase in affinity to RBD.

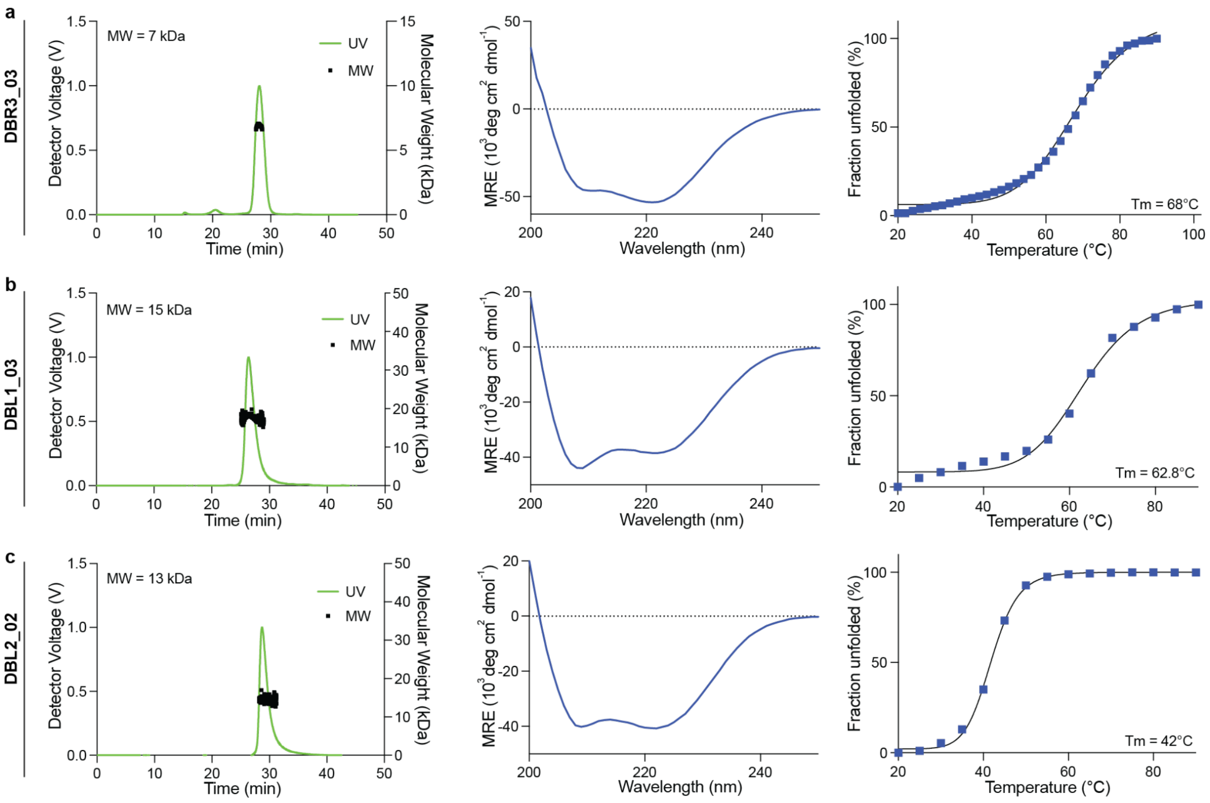

**Supplementary Figure S12: Biophysical characterization of the designed binders.** From left to right: The oligomeric status was determined via multi-angled light scattering (MALS). Folding was measured using circular dichroism. Thermal stability was determined by plotting the ellipticity at 218 nm at increasing temperatures. **a**, DBR3_03, **b**, DBL1_03, **c**, DBL2_02.

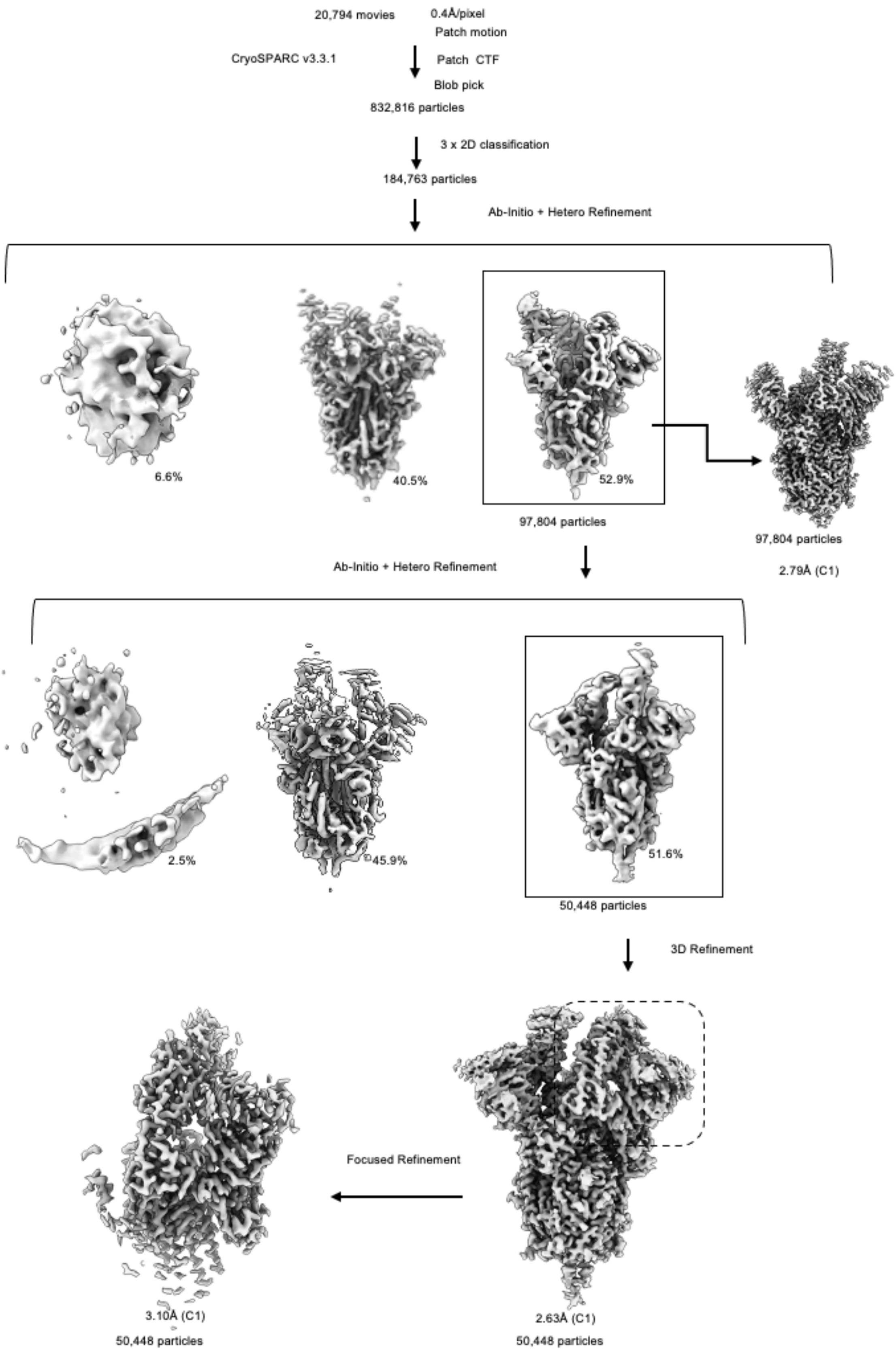

**Supplementary Figure S13: Cryo-EM data processing of the D614G Spike-DBR3_03 complex.** Image processing workflows performed in CryoSPARC v.3.3.1.

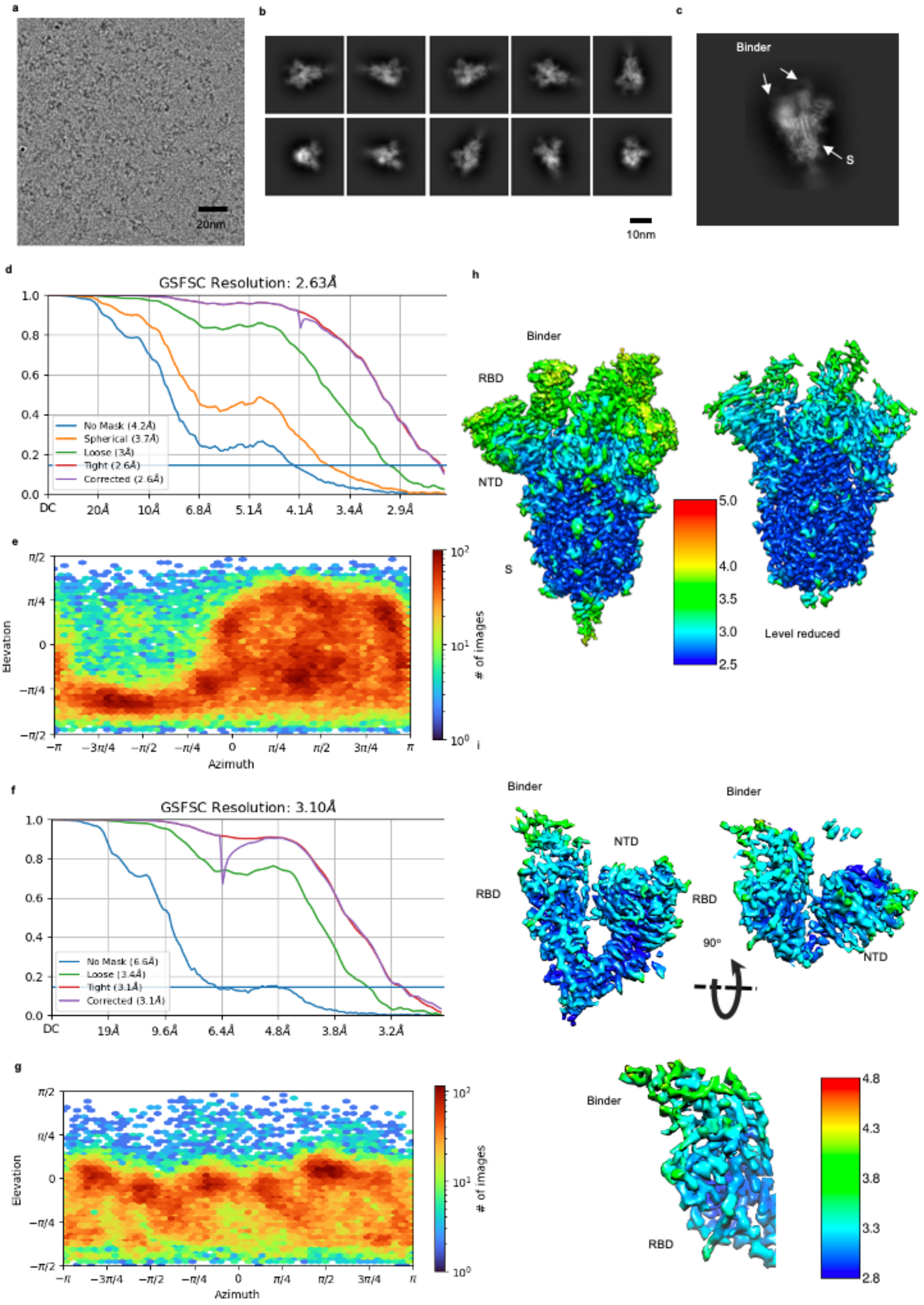

**Supplementary Figure S14: Details of Cryo-EM data processing for D614G Spike-DBR3_03 complex.** **a,** A representative raw micrograph of the Cryo-EM sample for D614G Spike-binder complex. **b,** The 2D classes of the D614G Spike-binder complex. **c,** A representative 2D class. **d,** Direction distribution of the particle alignment and **e,** FSC curves of the final overall map. **f,** Direction distribution and **g,** FSC curves of the locally refined map. **h,i,** Local resolution distribution of the overall and focused refined maps.

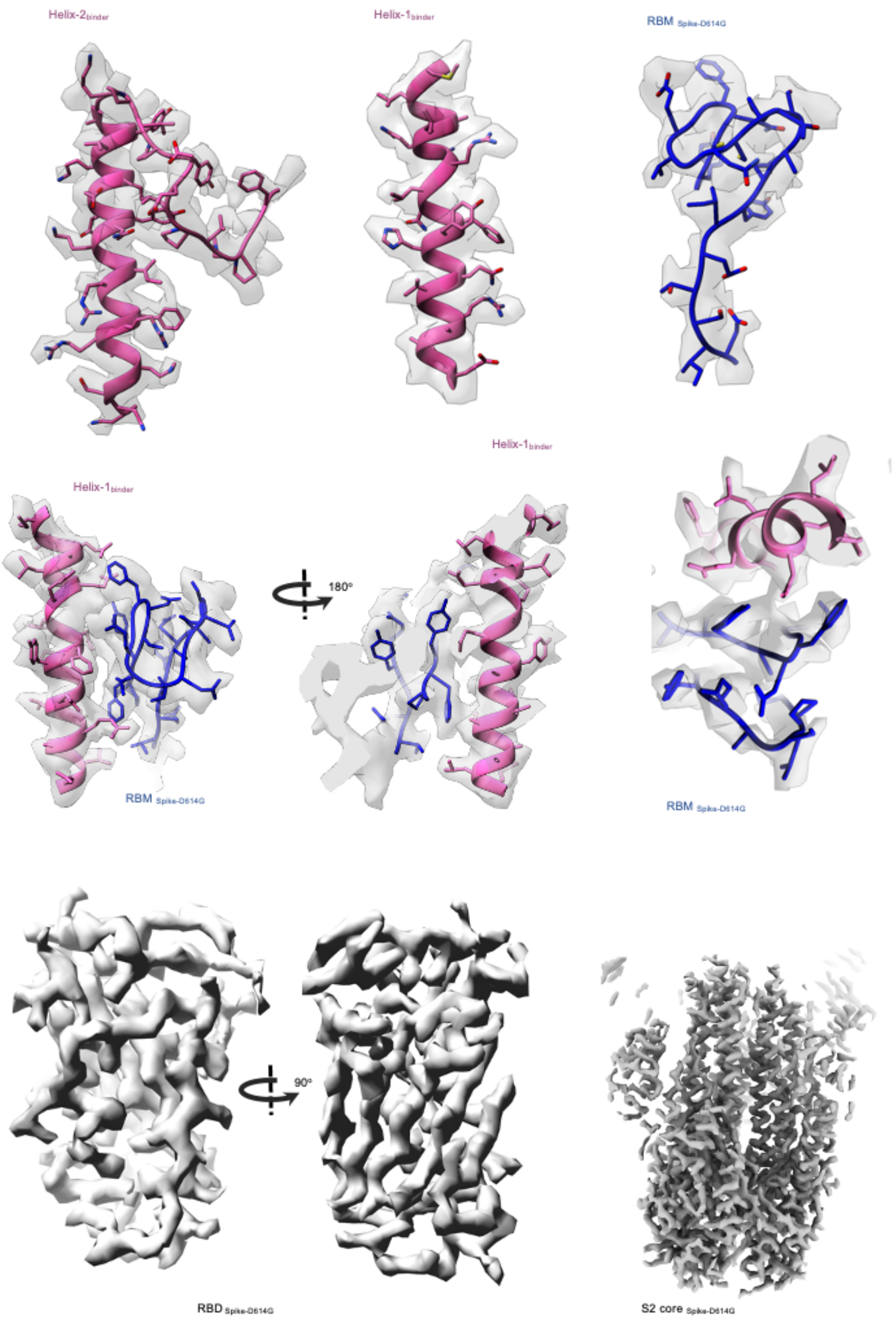

**Supplementary Figure S15: Highlights of the Cryo-EM densities of DBR3_03 with D614G spike.** Cryo-EM densities are shown as surfaces. RBM (receptor binding motif) in blue with DBR3_03 in pink. The atomic model is shown as stick or ribbon representation.

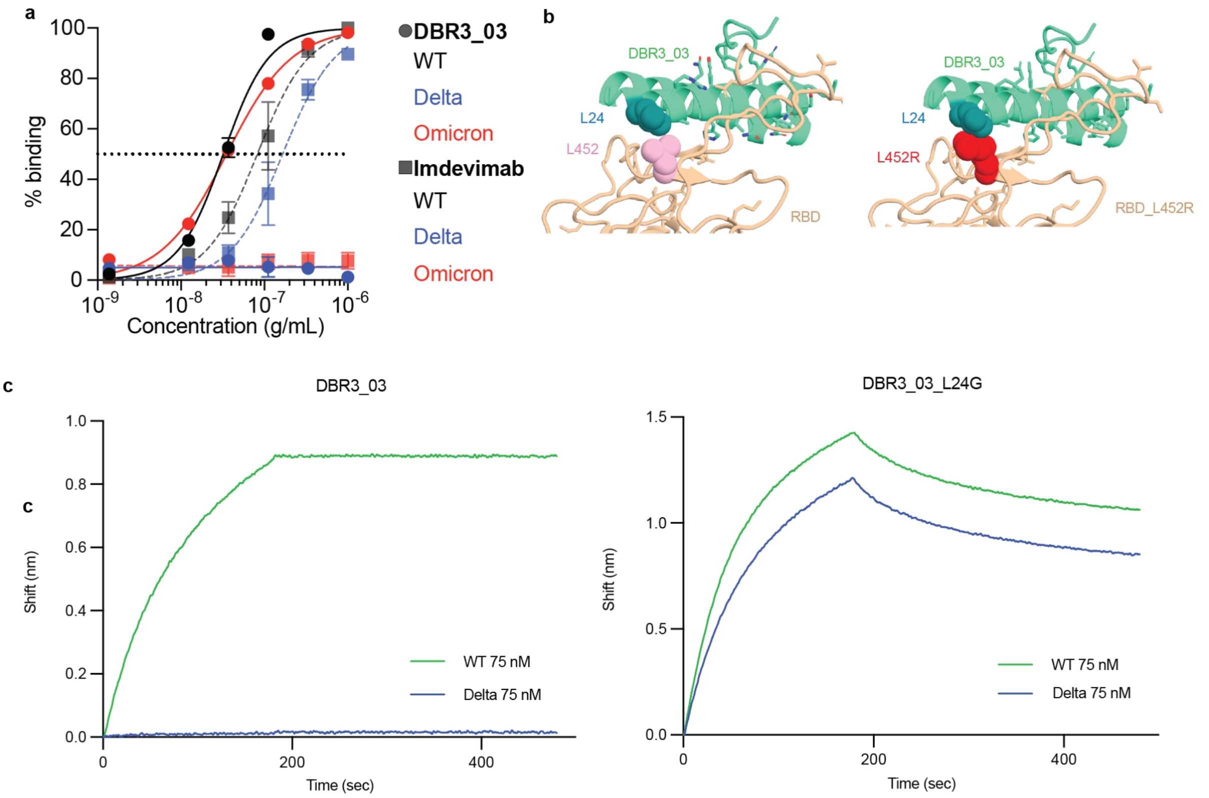

**Supplementary Figure S16: DBR3_03 binding is sensitive to the L452R mutation in the spike protein.** **a**, Luminex binding assay of DBR3_03 or Imdevimab (REGN10987) with beads functionalized with SARS-CoV-2 spike protein of indicated variants. DBR3_03 has an $EC_{50}$ of $3.2e^{-8}$ g/mL with WT and $3.5e^{-8}$ g/mL with omicron. Imdevimab has an $EC_{50}$ of $8.2e^{-8}$ g/mL with WT and $1.7e^{-7}$ g/mL with delta. **b**, The L452R mutation on the spike protein leads to a clash with the DBR3_03 binding. A L24G mutation is proposed to avoid the clash. **c**, BLI data with DBR3_03 (WT $K_D$<0.1 nM, delta $K_D$ not detected) or DBR3_03_L24G (delta $K_D$=6 nM, WT $K_D$=6 nM) immobilized on the tips, dipped into spike protein of different variants.

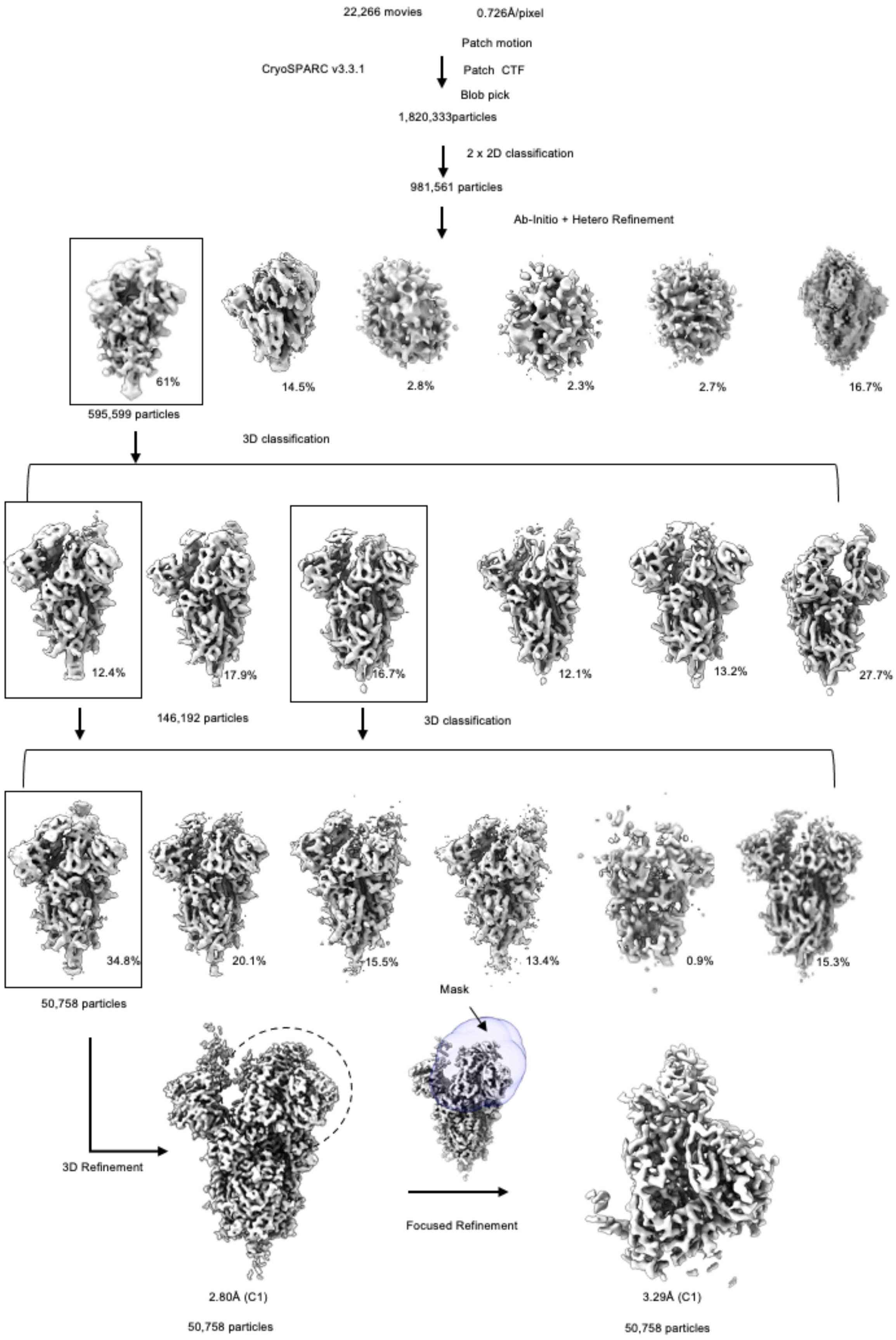

**Supplementary Figure 17: Cryo-EM data processing of the Omicron Spike-DBR3_03 complex.** Image processing workflows performed in CryoSPARC.

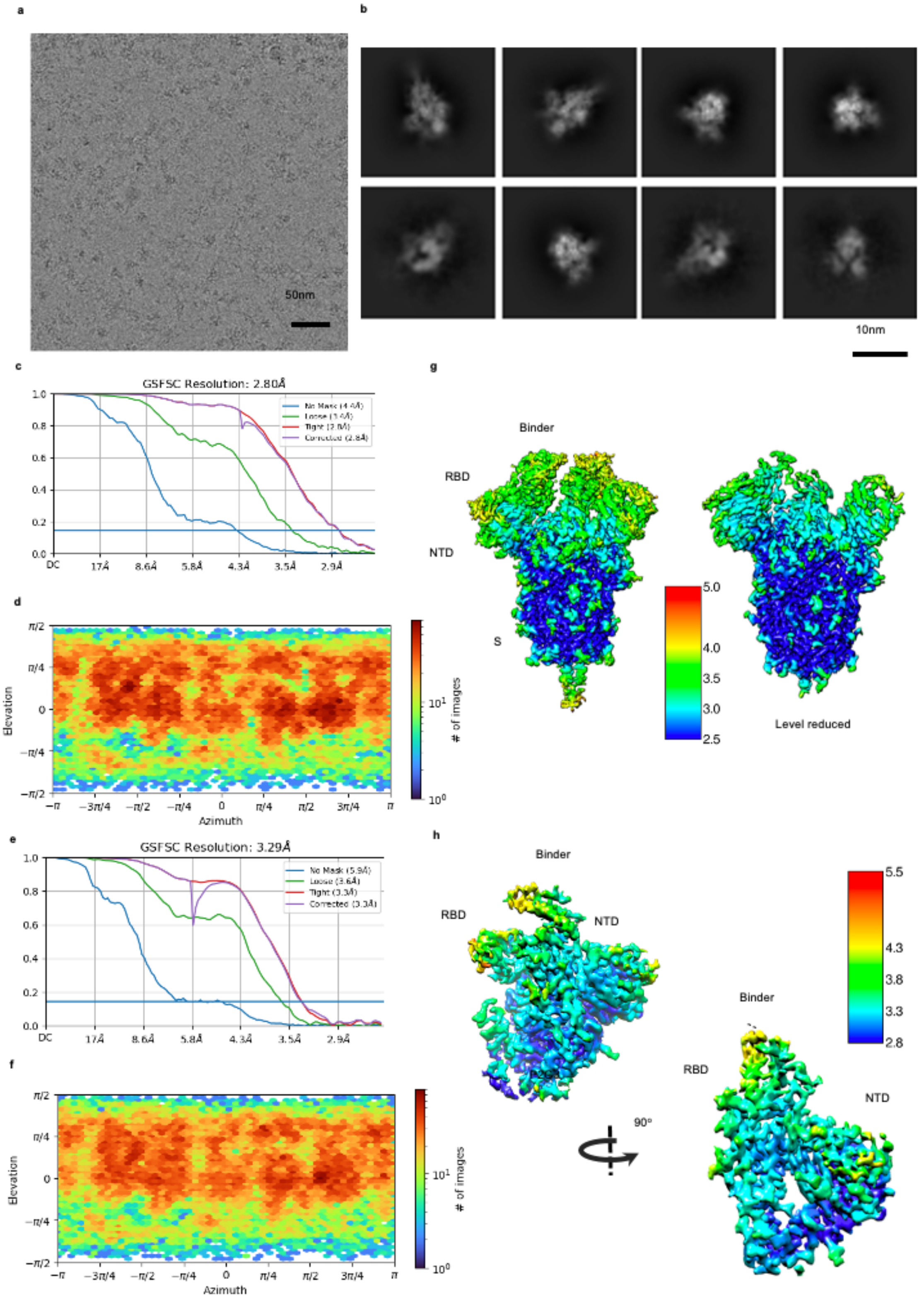

**Supplementary Figure 18: Details of Cryo-EM data processing for Omicron Spike-DBR3_03 complex.** **a,** A representative Cryo-EM micrograph for the D614G Spike-binder complex. **b,** The representative 2D classes of the omicron Spike-binder complex. **c,** Direction distribution of the particle alignment and **d,** FSC curves of the final overall map. **e,** Direction distribution and **f,** FSC curves of the locally refined map. **g,h,** Local resolution distribution of the overall and focused refined maps.

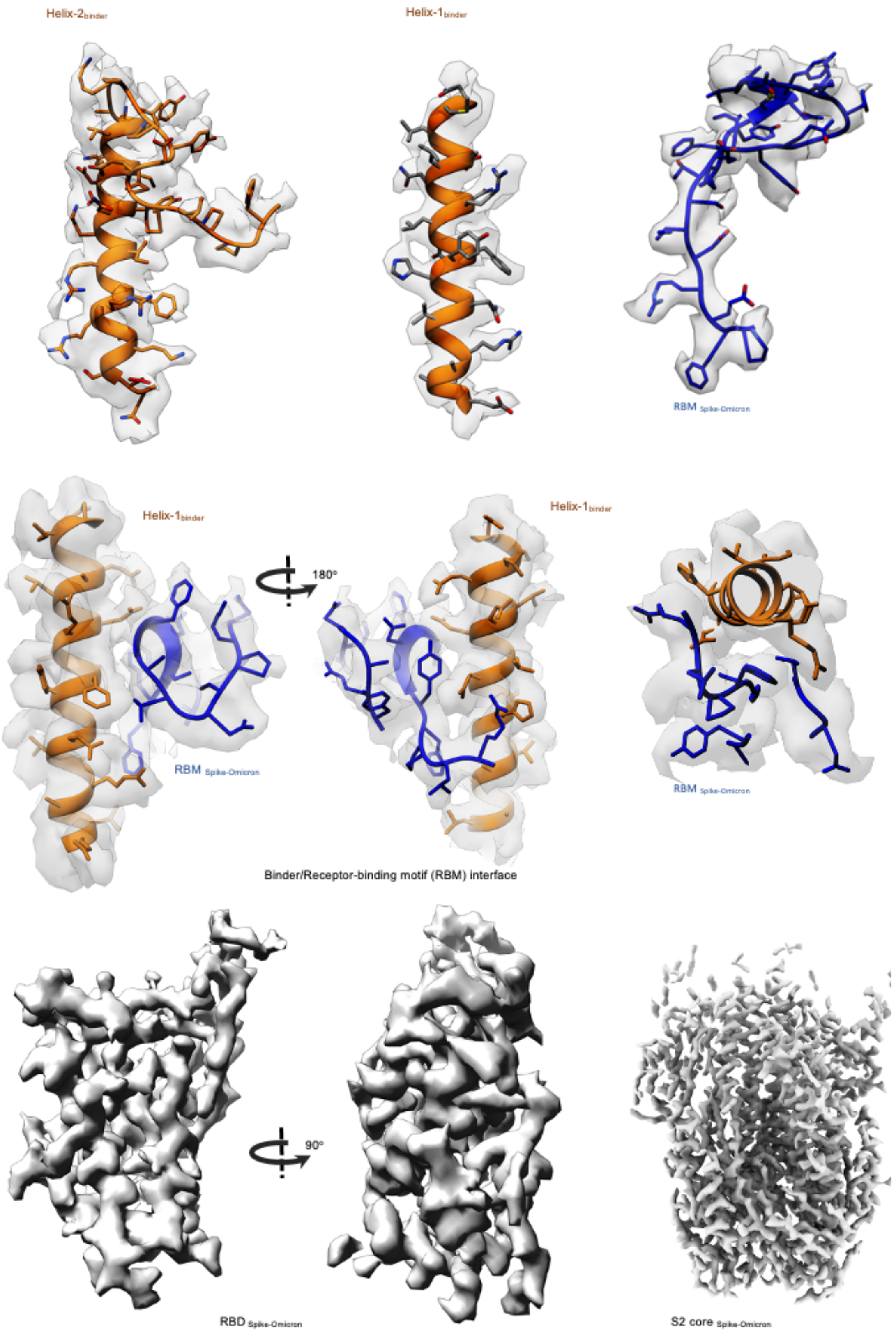

**Supplementary Figure 19: Highlights of the cryo-EM densities of DBR3_03 with Omicron spike.** Cryo-EM densities are shown as surfaces. RBM (receptor binding motif) in blue with DBR3_03 in orange. The atomic model is rendered as stick or ribbon representation.

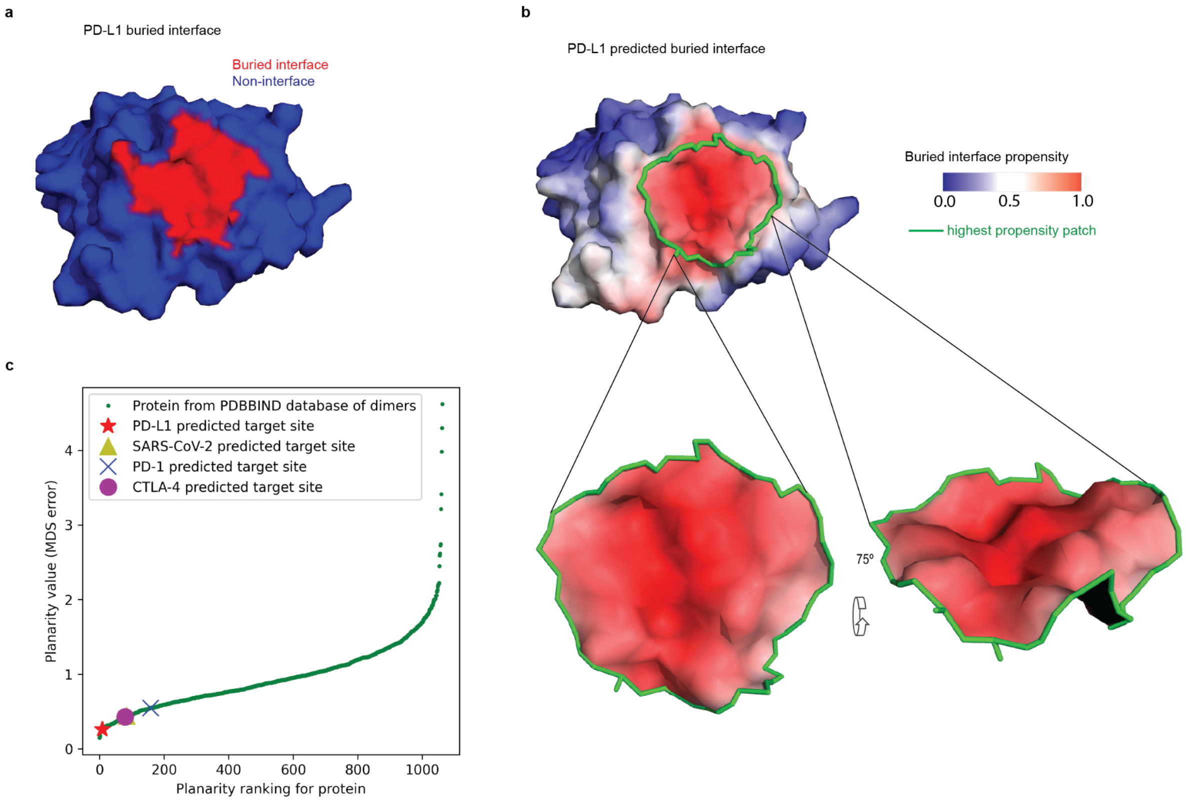

**Supplementary Figure S20: Planarity of the targeted interface sites. a**, Buried interface on PD-L1 upon complex formation with PD-1. **b**, (Top) PD-L1 predicted buried interface, with selected target patch marked with a green contour. (bottom) View of the selected target patch to show its planarity. **c**, Plotting of the planarity of each of 1068 dimeric protein interfaces. Y-axis: error in multidimensional scaling when flattening the patch from 3D to 2D. X-axis: ranking of each protein according to the planarity value with respect to the dataset of 1068 dimeric protein interfaces. The PD-L1 interface targeted in this work is marked with a red star, SARS-CoV-2 with a gold triangle, PD-1 with a blue X, and CTLA-4 with a magenta circle.

a

b

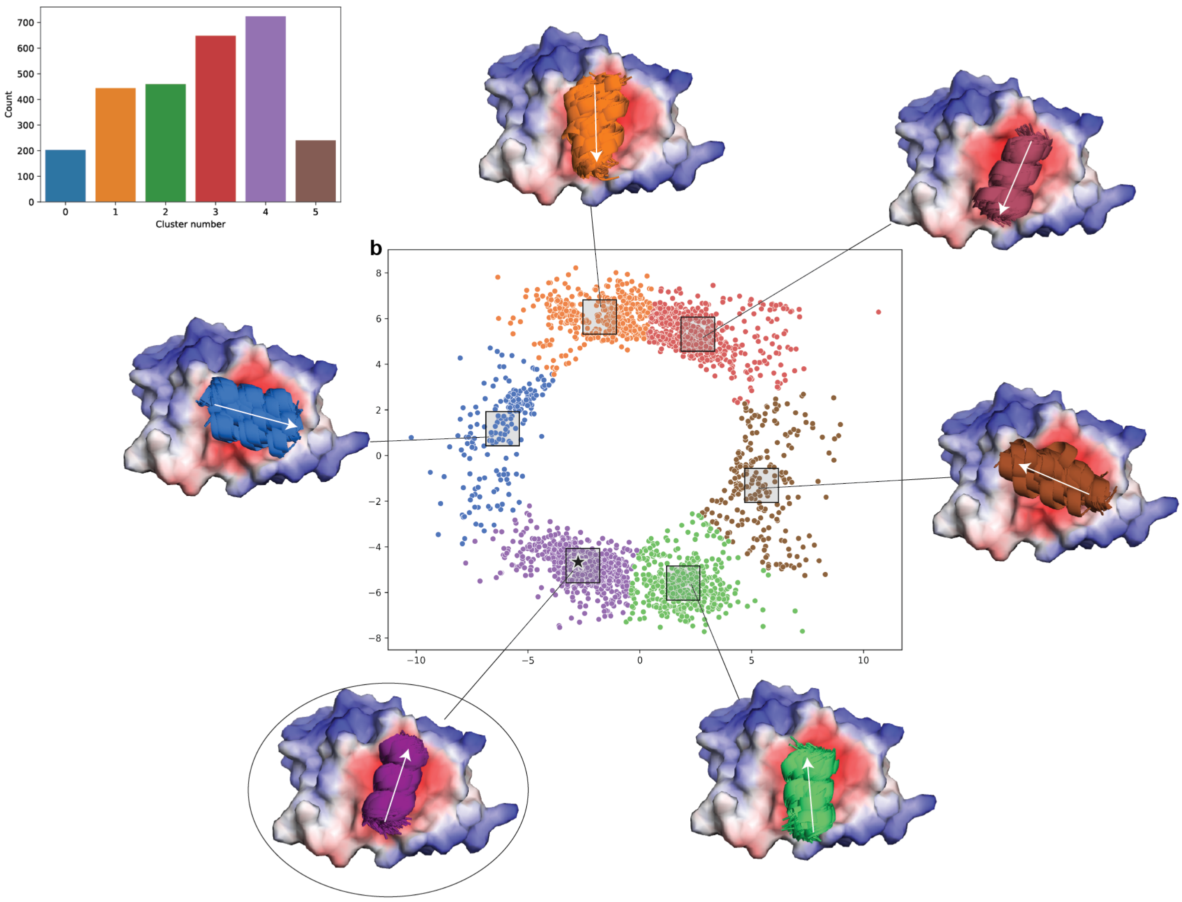

**Supplementary Figure S21: Clusters of putative binding seeds identified by MaSIF-seed docked on the PD-L1 surface (PDB ID: 5JDS).** 140 million patches from ~250,000 helices extracted from the PDB were compared and docked to the predicted interface in PD-L1 using MaSIF-seed. The top scoring seeds were selected for further processing. Twelve-amino acid fragments of these seeds that occupied the largest buried surface were then clustered using metric multidimensional scaling of all pairwise RMSDs between all seeds. **a,** Histogram of clusters, showing the prevalence of each orientation. **b,** Binding seed clusters in the multidimensional scaling plot. A box is drawn around the center of each cluster and the picture shows the selected helix orientation for all points inside the box. The circled binding seed cluster shows the helix orientation of the seed used for the PD-L1 designs. A star symbol shows the PD-L1 seed used for the designs.

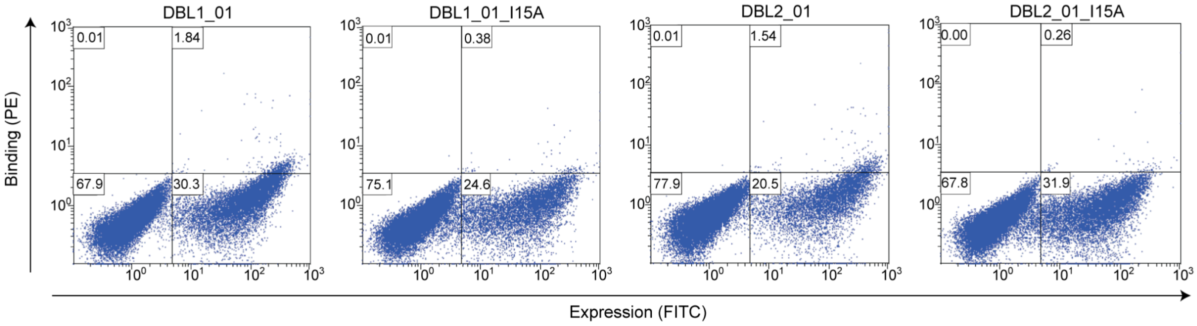

**Supplementary Figure S22: Binding signals of initial PD-L1 binder designs**. Binding measured on the surface of yeast with 15 µM PD-L1-Fc. Comparison of DBL1_01 and DBL2_01 with corresponding interface mutants.

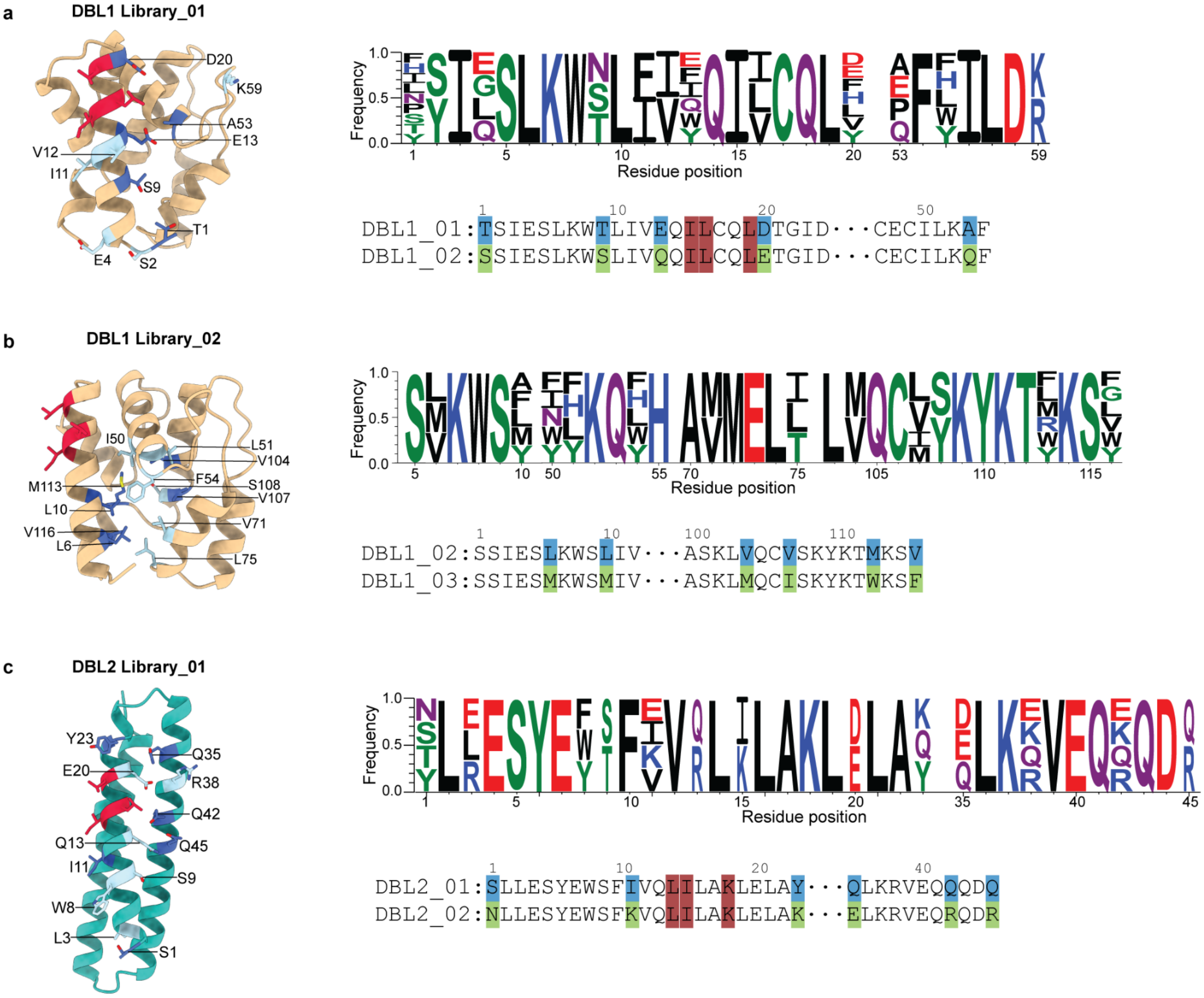

**Supplementary Figure S23: Composition and outcome of yeast display libraries.** **a**, Position of targeted residues in the structure of DBL1_01 to improve binding affinity. Logo plot of the allowed mutations in the library and alignment of initial design with library enriched design. **b**, Position of targeted residues in the structure of DBL1_02 to improve core packing. Logo plot of the allowed mutations in the library and alignment of DBL1_02 with library enriched design. **c**, Position of targeted residues in the structure of DBL2_01 to improve binding affinity and solubility. Logo plot of the allowed mutations in the library and alignment of initial design with library enriched design. Hotspot residues red, targeted residues light blue, mutated residues dark blue.

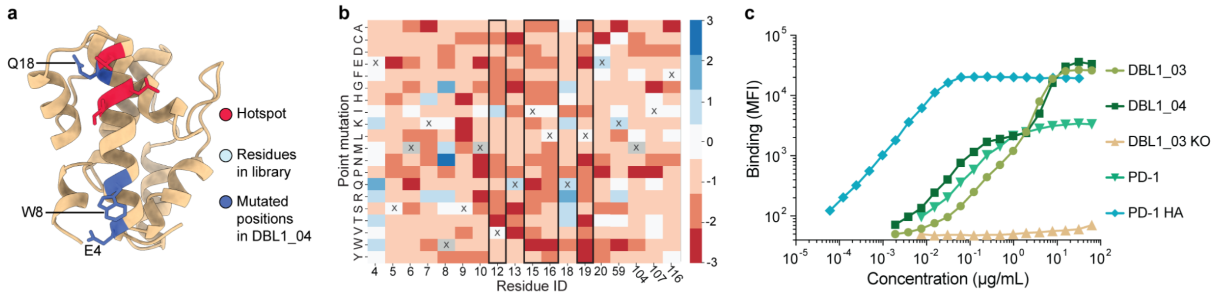

**Supplementary Figure S24: Complete SSM library of DBL1_03 and cell binding data. a,** Structural representation of all positions sampled in the SSM library (light blue). The four hotspot residues (red) were also sampled. Three positions were mutated in DBL1_04 (dark blue). **b,** Outcome of the entire SSM library. Blue indicates enrichment in the binding population, while red shows enrichment in the non-binding population. **c,** Binding of DBL1_03 and DBL1_04 to KARPAS299 cells expressing PD-L1 compared to binding of WT PD-1, a high affinity version of PD-1 (PD-1_HA)[34] and a V12R mutation of DBL1_03 (KO). All proteins contained a Fc domain.

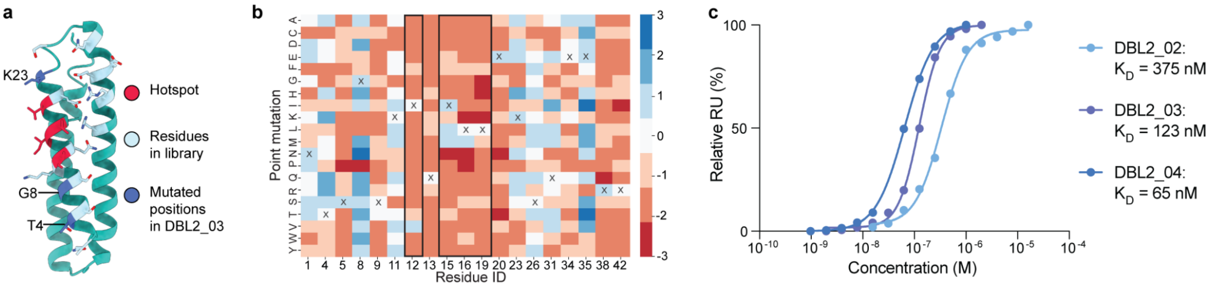

**Supplementary Figure S25: Overview of DBL2_03 SSM library. a,** Structural representation of all positions sampled in the SSM library (light blue). The four hotspot residues (red) were also sampled. Three positions were mutated in DBL2_04 (dark blue). Position 35 was not mutated in DBL_04, because all mutations in this position led to the inability of the soluble expression of the protein. **b,** Outcome of the entire SSM library. Blue indicates enrichment in the binding population, while red shows enrichment in the non-binding population. **c,** Binding affinities measured by SPR for the different versions of DBL2.

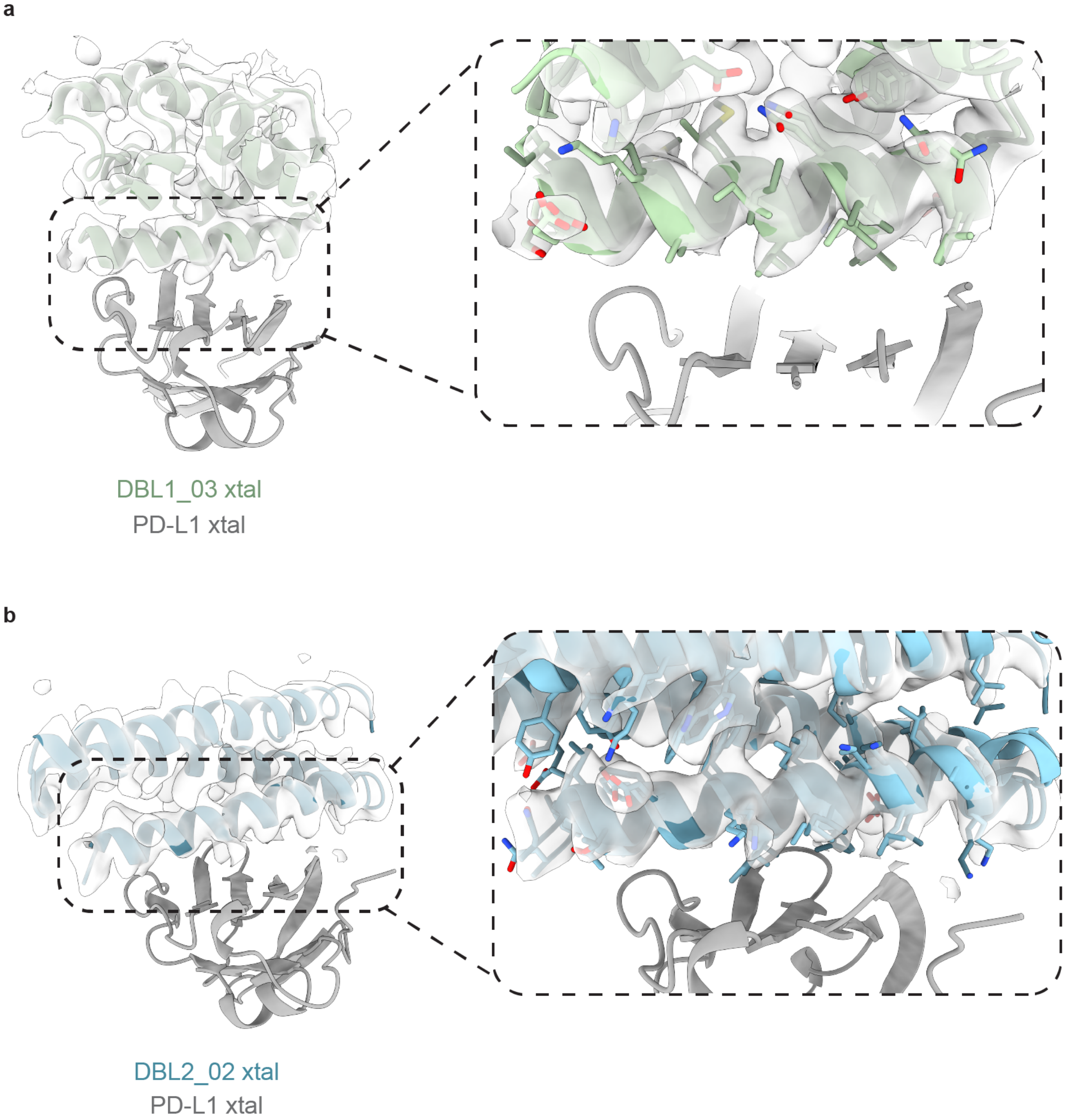

**Supplementary Figure S26: Electron density map of the crystalized DBL1_03 and DBL2_02**. **a**, Crystal structure of DBL1_03 (green) in complex with PD-L1 (gray). Refined 2mFo-mFc electron density map of the binder, contoured at 1.0$\sigma$, is rendered as a white surface **b**, Crystal structure of DBL2_02 (blue) in complex with PD-L1 (gray). Refined 2mFo-mFc electron density map of the binder, contoured at 1.0$\sigma$, is rendered as a white surface

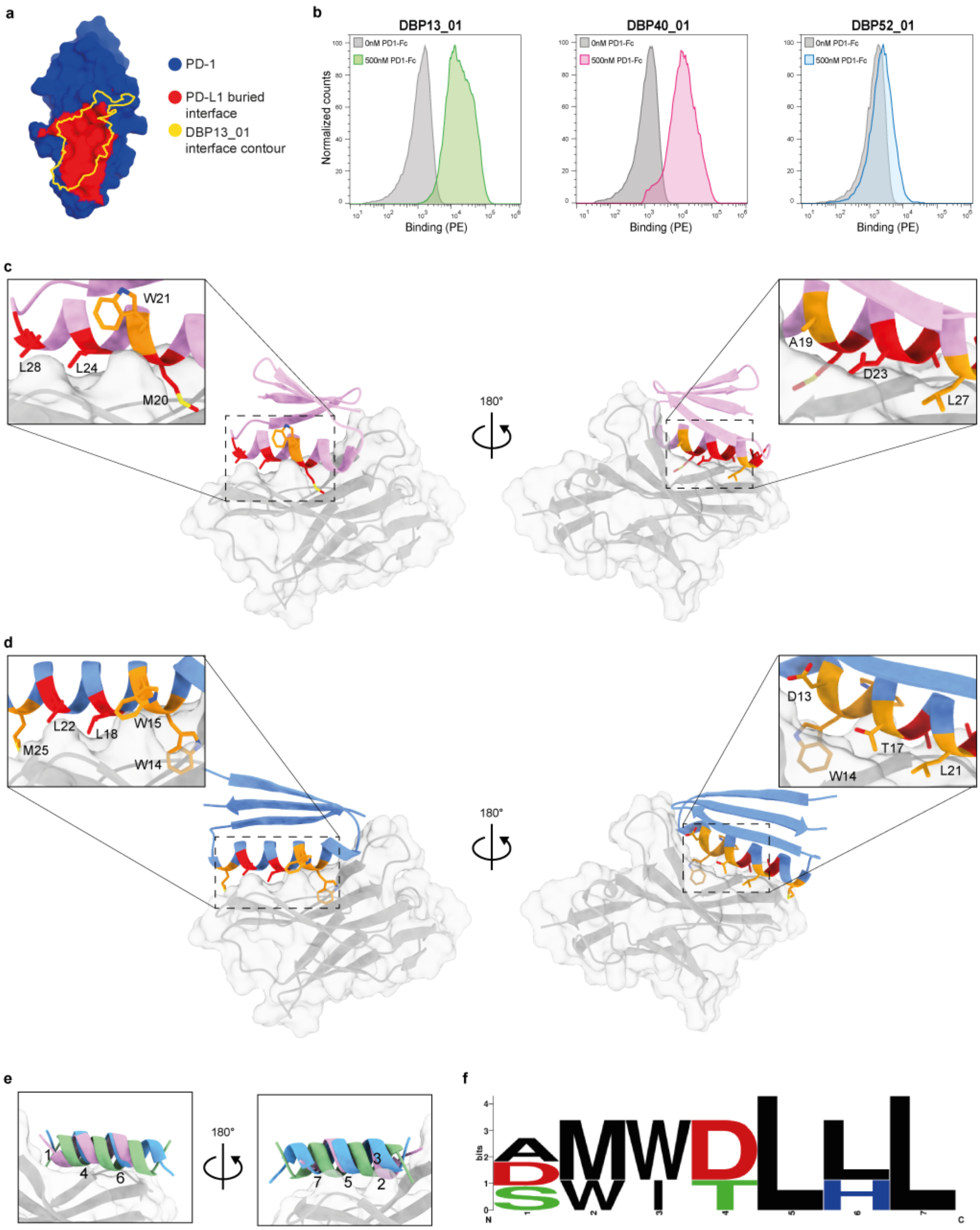

**Supplementary Figure S27: Overview and comparison between PD-1 binders. a,** PD-1 surface (blue) with the region targeted by PD-L1 (red) and the overlapping region targeted by DBP13_01 (yellow contour). **b**, Histograms of the binding signal (PE) measured on 3 yeast clones displaying designed binders against PD-1. Yeast cells were labeled with 500 nM PD-1-Fc (coloured) or secondary antibodies only (gray, negative control). **c-d**, Overview and close-up of DBP40_01 (a, pink) and DBP52_01 (b, blue) models in complex with PD-1 (gray). Interface seed residues similar to DBP13_01 are highlighted in red, while residues that are different are highlighted in orange. **e,** Seeds used to design DBP13_01 (green), DBP40_01 (pink) and DBP52_01 (blue) aligned with interface residues numbered. **f,** Sequence logo of the seed interface residues for the three PD-1 binders as numbered in e.

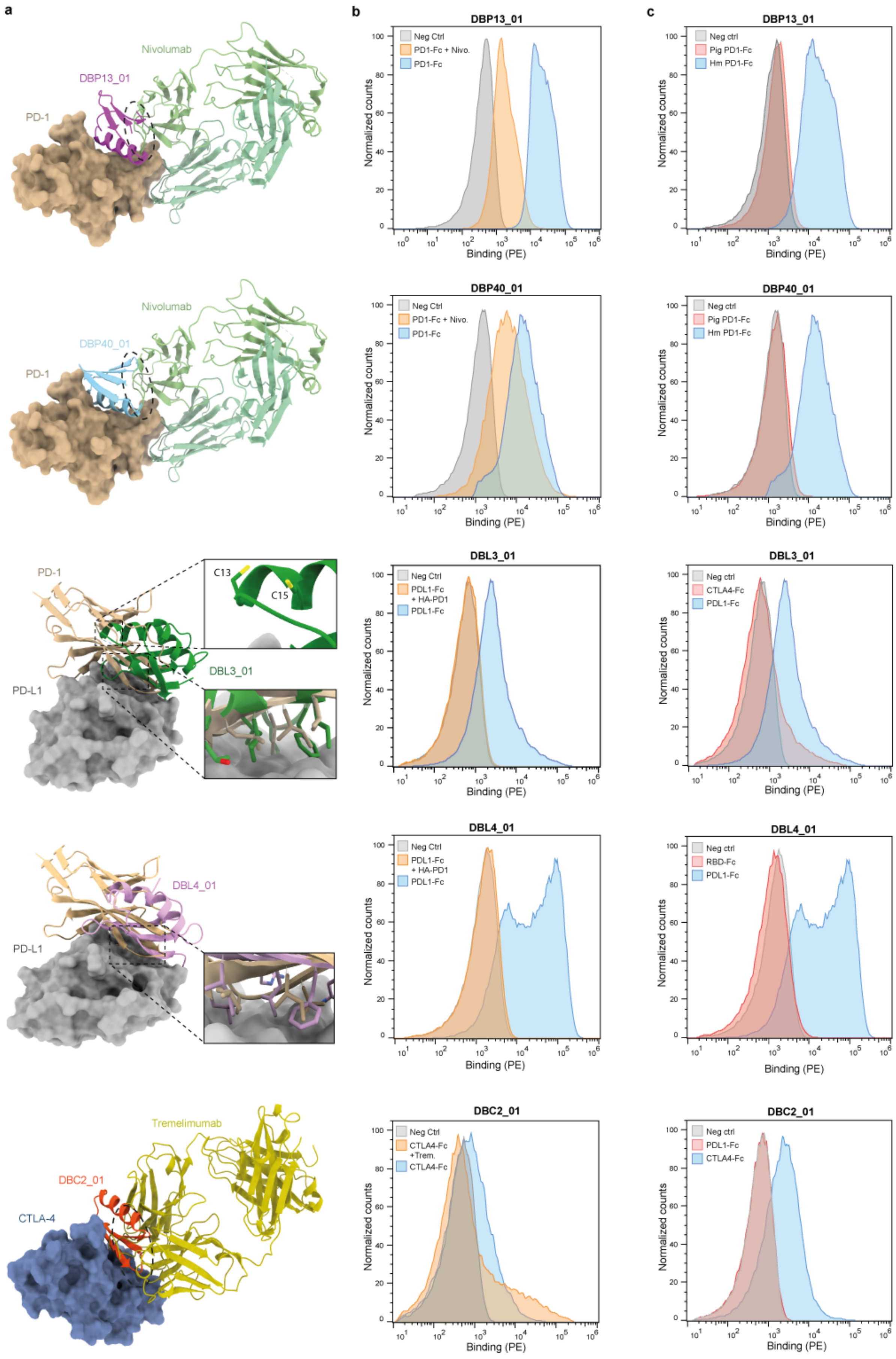

a
Nivolumab
DBP13_01
PD-1
Nivolumab
DBP40_01
PD-1
PD-1
C13
C15
DBL3_01
PD-L1
DBL4_01
PD-L1
Tremelimumab
DBC2_01
CTLA-4
b
DBP13_01
Neg Ctrl
PD1-Fc + Nivo.
PD1-Fc
DBP40_01
Neg Ctrl
PD1-Fc + Nivo.
PD1-Fc
DBL3_01
Neg Ctrl
PDL1-Fc + HA-PD1
PDL1-Fc
DBL4_01
Neg Ctrl
PDL1-Fc + HA-PD1
PDL1-Fc
DBC2_01
Neg Ctrl
CTLA4-Fc +Trem.
CTLA4-Fc
c
DBP13_01
Neg ctrl
Pig PD1-Fc
Hm PD1-Fc
DBP40_01
Neg ctrl
Pig PD1-Fc
Hm PD1-Fc
DBL3_01
Neg ctrl
CTLA4-Fc
PDL1-Fc
DBL4_01
Neg ctrl
RBD-Fc
PDL1-Fc
DBC2_01
Neg ctrl
PDL1-Fc
CTLA4-Fc
Normalized counts
Binding (PE)

**Supplementary Figure S28: Competition and specificity binding assay of the different optimized binders on the surface of yeast. a,** Competition between designed binders and a known protein binder (native binder or  monoclonal Fab) in complex with the target structure. **b**, Flow cytometry histograms showing fluorescence signals on the surface of yeast displaying the different binders. Yeast were labeled with 500 nM or their respective ligand (blue), 500 nM of blocked ligand pre-incubated with 10-fold molar excess of Fab or high-affinity PD-1 (HA-PD-1) (orange) or labeled with secondary antibodies only (gray, Neg Ctrl). **c,** Flow cytometry histograms showing fluorescence signal on the surface of yeast displaying the different binders and labeled with 500 nM of unrelated protein ligand (red) or labeled with secondary antibodies only (gray, Neg Ctrl).

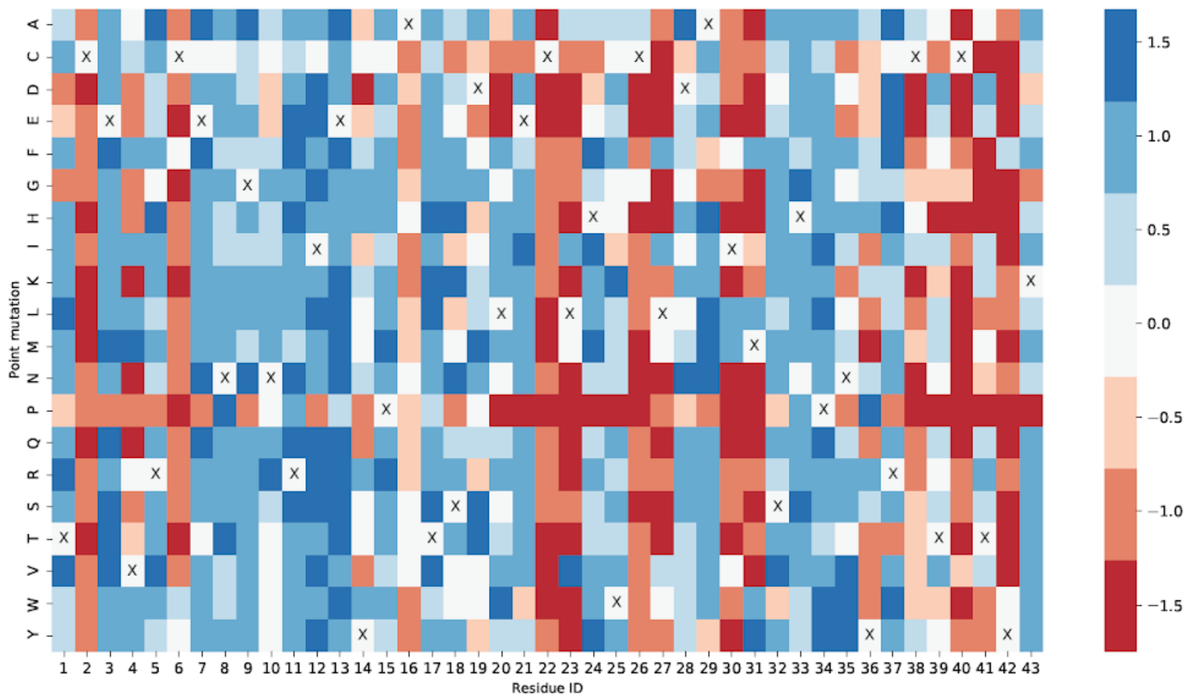

**Supplementary Figure S29: SSM of DBP13_01.** Heatmap covering all positions of DBP13_01. Yeast displaying point mutants were analyzed by flow cytometry and subsequently binding and non- binding populations were sorted. For each mutation the log-ratio between the enrichment in binding versus non-binding populations was computed. Mutations in red highlight a deleterious effect on binding, while mutations in blue indicate an enrichment on the binding population.

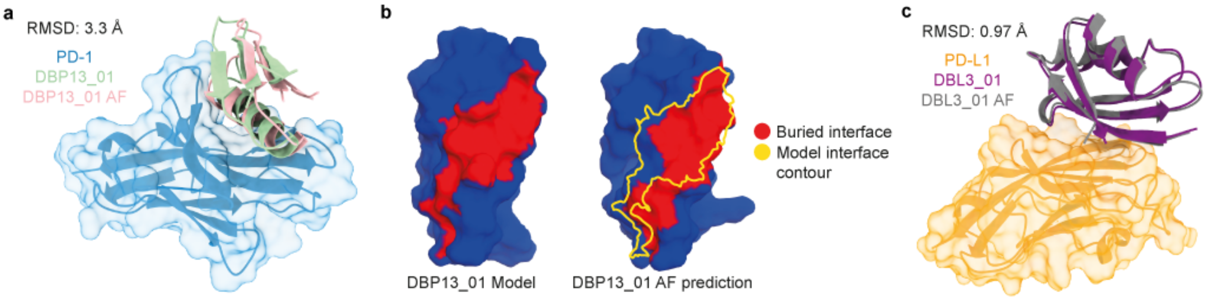

**Supplementary Figure S30: AF structure prediction of DBP13_01 in complex with PD-1.** **a**, Comparison of the DBP13_01 computational model (green) and the AlphaFold multimer (AF) prediction (red) on the surface of PD-1 (blue). **b**, Buried interfaces in both DBP13_01 model (left) and AF prediction (right) are shown in red with an overlap yellow, a yellow contour of the footprint of the original model is shown for ease of comparison. **c**, Comparison of the DBL3_01 computational model (purple) and the AlphaFold Multimer (AF) prediction (gray) on the surface of PD-L1 (orange).

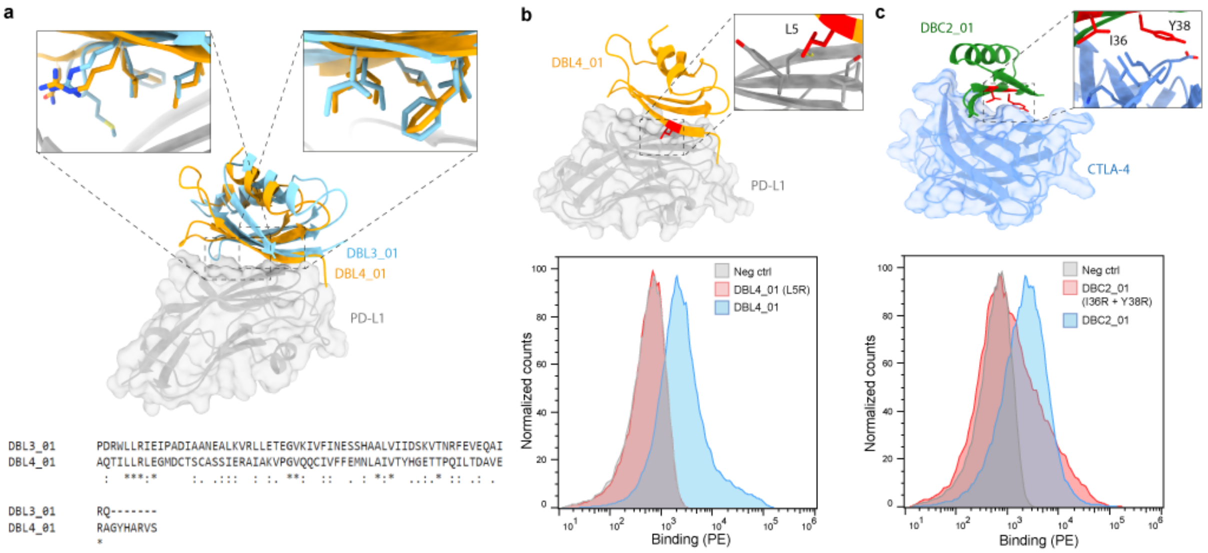

**Supplementary Figure S31: DBL3_01 and DBL4_01 comparison and DBL4_01 and DBC2_01 knock-out mutants. a**, superposition between DBL3_01 (cyan) and DBL4_01 (orange) in complex with PD-L1 (gray). Multiple sequence alignment of the two designs is shown at the bottom. **b**, DBL4_01 (orange) in complex with PD-L1 (gray) with knock-out mutant highlighted in red. Flow cytometry histograms showing fluorescence signals on the surface of yeast displaying DBL4_01 or the knock-out mutant, compared to unlabeled yeast (Neg Ctrl). **c**, DBC2_01 (green) in complex with CTLA-4 (blue) with two knock-out mutants highlighted in red. Flow cytometry histograms showing fluorescence signals on the surface of yeast displaying DBC2_01 or the knock-out mutants, compared to unlabeled yeast (Neg Ctrl).

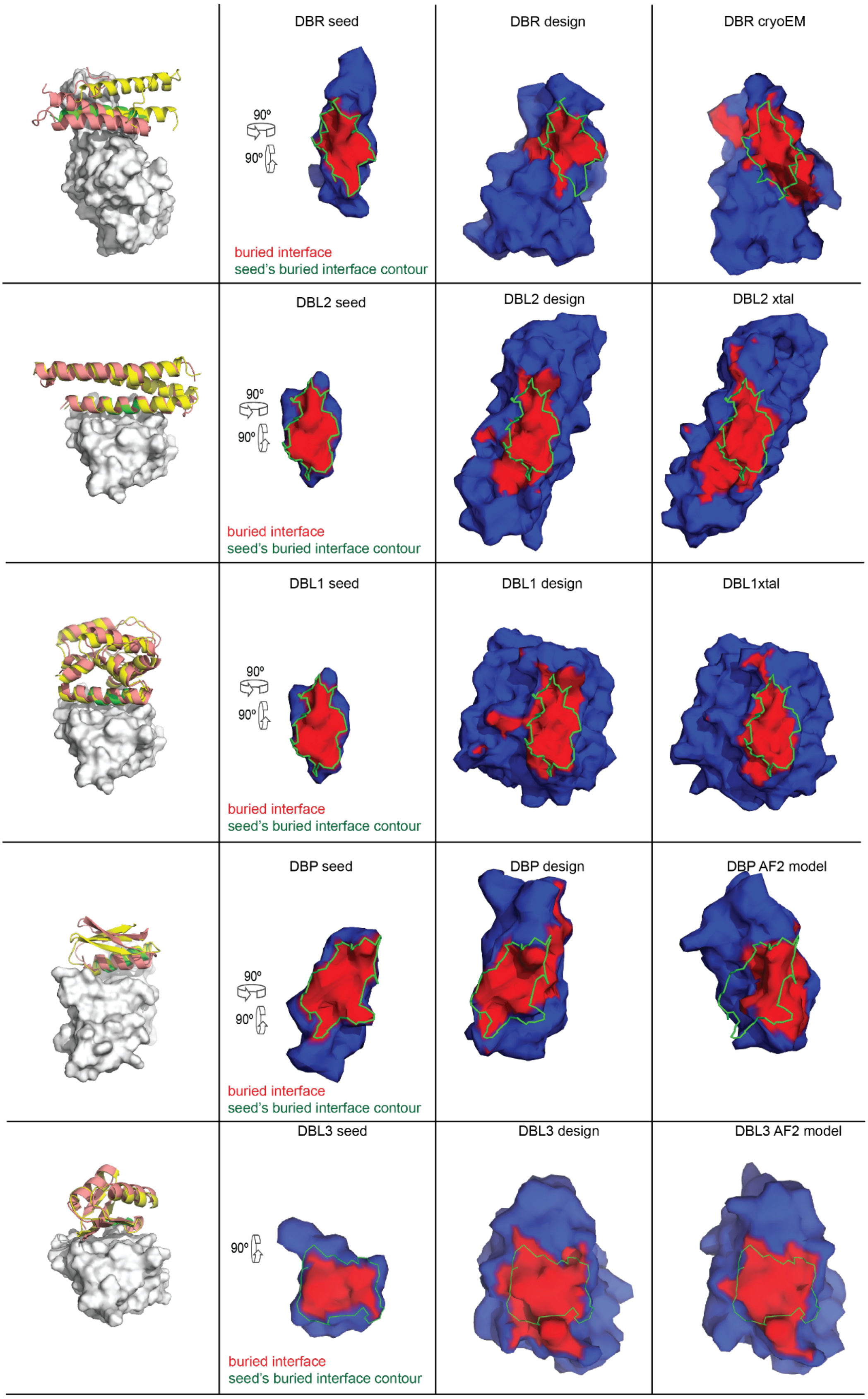

**Supplementary Figure S32: Surface comparison between seeds, designs and final/predicted structures**. Buried interfaces of models/structures when in complex with their target are colored in red, while non-buried regions colored in blue. The contour of the buried interface of the initial binding seed is drawn in green and is shown for the initial seed, for the designs and for the final/predicted structures.

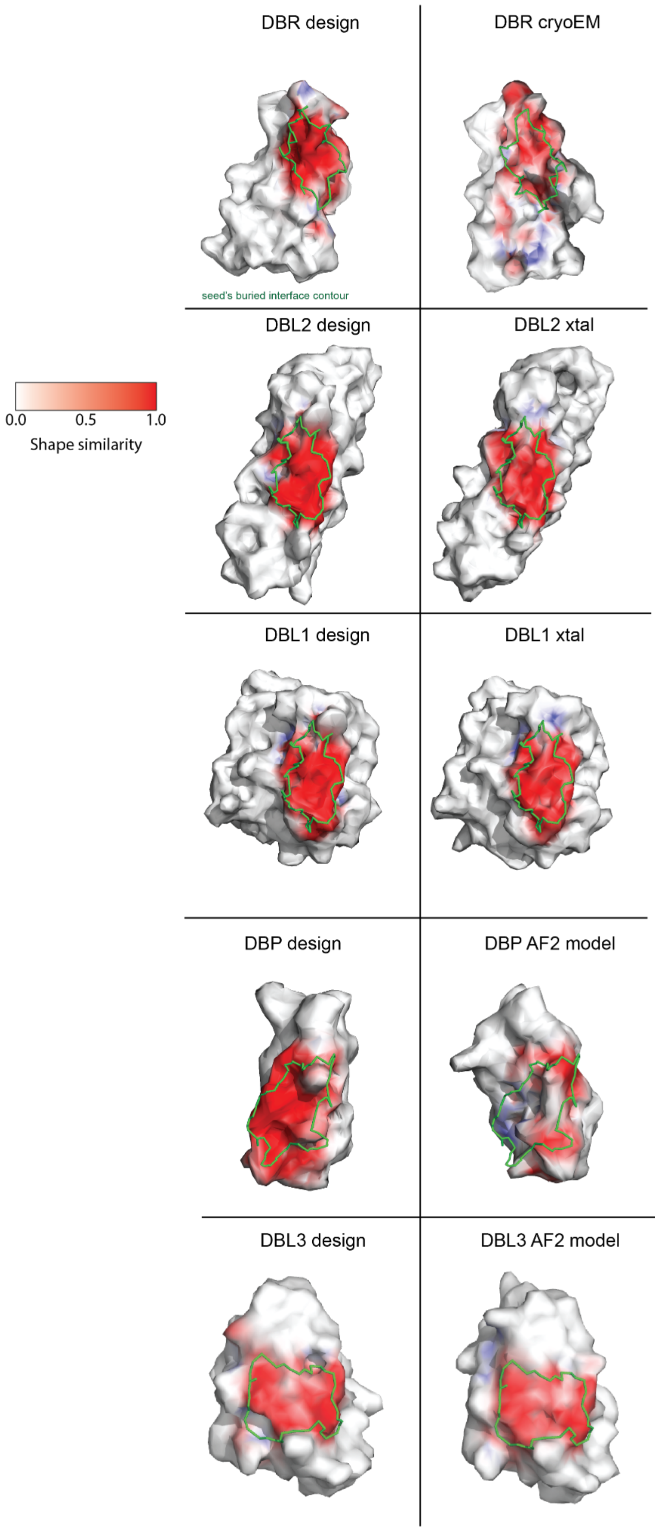

**Supplementary Figure S33: Surface similarity of the computational designs, experimentally solved structures or AF models relative to initial binding seeds.** Each complex was aligned to the target protein (RBD, PD-L1 and PD-1), and the surface similarity of the computational design, the experimental structure or AF model to the binding seed is shown in a gradient from white to red. The buried surface area of the initial binding seed is shown by a green contour. The surface similarity was calculated in the same way as shape complementarity but normal vectors are not inverted during the process, i.e. the normal vectors for both surfaces point outwards of the molecular surface. Briefly, pairs of nearest vertices between the surface of the design or structure/model and the initial binding seed were computed based on the nearest neighbor of the aligned model. The shape similarity was evaluated by computing the dot product of the vertex pairs normal vectors yielding the enclosed angle and scaling it with the distance of the vertex pair. The resulting values are colored in a gradient from white to red and range from 0 (colored in white) indicating no similarity, to 1 (colored in red) indicating high similarity.

**Supplementary Table 1:** *Extended Benchmark of MaSIF-seed against other docking methods in recovering the native binder in the correct conformation from co-crystal structures for 31 helix-receptor complexes or 83 non-helix seed-receptor complexes, discriminating between 1000 decoys.* [a]*Benchmarked method.* [b-d]*Number of receptors for which the method recovered the native binding motif (<3 Å iRMSD) within the* [b]*top 1,* [c]*top 10, and* [d]*top 100 results.* [e]*Number of receptors for which the method did not recover the native binding motif in the top 100 results.* [f]*Average running time in minutes, excluding pre-computation time.*

| | **Method[a]** | **# in top 1[b]** | **# in top 10[c]** | **# in top 100[d]** | **>100[e]** | **Avg time (m)[f]** |
|---|---|---|---|---|---|---|
| **Helical seeds** | MaSIF-seed | 18 | 18 | 20 | 11 | 15 |
| | PatchDock+MaSIF-site | 3 | 5 | 11 | 20 | 86 |
| | ZDOCK | 3 | 4 | 8 | 23 | 2715 |
| | ZDOCK+ MaSIF-site | 1 | 6 | 10 | 21 | 2485 |
| | ZDOCK+ ZRank2 | 6 | 12 | 21 | 10 | 2946 |
| | ZDOCK+ ZRank2+ MaSIF-site | 5 | 11 | 19 | 12 | 2710 |
| **Non-helical seeds** | MaSIF-seed | 41 | 47 | 49 | 34 | 118 |
| | ZDock | 7 | 9 | 22 | 61 | 2206 |
| | ZDock+ ZRank2 | 21 | 33 | 45 | 38 | 2400 |

**Supplementary Table 2:** Sequences of the designed proteins.

| Design | Sequence | # of mutations from WT | # of mutations from design_01 | Mutations |
|---|---|---|---|---|
| DBL1 native scaffold (PDB ID: 3S0D) | MTIEELKTRLHTEQSVCKTETGI DQQKANDVIEGNIDVEDKKV QLYCECILKNFNILDKNNVFKP QGIKAVMELLIDENSVKQLVS DCSTISEENPHLKASKLVQCVS KYKTMKSVDFL | | | |
| DBL1_01 | TSIESLKWTLIVEQILCQLDTGI DQQKANDVIEGNIDVEDKKV QLYCECILKAFHILDKNNVFKP QGIKAVMELLIDENSVKQLVS DCSTISEENPHLKASKLVQCVS KYKTMKSVDFL | 14 | | M1T, T2S, E5S, T8W, R9T, H11I, T12V, S15I, V16L, K18Q, T19L, E20D, N53A, N55H |
| DBL1_02 | SSIESLKWSLIVQQILCQLETGI DQQKANDVIEGNIDVEDKKV QLYCECILKQFHILDKNNVFKP QGIKAVMELLIDENSVKQLVS DCSTISEENPHLKASKLVQCVS KYKTMKSVDFL | 14 | 5 | T1S, T9S, E13Q, D20E, A53Q |
| DBL1_03 | SSIESMKWSMIVQQILCQLET GIDQQKANDVIEGNIDVEDKK VQLYCECILKQFHILDKNNVFK PQGIKAVMELLIDENSVKQLVS DCSTISEENPHLKASKLMQCIS KYKTWKSFDFL | 20 | 11 | L6M, L10M, V104M, V107I, M113W, V116F |
| DBL1_04 | SSIESMKWSMIRQQILCQLET GIDQQKANDVIEGNIDVEDKK VQLYCECILKQFHILDKNNVFK PQGIKAVMELLIDENSVKQLVS DCSTISEENPHLKASKLMQCIS KYKTWKSFDFL | 21 | 14 | E4T, W8N, Q18R |
| DBL2 native scaffold (PDB ID: 3ONJ) | SLLISYESDFKTTLEQAKASLAE APSQPLSQRNTTLKHVEQQQ DELFDLLDQMDVEVNNSIGDA SERATYKAKLREWKKTIQSDIK RPLQSLVDSGD | | | |
| DBL2_01 | SLLESYEWSFIVQLILAKLELAY APSQPLSQRNEQLKRVEQQQ DQLFDLLDQMDVEVNNSIGD ASERATYKAKLREWKKTIQSDI KRPLQSLVDSGD | 15 | | I4E, S8W, D9S, K11I, T12V, T13Q, E15I, Q16L, A19L, S20E, E23Y, T34E, T35Q, H38R, E45Q |

| | | | | |
|---|---|---|---|---|
| DBL2_02 | NLLESYEWSFKVQLILAKLELAKAPSQPLSQRNEELKRVEQRQDRLFDLLDQMDVEVNNSIGDASERATYKAKLREWKKTIQSDIKRPLQSLVDSGD | 16 | 6 | S1N, I11K, Y23K, Q35E, Q42R, Q45R |
| DBL2_03 | NLLTSYEGSFKIQLILAKLELAKAPSQPLSQRNEELKRVEQRQDRLFDLLDQMDVEVNNSIGDASERATYKAKLREWKKTIQSDIKRPLQSLVDSGD | 16 | 9 | E4T, W8G, V12I |
| DBL2_04 | NLLRSYENSFKIQLILAKLELAHAPSQPLSQRNEELKRVEQRQDRLFDLLDQMDVEVNNSIGDASERATYKAKLREWKKTIQSDIKRPLQSLVDSGD | 16 | 9 | T4R, G8N, K23H |
| DBR_01 | STNMLEALQQRLHKYAAVVSRAALENNSGKARRFGRIVKQYEDAIKLYKAGKPVPYDELPVPPGFG | 8 | | E13H, Q16A, S17A, E19V, A20S, A21R, K23A, A24L |
| DBR_02 | STNMLEALQQRLQFYFGVVSRAALENNSGKARRFGRIVKQYEDAIKLYKAGKPVPYDELPVPPGFG | 9 | 4 | H13Q, K14F, A16F, A17G |
| DBR_03 | STNMLEALQQRLQFYHGQVARAALENNSGKARRFGRIVKQYEDAIKLYKAGKPVPYDELPVPPGFG | 9 | 6 | F16H, V18Q, S20A |
| DBR_03_KO | STNMLEALQQRLQFYHRQVRRAALENNSGKARRFGRIVKQYEDAIKLYKAGKPVPYDELPVPPGFG | 9 | | G17R, A20R |
| DBP13_01 | TCEVRCENGNRIEYPATSDLECLHWCLDAIMSHPNYRCTCTHK | 10 | | Q10N, E20L, E23L, R24H, R27L, K28D, K30I, K31M, E32S, F33H |
| DBP13_01 (native scaffold) | TCEVRCENGQRIEYPATSDEECERWCRKAKKEFPNYRCTCTHK | | | |
| DBP40_01 | SQVTWNGVTVTNDNPSQSAMWADLIALLYQGEVRVKDGRWEIH | 12 | | I1S, F12N, E16S, E17Q, A18S, E19A, K20M, Y21W, K23D, K24L, K27L, E28L |
| DBP40_01 (native scaffold) | IQVTWNGVTVTFDNPEEAEKYAKKIAKEYQGEVRVKDGRWEIH | | | |
| DBP52_01 | QKETRHCSGRSCDWWATLW | 13 | | Q10R, R11S, E13D, |

| | | | | |
|---|---|---|---|---|
| | CLLCAMKGKRVRCRQHGQQVEVQCDK | | | Q14W, E15W, R17T, R18L, E21L, E22L, K24A, K25M, K34Q, N37Q |
| DBP52_01 (native scaffold) | QKETRHCSGQRCEQEARRWCEECKKKGKRVRCRKHGNQVEVQCDK | | | |
| DBL3_01 | AQTILLRLEGMDCTSCASSIERAIAKVPGVQQCIVFFEMNLAIVTYHGETTPQILTDAVERAGYHARVS | 11 | | N5L, Q7R, S32Q, Q34I, N36F, A38E, L39M, E40N, Q41L, V43I, S45T |
| DBL3_02 | AQTILLRLEGMDSTSSASSIERAIAKVPGVQQCIVFFEMNLAIVTYHGETTPQILTDAVERAGYHARVS | 13 | 2 | N5L, Q7R, C13S, C16S, S32Q, Q34I, N36F, A38E, L39M, E40N, Q41L, V43I, S45T |
| DBL3_01 (Native scaffold) | AQTINLQLEGMDCTSCASSIERAIAKVPGVQSCQVNFALEQAVVSYHGETTPQILTDAVERAGYHARVL | | | |
| DBL4_01 | PDRWLLRIEIPADIAANEALKVRLLETEGVKIVFINESSHAALVIIDSKVTNRFEVEQAIRQ | 12 | | Y2D, V3R, S4W, S5L, E32I, L34F, A36N, E38S, E39S, S41A, Y43L, K45I |
| DBL4_01 (Native scaffold) | PYVSSLRIEIPADIAANEALKVRLLETEGVKEVLIAEEEHSAYVKIDSKVTNRFEVEQAIRQA | | | |
| DBC2_01 | AFITIMDGEEKARKYAKMLKKQNLKVIVLMANGKWIIYAK | 11 | | K1A, T3I, T5I, E25K, H27I, R29L, V30M, E31A, V36I, T38Y, E40K |
| DBC2_01 (Native scaffold) | KFTTTMDGEEKARKYAKMLKKQNLEVHVRVENGKWVITAE | | | |

**Supplementary Table 3:** SARS-CoV-2 variant mutations.

| Variant (graph label in bold) | Mutations | $EC_{50}$ of DBR3_03 with variant: |
|---|---|---|
| D614G / **WT** | D614G | 6.61e-8 g/mL |
| B.1.1.7 / **Alpha** | Δ69-70, Δ144, N501Y, A570D, D614G, P681H, T716I, S982A, D1118H | 6.56e-8 g/mL |
| B.1.351 / **Beta** | L18F, D80A, D215G, Δ242-244, R246I, K417N, E484K, N501Y, D614G, A701V | 6.11e-7 g/mL |
| B.11.28.1 / **Gamma** | L18F, T20N, P26S, D138Y, R190S, K417T, E484K, N501Y, D614G, H655Y, T1027I, V1176F | 6.76e-7 g/mL |
| B.1.526 / **Iota** | L5F, T95I, D253G, E484K, D614G, A701V | 4.13e-7 g/mL |
| B.1.617.1 / **Kappa** | E154K, L452R, E484Q, D614G, P681R, Q1071H | NA |
| B.1.617.2 / **Delta** | T19R, Δ156-157, R158G, L452R, T478K, D614G, P681R, D950N | NA |
| **Lambda** | G75V, T76I, R246N, Δ247-253, L452Q, F490S, D614G, T859N | NA |
| **Omicron BA.1** | A67V, Δ69-70, T95I, G142D, Δ143-145, Δ211, L212I, ins214EPE, G339D, S371L, S373P, S375F, K417N, N440K, G446S, S477N, T478K, E484A, Q493K, G496S, Q498R, N501Y, Y505H, T547K,D614G, H655Y, N679K, P681H, N764K, D796Y, N856K, Q954H, N969K, L981F | 5.68e-8 g/mL |
| **Omicron BA.2** | T19I, Δ24-26, A27S, G142D, V213G G339D, S371F, S373P, S375F, T376A, D405N, R408S, K417N, N440K, S477N, T478K, E484A, Q493R, Q498R, N501Y, Y505H, D614G, H655Y, N679K, P681H, N764K, D796Y, Q954H, N969K | 4.35e-8 g/mL |

**Supplementary Table 4:** Summary of binding candidates obtained after deep sequencing with the optimized design pipeline. Binding seeds were helical (H) or strand (E). Deep sequencing data comprises reads from the non-binding (Neg reads) and binding population (Pos reads). The enrichment score is calculated based on the logarithm of the ratio between positive and negative reads. Computational models of the complexes were predicted by AlphaFold Multimer (AF) and aligned with respect to the target. Binding signals detected on the surface of yeast (at 500 nM ligand) were categorized as negative(-), marginal (-/+), weak (+), moderate (++) or high (+++). Marginal and weak binding signals were not further characterized (Competition assay, knock-out mutants and negative controls).

| Design | Target | Scaffold origin | Scaffold name | Motif | Neg reads | Pos reads | Enrichment | AF RMSD [Å] | Binding | Competition | Knock-out | Neg Ctrl |
|---|---|---|---|---|---|---|---|---|---|---|---|---|
| DBP13_01 | PD-1 | Miniprotein | EEHE_2.1_02 | H | 776 | 277011 | 2,5526 | 3,3 | +++ | OK | OK | OK |
| DBP40_01 | PD-1 | Miniprotein | EEHEE_rd4_0499 | H | 15 | 18596 | 3,0933 | 7,1 | +++ | OK | Not tested | OK |
| DBP48_01 | PD-1 | Miniprotein | EHEE_rd4_0510 | H | 225 | 2354 | 1,0196 | 16,9 | - | N/A | N/A | N/A |
| DBP52_01 | PD-1 | Miniprotein | EHEE_1.7_09 | H | 12 | 934 | 1,8912 | 15,5 | + | Not tested | Not tested | Not tested |
| DBC1_01 | CTLA-4 | Miniprotein | EEHE_2.1_06 | E | 3 | 1107 | 2,567 | 6,2 | - | N/A | N/A | N/A |
| DBC2_01 | CTLA-4 | Miniprotein | EHEE_rd4_0923 | E | 172 | 39697 | 2,3632 | 34,6 | ++ | OK | OK | OK |
| DBC3_01 | CTLA-4 | Miniprotein | EHEE_rd4_0042 | E | 31 | 3797 | 2,0881 | 30,4 | -/+ | N/A | N/A | N/A |
| DBC4_01 | CTLA-4 | Miniprotein | EHEE_rd4_0924 | E | 55 | 4018 | 1,8636 | 34,9 | -/+ | N/A | N/A | N/A |
| DBC5_01 | CTLA-4 | Miniprotein | EHEE_rd4_0448 | E | 35 | 1203 | 1,5362 | 24,3 | - | N/A | N/A | N/A |
| DBC6_01 | CTLA-4 | Miniprotein | EHEE_rd4_0357 | E | 109 | 1281 | 1,0701 | 22,4 | - | N/A | N/A | N/A |
| DBC7_01 | CTLA-4 | Miniprotein | EHEE_rd4_0924 | E | 114 | 1333 | 1,0679 | 38,4 | - | N/A | N/A | N/A |
| DBC8_01 | CTLA-4 | Miniprotein | EHEE_rd4_0636 | E | 3 | 2162 | 2,8577 | 33,6 | - | N/A | N/A | N/A |
| DBC9_01 | CTLA-4 | Miniprotein | EHEE_rd4_0811 | E | 44 | 1166 | 1,4232 | 32,2 | - | N/A | N/A | N/A |
| DBL3_01 | PD-L1 | PDB | 4A48 (B) | E | 654 | 44391 | 1,8317 | 1,2 | ++ | OK | OK | OK |
| DBL4_01 | PD-L1 | PDB | 4Q2M (A) | E | 306 | 10238 | 1,5245 | 9,1 | ++ | OK | OK | OK |
| DBL5_01 | PD-L1 | Miniprotein | EHEE_rd4_0017 | E | 340 | 5443 | 1,2044 | 15,6 | - | N/A | N/A | N/A |

**Supplementary Table 5:** Antibodies used in flow cytometry experiments.

| Antibody | Catalog number | Supplier | Dilution |
|---|---|---|---|
| Anti-HA, FITC | A190-138F | Bethyl | 1:100 |
| Anti-V5 mouse | MA5-15253 | Invitrogen | 1:333 |
| Anti-mouse, FITC | F0257 | Sigma | 1:100 |
| Anti-His, PE | 130-120-787 | Miltenyi Biotec | 1:50 |
| Anti-Myc, FITC | SAB4700448 | Sigma | 1:100 |
| Anti-human IgG, PE | 12-4998-82 | Invitrogen | 1:100 |

**Supplementary Table 6:** Primer sequences.

| Library: | Primer name: | Primer sequence: |
|---|---|---|
| DBR_01 | 5vny_rev_1 | CAGACGTTGCTGTAGGGCCTCAAGCATGTTCGTGCTGCTAGCAGCGTAGTCTGGAACG |
| DBR_01 | 5vny_fwd_2a | GAGGCCCTACAGCAACGTCTGCWMARATACKYCRBRGTABNARSCNNSGCGGSACTTGAGAATAATAGTGGAAAAGCAAGAAGATTTGGCAGGATC |
| DBR_01 | 5vny_fwd_2b | GAGGCCCTACAGCAACGTCTGCWMYWCTACKYCRBRGTABNARSCNNSGCGGSACTTGAGAATAATAGTGGAAAAGCAAGAAGATTTGGCAGGATC |
| DBR_01 | 5vny_rev_3 | ACAGGTTTTCCAGCTTTATACAACTTAATTGCGTCCTCGTATTGTTTAACGATCCTGCCAAATCTTCTTGCTTT |
| DBR_01 | 5vny_fwd_4 | ATTAAGTTGTATAAAGCTGGAAAACCTGTACCATACGACGAACTACCTGTCCCGCCAGGATTCGGCGGATCCCAGGAACTGACAACTATATG |
| DBL1_L1 | 3S0D_fw1 | GCCTTAGCTCAACCGGTTATTTCTACTACCGTCGGTTCCGCTGCAGAAGGCTCTTTGGACAAGAG |
| DBL1_L1 | 3S0D_rev1 | GCTAGCAGCGTAGTCTGGAACGTCGTATGGGTAAGCTTCTCTCTTGTCCAAAGAGCCTTCT |
| DBL1_L1 | 3S0D_fw3a | CCAGACTACGCTGCTAGCHHCTMCATTSWAAGTTTGAAGTGGAVCTTAWTCRTASAACAAATTVTATGTCAACTTBWCACGGGGATTGACCAGCA |
| DBL1_L1 | 3S0D_fw3b | CCAGACTACGCTGCTAGCHHCTMCATTSWAAGTTTGAAGTGGAVCTTAWTCRTASAACAAATTVTATGTCAACTTGAAACGGGGATTGACCAGCA |
| DBL1_L1 | 3S0D_fw3c | CCAGACTACGCTGCTAGCHHCTMCATTSWAAGTTTGAAGTGGAVCTTAWTCRTATGGCAAATTVTATGTCAACTTBWCACGGGGATTGACCAGCA |
| DBL1_L1 | 3S0D_fw3d | CCAGACTACGCTGCTAGCHHCTMCATTSWAAGTTTGAAGTGGAVCTTAWTCRTATGGCAAATTVTATGTCAACTTGAAACGGGGATTGACCAGCA |
| DBL1_L1 | 3S0D_fw3e | CAGACTACGCTGCTAGCHHCTMCATTSWAAGTTTGAAGTGGAVCTTAWTCRTAWWCCAAATTVTATGTCAACTTBWCACGGGGATTGACCAGCA |
| DBL1_L1 | 3S0D_fw3f | CCAGACTACGCTGCTAGCHHCTMCATTSWAAGTTTGAAGTGGAVCTTAWTCRTAWWCCAAATTVTATGTCAACTTGAAACGGGGATTGACCAGCA |
| DBL1_L1 | 3S0D_rev4 | CATTCGCAATATAGTTGGACTTTTTTGTCCTCCACGTCAATGTTCCCCTCAATCACGTCATTCGCCTTCTGCTGGTCAATCCCCGT |
| DBL1_L1 | 3S0D_fw5a | GTCCAACTATATTGCGAATGTATACTAAAASMATTCTGGATACTTGATARAAATAATGTTTTTAAGCCCCAGGGAATTAAAGC |
| DBL1_L1 | 3S0D_fw5b | GTCCAACTATATTGCGAATGTATACTAAAASMATTCYWCATACTTGATARAAATAATGTTTTTAAGCCCCAGGGAATTAAAGC |

| | | |
|---|---|---|
| DBL1_L1 | 3S0D_rev6 | GATATAGTGCTACAGTCGGAGACAAGCTGTTTAACGCTATTTTCATCTATTAACAGTTCCATCACAGCTTTAATTCCCTGGGGCTT |
| DBL1_L1 | 3S0D_fw7 | CTCCGACTGTAGCACTATATCAGAAGAGAACCCACATCTTAAGGCCAGTAAACTGGTTCAGTGCGTGAGTAAATACAAAACCATGAAAAGCGTGG |
| DBL1_L1 | 3S0D_rev8 | GAGTACGGCGTCGATTCTAAAGTTGGTGAGGGGATTTGCTCGCATATAGTTGTCAGTTCCTGGGATCCCAAGAAGTCCACGCTTTTCATGGTTTTG |
| DBL1_L2 | 3S0D_core_fw3a | CCAGACTACGCTGCTAGCTCCTCCATTGAAAGTVTGAAGTGGAGCMTGATCGTACAACAAATTCTATGTCAACT |
| DBL1_L2 | 3S0D_core_fw3b | CCAGACTACGCTGCTAGCTCCTCCATTGAAAGTVTGAAGTGGAGCTWCATCGTACAACAAATTCTATGTCAACT |
| DBL1_L2 | 3S0D_core_rev4 | CGTCAATGTTCCCCTCAATCACGTCATTCGCCTTCTGCTGGTCAATCCCCGTTTCAAGTTGACATAGAATTTGTTGTACGAT |
| DBL1_L2 | 3S0D_core_fw5 | TGATTGAGGGGAACATTGACGTGGAGGACAAAAAAGTCCAAC |
| DBL1_L2 | 3S0D_core_rev6a | GGCTTAAAAACATTATTTTTATCAAGTATGTGCCATTGTTTGWRCCAACATTCGCAATATAGTTGGACTT |
| DBL1_L2 | 3S0D_core_rev6b | GGCTTAAAAACATTATTTTTATCAAGTATGTGGWRTTGTTTGWRCCAACATTCGCAATATAGTTGGACTT |
| DBL1_L2 | 3S0D_core_rev6c | GGCTTAAAAACATTATTTTTATCAAGTATGTGCCATTGTTTGWRGWWACATTCGCAATATAGTTGGACTT |
| DBL1_L2 | 3S0D_core_rev6d | GGCTTAAAAACATTATTTTTATCAAGTATGTGGWRTTGTTTGWRGWWACATTCGCAATATAGTTGGACTT |
| DBL1_L2 | 3S0D_core_fw7a | CACATACTTGATAAAAATAATGTTTTTAAGCCCCAGGGAATTAAAGCTRTGATGGAACTGACTATAGATGAAAATAGCGTTAAACAGCTT |
| DBL1_L2 | 3S0D_core_fw7b | CACATACTTGATAAAAATAATGTTTTTAAGCCCCAGGGAATTAAAGCTRTGATGGAACTGMTAATAGATGAAAATAGCGTTAAACAGCTT |
| DBL1_L2 | 3S0D_core_rev8 | CAGTTTACTGGCCTTAAGATGTGGGTTCTCTTCTGATATAGTGCTACAGTCGGAGACAAGCTGTTTAACGCTATTTTCATC |

| | | |
|---|---|---|
| DBL1_L2 | 3S0D_core_fw9a | CACATCTTAAGGCCAGTAAACTGRYGCAGTGCVTRTMCAAGTACAAGACCTWCAAAAGCKKGGATTTCCTTGGATCCCAGGA |
| DBL1_L2 | 3S0D_core_fw9b | CACATCTTAAGGCCAGTAAACTGRYGCAGTGCVTRTMCAAGTACAAGACCTWCAAAAGCTWCGATTTCCTTGGATCCCAGGA |
| DBL1_L2 | 3S0D_core_fw9c | CACATCTTAAGGCCAGTAAACTGRYGCAGTGCVTRTMCAAGTACAAGACCWKGAAAAGCTWCGATTTCCTTGGATCCCAGGA |
| DBL1_L2 | 3S0D_core_rev10 | GAGTACGGCGTCGATTCTAAAGTTGGTGAGGGGATTTGCTCGCATATAGTTGTCAGTTCCTGGGATCCAAGGAAATC |
| DBL2_L1 | 3ONJ_fw1 | GCCTTAGCTCAACCGGTTATTTCTACTACCGTCGGTTCCGCTGCAGAAGGCTCTTTGGACAAGAG |
| DBL2_L1 | 3ONJ_rev2 | GCTAGCAGCGTAGTCTGGAACGTCGTATGGGTAAGCTTCTCTCTTGTCCAAAGAGCCTTCTG |
| DBL2_L1 | 3ONJ_fw3a | CAGACTACGCTGCTAGCWMTCTTSDAGAGAGTTATGAATGGASCTTTRWAGTCCRATTGAWATTGGCTAAGTTGGAMCTGGCCMRGGCGCCATCACAGCC |
| DBL2_L1 | 3ONJ_fw3b | CAGACTACGCTGCTAGCWMTCTTSDAGAGAGTTATGAATGGASCTTTRWAGTCCRATTGAWATTGGCTAAGTTGGAMCTGGCCTATGCGCCATCACAGCC |
| DBL2_L1 | 3ONJ_fw3c | CAGACTACGCTGCTAGCWMTCTTSDAGAGAGTTATGAATWTASCTTTRWAGTCCRATTGAWATTGGCTAAGTTGGAMCTGGCCMRGGCGCCATCACAGCC |
| DBL2_L1 | 3ONJ_fw3d | CAGACTACGCTGCTAGCWMTCTTSDAGAGAGTTATGAATWTASCTTTRWAGTCCRATTGAWATTGGCTAAGTTGGAMCTGGCCTATGCGCCATCACAGCC |
| DBL2_L1 | 3ONJ_rev4a | CATCTGGTCCAGTAAATCGAATAATYGATCTTGACGCTGTTCAACACGTTTAAGKTSCTCATTACGTTGAGACAAAGGCTGTGATGGCGC |
| DBL2_L1 | 3ONJ_rev4b | CATCTGGTCCAGTAAATCGAATAATYGATCTTGCTBCTGTTCAACACGTTTAAGKTSCTCATTACGTTGAGACAAAGGCTGTGATGGCGC |
| DBL2_L1 | 3ONJ_rev4c | CATCTGGTCCAGTAAATCGAATAATYGATCTTGACGCTGTTCAACCTBTTTAAGKTSCTCATTACGTTGAGACAAAGGCTGTGATGGCGC |
| DBL2_L1 | 3ONJ_rev4d | CATCTGGTCCAGTAAATCGAATAATYGATCTTGCTBCTGTTCAACCTBTTTAAGKTSCTCATTACGTTGAGACAAAGGCTGTGATGGCGC |
| DBL2_L1 | 3ONJ_fw5 | TTATTCGATTTACTGGACCAGATGGATGTGGAGGTTAATAACAGCATCGGGGACGCATCAGAACGCGCCACTTATAAAG |
| DBL2_L1 | 3ONJ_rev6 | GCTTGATGTCGGACTGGATCGTTTTTTTCCACTCGCGTAACTTTGCTTTATAAGTGGCGCGTTCT |

| | | |
|---|---|---|
| DBL2_L1 | 3ONJ_fw7 | CCAGTCCGACATCAAGCGCCCGCTTCAGAGTTTGGTTGATAGTGGCGATGGATCCCAGGAACTGACAA |
| DBL2_L1 | 3ONJ_rev8 | GAGTACGGCGTCGATTCTAAAGTTGGTGAGGGGATTTGCTCGCATATAGTTGTCAGTTCCTGGGATCC |

**Supplementary Table 7:** Target protein sequences.

| Protein target: | Sequence: | Notes: |
|---|---|---|
| PD-1 | LDSPDRPWNPPTFSPALLVVTEG<br>DNATFTCSFSDTSESFVLNWYRM<br>SPSDQTDKLAAFPEDRSQPGQDS<br>RFRVTQLPNGRDFHMSVVRARR<br>NDSGTYLCGAISLAPKAQIKESLRA<br>ELRVTERRAEVPTAHPSPSPRPAG<br>QFQ | Mutated glycosylation site (N->D) and mutated free cysteines (C->S) underlined |
| CTLA4 | KAMHVAQPAVVLASSRGIASFVC<br>EYASPGKATEVRVTVLRQADSQV<br>TEVCAATYMMGNELTFLDDSICT<br>GTSSGNQVNLTIQGLRAMDTGLY<br>ICKVELMYPPPYYLGIGDGTQIYVI<br>DPEPCPDSD | Mutated glycosylation site underlined (N->D) |
| RBD (WT) | RVQPTESIVRFPNITNLCPFGEVF<br>NATRFASVYAWNRKRISNCVADY<br>SVLYNSASFSTFKCYGVSPTKLND<br>LCFTNVYADSFVIRGDEVRQIAPG<br>QTGKIADYNYKLPDDFTGCVIAW<br>NSNNLDSKVGGNYNYLYRLFRKS<br>NLKPFERDISTEIYQAGSTPCNGV<br>EGFNCYFPLQSYGFQPTNGVGYQ<br>PYRVVVLSFELLHAPATVCGPKKS<br>TNLVKNKCVNFNFNGLTGTGVLT<br>ESNKKFLPFQQFGRDIADTTDAV<br>RDPQTLEILDITPCS | |
| PD-L1 | SFTVTVPKDLYVVEYGSNMTIECK<br>FPVEKQLDLAALIVYWEMEDKNII<br>QFVHGEEDLKVQHSSYRQRARLL<br>KDQLSLGNAALQITDVKLQDAGV<br>YRCMISYGGADYKRITVKVNAPY<br>NKINQRILVVDPVTSEHELTCQAE<br>GYPKAEVIWTSSDHQVLSGKTTT<br>TNSKREEKLFNVTSTLRINTTTNEI<br>FYCTFRRLDPEENHTAELVIPELPL<br>AHPPNERTD | |

**Supplementary Table 8:** Crystallographic data collection and refinement statistics.

| | DBL1_03-PD-L1 | DBL2_02-PD-L1 |
|---|---|---|
| **Data collection** | | |
| Space group | $P\ 4_2\ 2_1\ 2$ | $P\ 2_1\ 2_1\ 2_1$ |
| Cell dimensions | | |
| *a*, *b*, *c* (Å) | 97.93, 97.93, 106.11 | 85.41, 116.08, 149.61 |
| a, b, g(°) | 90.00, 90.00, 90.00 | 90.00, 90.00, 90.00 |
| Wavelength (Å) | 0.97889 | 0.97918 |
| Resolution (Å) | 48.97 - 3.85 (2.95 - 2.85) | 41.06 - 2.85 (3.05 - 2.95) |
| Unique reflections | 12591 (1241) | 31885 (3159) |
| $R_{merge}$ | 0.141 (3.126) | 0.172 (3.326) |
| *I* / s*I* | 20.7 (1.1) | 12.2 (0.9) |
| CC1/2 | 0.998 (0.554) | 0.999 (0.368) |
| Completeness (%) | 99.9 (100.0) | 99.3 (99.3) |
| Redundancy | 25.4 (26.9) | 13.0 (12.8) |
| **Refinement** | | |
| Resolution (Å) | 48.97 - 2.85 | 41.05 - 2.95 |
| No. reflections | 12583 | 31812 |
| $R_{work}$ / $R_{free}$ | 0.3140/0.3366 | 0.2787/0.3026 |

| | | |
|---|---|---|
| No. atoms | | |
| Protein | 2598 | 9611 |
| Ligand/ion | 0 | 0 |
| Water | 21 | 0 |
| *B*-factors | | |
| Protein | 126.1 | 130.9 |
| Ligand/ion | - | - |
| Water | 101.5 | - |
| R.m.s. deviations | | |
| Bond lengths (Å) | 0.004 | 0.002 |
| Bond angles (°) | 0.750 | 0.480 |
| Ramachandran plot | | |
| Favored (%) | 94.30 | 95.86 |
| Allowed (%) | 5.70 | 4.14 |
| Outliers (%) | 0.00 | 0.00 |

*Values in parentheses are for highest-resolution shell.

**Supplementary Table 9:** Cryo-EM data collection and model validation statistics.

| Data collection and processing | D614G-binder | Omicron-binder full | Omicron-binder local |
|---|---|---|---|
| Microscope | TFS Titan Krios G4 + E-CFEG | TFS Titan Krios G4 + E-CFEG | |

| | | | |
|---|---|---|---|
| Detector | Falcon 4 | Falcon 4 | |
| Magnification (nominal) | 195K | 165K | |
| Pixel size (Å) | 0.40 | 0.726 | |
| Voltage (Kv) | 300 | 300 | |
| Electron exposure (e-/Å²) | 80 | 60 | |
| Dose rate (e-/px/s) | 4.53 | 5.4 | |
| Exposure times (seconds) | 2.82 | 5.85 | |
| Defocus range (um) | 0.8-2.0 | 0.8-2.5 | |
| Micrographs | 20 794 | 22 266 | |
| Initial particle images (No.) | 832 816 | 1 820 333 | |
| Final particle images (No.) | 67 432 | 50 758 | |
| Map resolution (Å) | 2.63 | 2.80 | 3.29 |
| FSC threshold (cutoff) | 0.143 | 0.143 | 0.143 |
| Symmetry | C1 | C1 | C1 |
| Refinement | | | |
| Initial model used | 7BNO | 7QO7 | --- |
| Map sharpening B factor (Å²) | -33.7 | -33.8 | 52.7 |
| Model composition | | | |

| | | | |
|---|---|---|---|
| Non Hydrogen atoms<br>Protein residues / Nucleotide<br>Ligands | 25948<br>3236/0<br>NAG:46 | 28058<br>3429/0<br>BMA:12<br>NAG:76 | 2121<br>261/0<br>NAG:2 |
| B factors (Å²) | | | |
| protein<br>Ligand | 2.00/198.38/88.48<br>31.30/175.52/79.67 | 0.11/126.79/59.39<br>28.80/129.22/79.33 | 33.55/111.45/61.63<br>58.45/64.51/61.48 |
| R.m.s.d deviations | | | |
| Bond lengths (Å) | 0.004(0) | 0.002(3) | 0.003(0) |
| bond angles (°) | 0.687(35) | 0.534 (18) | 0.577(0) |
| Validation | | | |
| MolProbity score | 1.73 | 1.85 | 1.96 |
| Clash score | 6.36 | 9.53 | 9.82 |
| Poor rotamers (%) | 0.00 | 0.00 | 0.00 |
| Ramachandran plot | | | |
| Favored (%) | 94.36 | 95.05 | 93.00 |
| Allowed (%) | 5.45 | 4.77 | 7.00 |
| Disallowed (%) | 0.19 | 0.18 | 0.00 |
| PDB | 7ZSS | 7ZRV | 7ZSD |

| | | | |
|---|---|---|---|
| EMDB | 14947 | 14922 | 14930 |
